# “Multiplexed screen identifies a *Pseudomonas aeruginosa*-specific small molecule targeting the outer membrane protein OprH and its interaction with LPS”

**DOI:** 10.1101/2024.03.16.585348

**Authors:** Bradley E. Poulsen, Thulasi Warrier, Sulyman Barkho, Josephine Bagnall, Keith P. Romano, Tiantian White, Xiao Yu, Tomohiko Kawate, Phuong H. Nguyen, Kyra Raines, Kristina Ferrara, Aaron Golas, Michael Fitzgerald, Andras Boeszoermenyi, Virendar Kaushik, Michael Serrano-Wu, Noam Shoresh, Deborah T. Hung

## Abstract

**SUMMARY:** The surge of antimicrobial resistance threatens efficacy of current antibiotics, particularly against *Pseudomonas aeruginosa*, a highly resistant gram-negative pathogen. The asymmetric outer membrane (OM) of *P. aeruginosa* combined with its array of efflux pumps provide a barrier to xenobiotic accumulation, thus making antibiotic discovery challenging. We adapted PROSPECT^1^, a target-based, whole-cell screening strategy, to discover small molecule probes that kill *P. aeruginosa* mutants depleted for essential proteins localized at the OM. We identified BRD1401, a small molecule that has specific activity against a *P. aeruginosa* mutant depleted for the essential lipoprotein, OprL. Genetic and chemical biological studies identified that BRD1401 acts by targeting the OM β-barrel protein OprH to disrupt its interaction with LPS and increase membrane fluidity. Studies with BRD1401 also revealed an interaction between OprL and OprH, directly linking the OM with peptidoglycan. Thus, a whole-cell, multiplexed screen can identify species-specific chemical probes to reveal novel pathogen biology.

## INTRODUCTION

The alarming rise of antibiotic resistance worldwide highlights the urgent need for rapid discovery and development of novel antibiotics. The field of antibiotic discovery, however, has faced significant challenges in the scientific, regulatory, and economic realms.^1,2^ Tackling gram-negative pathogenic species, in particular WHO prioritized pathogens *Pseudomonas aeruginosa*, *Acinetobacter baumannii*, and two *Enterobacteriaceae* species, *Escherichia coli* and *Klebsiella pneumoniae*,^3^ has been particularly challenging because of the inability of small molecules to easily penetrate the bacterial cell wall and accumulate intracellularly to engage the target.^4,5^

The gram-negative cell envelope consists of three layers, an inner cytoplasmic membrane, a relatively thin peptidoglycan (PG) layer and a lipid-laden outer membrane (OM) decorated with lipopolysaccharide (LPS) on the outer leaflet. In addition to the permeability barrier posed by the asymmetric OM, small molecules are actively pumped out by an array of efflux transporters spanning the envelope.^6^ To date, with few exceptions such as β-lactams and glycopeptides, most antibiotics must cross the double membrane of gram-negative bacteria and accumulate within the cytoplasm to engage their respective targets. However, more recent large-scale screening efforts have been unable to effectively identify hits, even in efflux-deficient strains because of these combined challenges.^5^ To circumvent the need for xenobiotic intracellular accumulation, targeting essential proteins localized in the OM or periplasmic space is a potentially attractive strategy.^7^ Examples are murepavadin (POL7080), a first-in-class antibiotic that targets the OM-localized lipopolysaccharide transport protein, LptD, in *P. aeruginosa*,^8^ darobactin, a natural product that inhibits the OM β-barrel protein, BamA,^9^ and zosurabalpin, a macrocyclic peptide that inhibits the inner membrane LPS transporter complex, LptB_2_FGC.^10,11^

Chemical screening for small molecules with antibacterial activity has historically been either target-based (i.e., a purified protein inhibition assay) or whole-cell (i.e., growth inhibition). Combining the two approaches, target-based whole-cell screening is based on the concept of altered sensitization to an inhibitor when dosage of its target is altered, first pioneered in yeast.^12^ This principle has been applied to antibiotic discovery by high throughput screening against genetically engineering strains that under-express or deplete (hypomorphs) essential gene targets, thus hypersensitizing to corresponding inhibitors.^13,14^ Recently, in a strategy called PROSPECT (<u>Pr</u>imary screening <u>o</u>f <u>S</u>trains to <u>P</u>rioritize <u>E</u>xpanded <u>C</u>hemistry and <u>T</u>argets),^15^ we reported a highly multiplexed strategy in *Mycobacterium tuberculosis* that allowed high-throughput chemical screening of a large set of pooled, barcoded hypomorphs, each depleted for a different essential target. In PROSPECT, the census of each hypomorph in each well of a 384-well plate, in response to each different compound, is deconvoluted based on enumerating amplified barcode counts by next-generation sequencing. PROSPECT enables the discovery of greater than 100 times the number of active molecules compared to simply screening wildtype *M. tuberculosis* and provides biological insight into compound mechanisms from primary screening data based on the identity of hypersensitized strains.^15^ PROSPECT, with its early association between chemical structures and biological activities, can thus be applied to the large-scale discovery of new chemical probes to gain insight into essential bacterial functions.

Here we have adapted the PROSPECT strategy to target *P. aeruginosa* but in a more focused, rather than genome-wide manner, in a method termed mini-PROSPECT. We chose to specifically target essential, OM-embedded proteins (OMPs) or OM-associated proteins (OMAPs) in *P. aeruginosa* to enable the discovery of compounds that would circumvent the hurdle of intracellular accumulation and enable the study of functional interactions among these OMPs. In addition to a more focused version of PROSPECT, we made several advances including the development of a new library construction method to dramatically decrease the cost and complexity of screening and a new analytical method for identifying active molecules, both of which enable the method to be more easily and widely applied.

To identify chemical probes to study the biology of *P. aeruginosa* OMPs, we engineered a set of barcoded strains that were depleted for OMPs or OMAPs essential for survival.^16^ We then performed a pooled, multiplexed, high-throughput mini-PROSPECT chemical screen of ∼54,000 compounds, leading to the identification of a novel compound, BRD1401 (Fig 4A) that specifically inhibited a strain depleted of the essential OM lipoprotein, OprL (known as Pal in other gram-negative species). Whole genome sequencing of resistant clones and biochemical studies showed that BRD1401 targets a non-essential OM protein, OprH, to inhibit growth of the *oprL*-hypomorph. OprH, largely absent in all other gram-negative species aside from *Pseudomonas,* is a β-barrel OMP that is known to bind and stabilize LPS, especially when magnesium ions are depleted from the environment.^17–19^ Here we show that BRD1401 disrupts OprH’s interactions with LPS to decrease LPS organization and increase membrane fluidity, impairing survival. We also show a direct genetic and biochemical interaction between OprH and OprL, resulting in an OprH-OprL complex that bridges the OM and PG. Thus, mini-PROSPECT screening led to the identification of a novel *P. aeruginosa*-specific inhibitor, BRD1401, which revealed new biology linking OprL to OprH in the OM.

**Figure 1.**
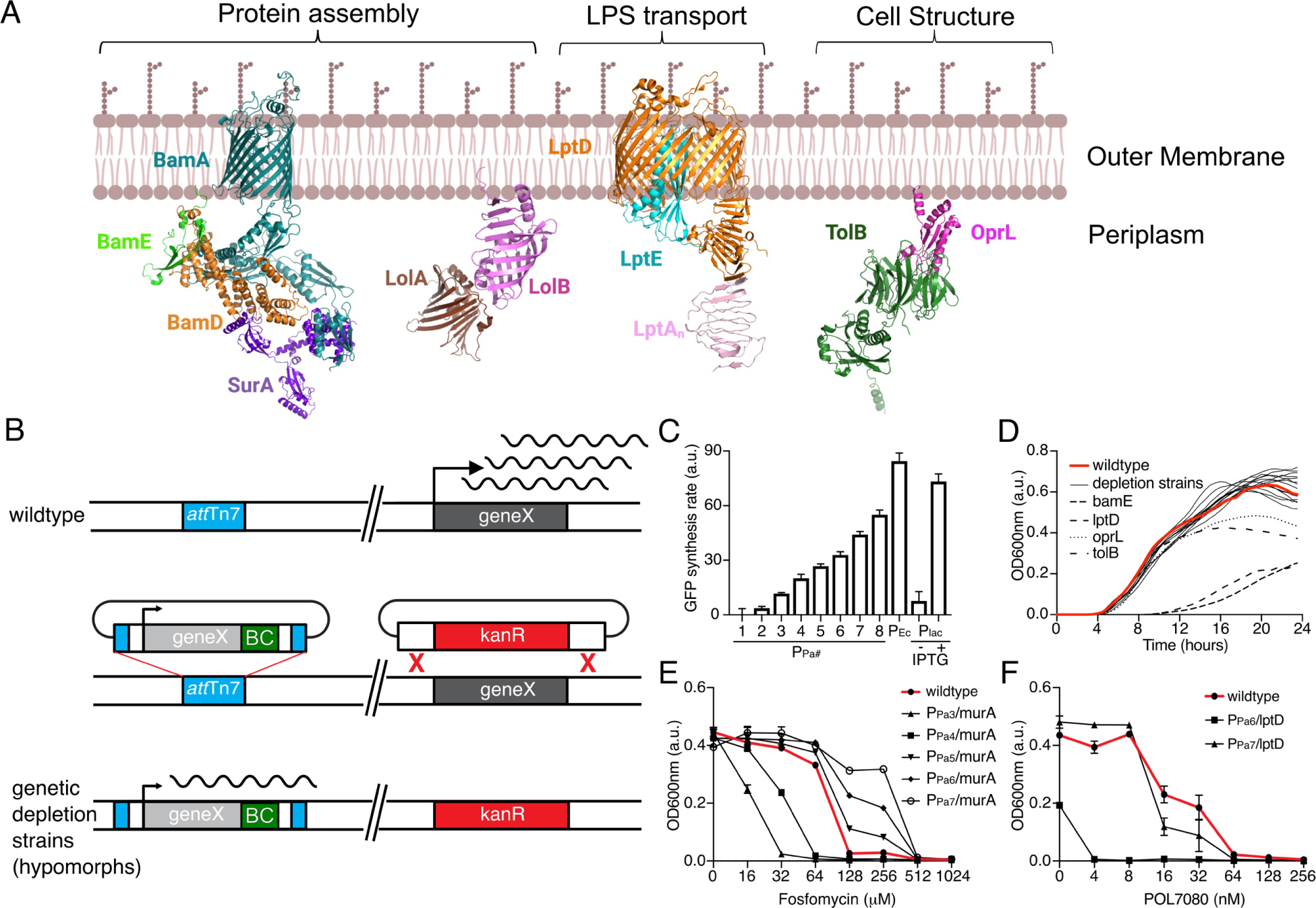
Genetic depletion strains to increase small molecule sensitivity. (A) Schematic of the *P. aeruginosa* essential periplasmic and OM proteins that are the primary targets in this study. Structures of *P. aeruginosa* BamE (7JRK), LolA (2W7Q), LptDE (5IVA) and LptA (4UU4) are from Protein Data Bank, while all other structures are predictions by AlphaFold. (B) Each gene of interest (light grey) driven by one of eight constitutive promoters (P_Pa1-8_) was cloned into the mini-Tn7 suicide delivery system with a unique barcode (BC, green) and integrated at a single conserved *att*Tn7 site in *P. aeruginosa* (blue); the native gene copy (dark grey) was replaced by allelic exchange with a kanamycin resistance gene (red). (C) Quantification of expression strength of promoter panel. The library of eight constitutive promoters (P_Pa1-8_) used for creating depletion strains were each integrated at the *att*Tn7 site driving GFP expression in *P. aeruginosa*, demonstrating varying expression levels as measured by GFP fluorescence. The *E. coli* consensus promoter (P_Ec_) and the IPTG inducible lac promoter (P_lac_) with or without inducer are shown for comparison. Values are subtracted by the wild type PA14 background (n=4; error bars indicate SEM). (D) Growth kinetic of *P. aeruginosa* depletion strains (black solid lines) and the wildtype strain PA14 (red) measured by an absorbance (OD_600nm_)-based assay. All strains except for *bamE*-and *lptD*-hypomorph strains (dashed lines) grew with similar rates. Two depletion strains (*oprL*-and *tolB*-hypomorph strains, dashed lines) grew to a lower, final, optical density. An average of 16 replicates is shown. (E) Impact of MurA expression on fosfomycin activity measured by an absorbance (OD_600nm_)-based growth assay. OD_600nm_ of 5 hypomorph strains constructed using promoters P_Pa3-7_ to drive expression of MurA, the target of fosfomycin, exposed to a dose response of the compound for 16 hours (n=3; error bars indicate SEM). F) Impact of LptD down regulation on activity of its cognate inhibitor, POL7080. OD_600nm_ of engineered *lptD*-hypomorph strains with the P_Pa6_ or P_Pa7_ promoters driving *lptD* expression after exposure to a dose response of POL7080 for 16 hours (n=3; error bars indicate SEM). Wildtype PA14 strain is highlighted in red in (E) and (F).

**Figure 2.**
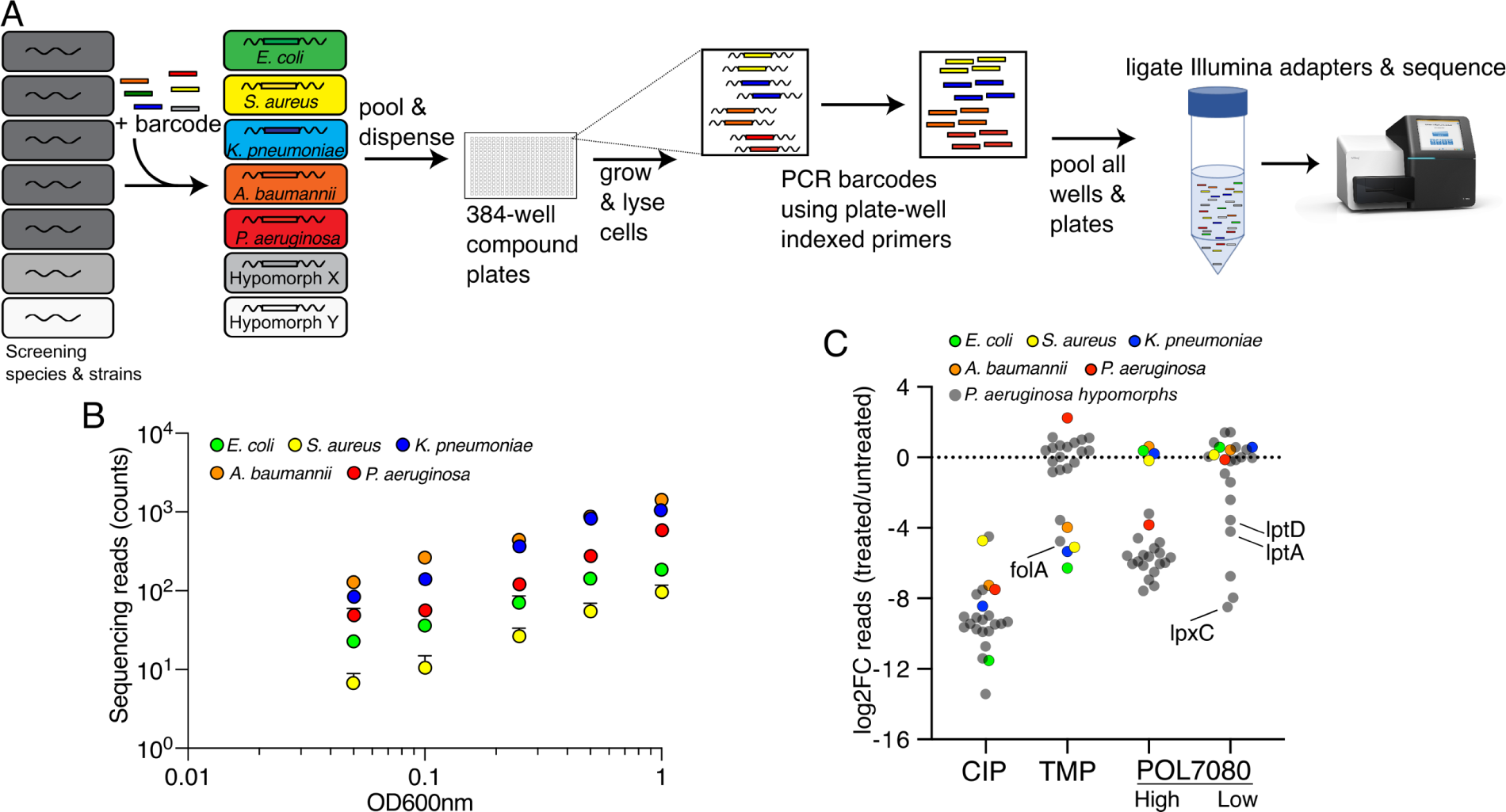
Development of a multiplexed chemical screening assay. (A) Schematic of the multiplexed screening strategy. 76-bp DNA barcode fragments (consisting of a common 26-bp 5’ PCR handle, a unique 24-bp barcode sequence, and a common 26-bp 3’ PCR handle) are chromosomally integrated into each species and hypomorph strain of interest. Strains are individually grown to mid-log phase and pooled before dispensing to 384-well plates +/-compound. After an incubation period in the presence of a chemical library, cells are lysed, and lysate is transferred to 384-well PCR plates containing PCR reagents including a 5’ primer containing a ‘well index’ and a 3’ primer containing a ‘plate index’, to amplify the barcoded region. The PCR products are pooled, ligated with Illumina adaptors, sequenced via Illumina sequencing technology, and plate-well indices are demultiplexed to determine the read count of each strain per well. (B) Mean sequencing read counts of each barcoded species is proportional to OD_600nm_ in an evenly mixed pool of the 5 species (n=12; error bars indicate SD). (C) Growth as reflected in log_2_ fold-change of sequenced barcode reads^15^ (relative to DMSO control) of wildtype bacterial species (colored circles) and *P. aeruginosa* depletion strains (grey circles) in response to known antibacterial compounds relative to DMSO vehicle control grown in multiplex for 12 hours (n = 12). The broad-spectrum antibiotic ciprofloxacin (CIP, 8 μM) inhibits growth for all strains, while trimethroprim (TMP, 32 μM), and POL7080 (High, 64 nM; Low, 8 nM) inhibit only select strains and species. The target of TMP, FolA, and the target of POL7080, LptD, are indicated. At low concentrations of POL7080, other hypomorphs depleted for LPS synthesis/transport related genes such as *lpxC* and *lptA* are also depleted.

**Fig. 3.**
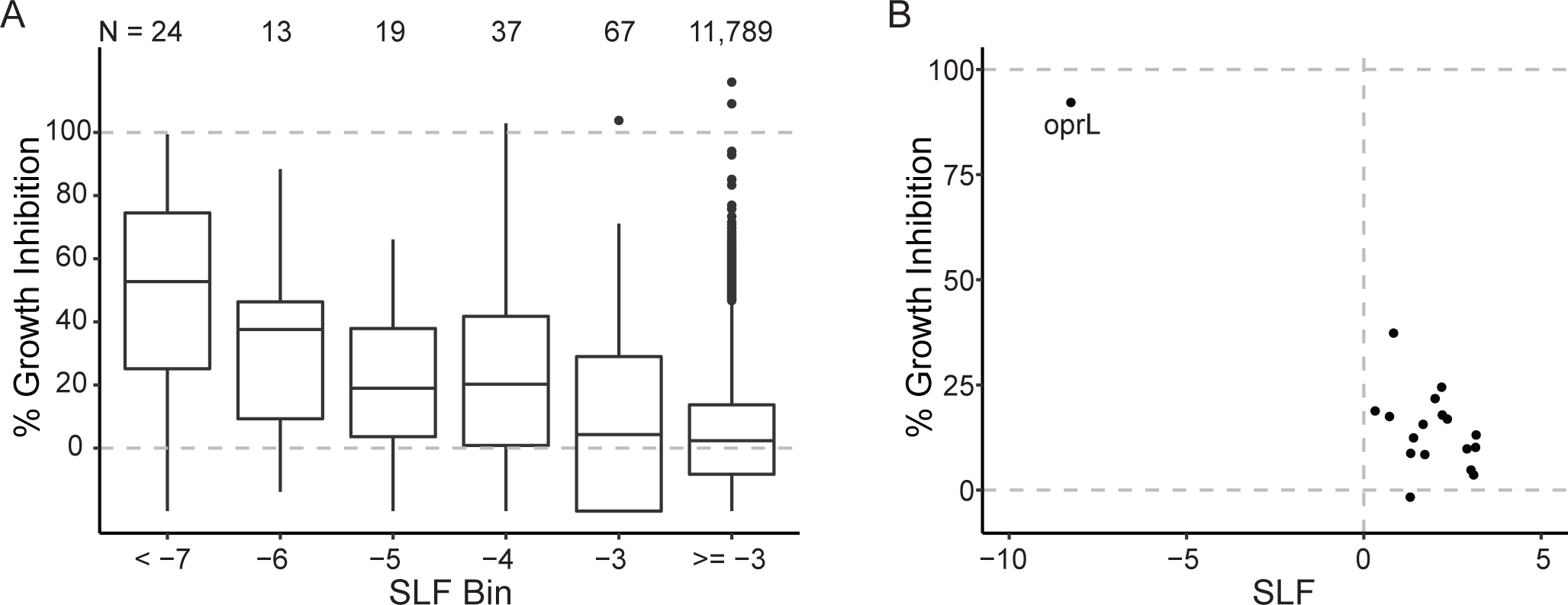
SLF metric from primary multiplexed screen validated as a measure of *P. aeruginosa* strain sensitivity to a given compound. (A) For individual compound-strain interactions tested in the demultiplexed assay, median % growth inhibition relative to growth in vehicle control (DMSO) and positive control (ciprofloxacin), as measured by absorbance (OD_600nm_), is plotted against its corresponding SLF value from the primary multiplexed screen. SLF values are assigned to an SLF bin, with a single bin number representing all SLF values between the listed bin value and the bin value to its left (e.g., bin “-5” indicates −6 ≤ SLF < −5). N indicates the number of compound-strain pairs included in each bin. Increasingly negative SLF values for specific *P. aeruginosa* strains under a particular treatment in the multiplexed screen correspond to increasing growth inhibition when the strain is individually tested against the compound in a demultiplexed OD_600nm_ assay. (B) BRD1401 is an example of a compound that was specifically active against the *P. aeruginosa oprL-*hypomorph strain in the multiplexed screen and whose activity was confirmed in the demultiplexed assay. Median % growth inhibition of BRD1401 in the demultiplexed assay versus its SLF in the multiplexed screen is plotted. Each point represents a different *P. aeruginosa* strain.

**Figure 4.**
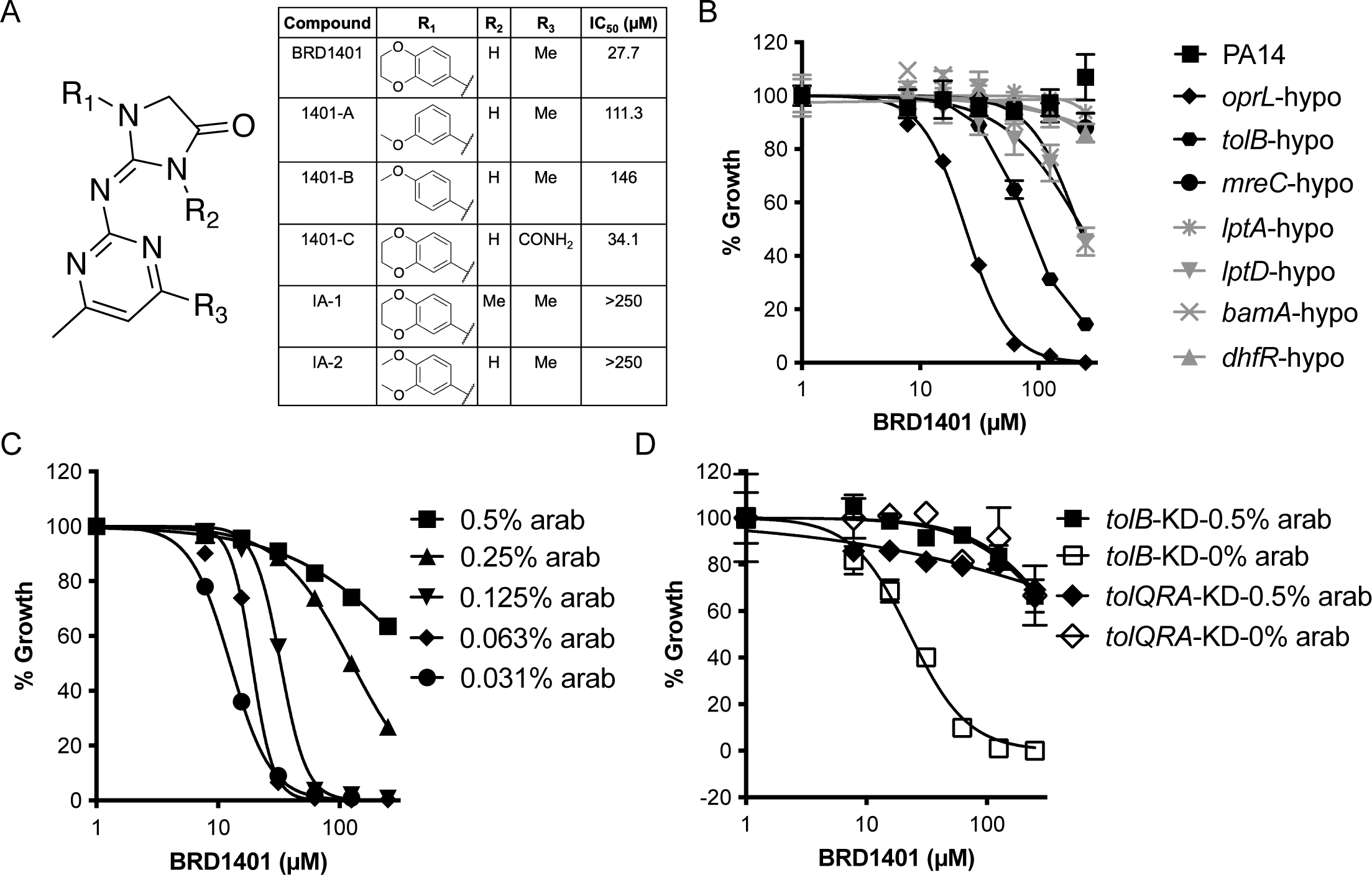
BRD1401 has specific activity towards *P. aeruginosa* when OprL or TolB is depleted. (A) Correctly assigned structure of BRD1401 and 5 synthesized analogs. Growth inhibitory activity against the *oprL*-KD strain, measured as IC_50_ in a demultiplexed, absorbance (OD_600nm_)-based growth assay (Figure S10E), is shown. (B) Activity of BRD1401 towards *P. aeruginosa* wildtype PA14 and a select subset of hypomorph strains in a demultiplexed, absorbance (OD_600nm_)-based growth assay shows specific inhibition of *P. aeruginosa* strains depleted of OprL or TolB. Normalized growth, calculated in % relative to vehicle and positive controls, is plotted against BRD1401 concentration (n= 3, error bars indicate SD). (C) Normalized growth of the *oprL*-KD strain in media with varying doses of arabinose and BRD1401 is plotted against BRD1401 concentration (n= 3, error bars indicate SD). *oprL* expression is under the control of the P_araBAD_ promoter. (D) Normalized growth of the *tolB*-KD and *tolQRA*-KD strains in media with either high (0.5%) or low (0%) arabinose and BRD1401 is plotted against BRD1401concentration (n= 3, error bars indicate SD). *tolB and tolQRA* expression is under the control of the P_araBAD_ promoter in the *tolB*-KD and *tolQRA*-KD strains, respectively. BRD1401 activity is correlated with decreased *oprL* and *tolB* expression but not of *tolQRA* complex.

## RESULTS

### Selecting *P. aeruginosa* targets for mini-PROSPECT chemical screening

We had previously defined a set of 321 core essential genes in *P. aeruginosa* using transposon insertion sequencing (Tn-Seq) and a statistical model, *FiTnEss*, to classify genes as essential versus nonessential in nine *P. aeruginosa* strains across five different media conditions.^16^ Here, we analyzed the subcellular localization of the proteins encoded by the 321 core essential genes using the prediction algorithm PSORTb V3.0, followed by literature confirmation, and identified 7 OMP/OMAPs and 4 periplasmic proteins, which are part of 4 complexes with the following functions (Figure 1A): β-barrel protein folding and assembly (BamADE and SurA),^20^ OM lipoprotein folding and localization (LolAB),^21^ lipopolysaccharide transport (LptADE),^22^ and structural integrity and cell division (TolB and OprL, also known as Pal).^23^ These proteins were selected for hypomorph strain construction and subsequent mini-PROSPECT chemical screening (Table 1). MreC, which has recently been shown to self-associate and thus regulate PG synthesis through a periplasmic functional domain, was also included, even though it is part of an inner membrane complex.^24,25^ In addition to these proteins, we included several cytosolic proteins with known small molecule inhibitors as controls. These included: FolA and FolP, which are inhibited by trimethoprim and sulfamethoxazole, respectively, and catalyze folate biosynthesis;^26,27^ MurA, which is inhibited by fosfomycin and is involved in Lipid I and Lipid II synthesis;^28^ GyrA, which is involved in the activity of the fluoroquinolones and is critical for DNA replication;^29^ LeuS, a leucyl-tRNA synthetase that is targeted by GSK2251052/AN3365;^30^ and LpxC, which is targeted by PF-04753299, and is involved in LPS biosynthesis.^31,32^

**Table 1.** Species and strains constructed for the multiplexed screen.

| Species | Strain/Gene | Subcellular Location | Engineered promoter<sup>#</sup> |  | Inhibitor sensitivity |
| --- | --- | --- | --- | --- | --- |
|  |  |  | Lowest | Highest |  |
| *E. coli* | CFT073 |  |  |  |  |
| *S. aureus* | Newman |  |  |  |  |
| *K. pneumoniae* | RB120 |  |  |  |  |
| *A. baumannii* | ATCC19606 |  |  |  |  |
| *P. aeruginosa* | PA14 |  |  |  |  |
| *P. aeruginosa* | *bamA* | Outer membrane | P<sub>Pa3</sub> | P<sub>Pa3</sub> | - |
| *P. aeruginosa* | *bamD* | Outer membrane | - | - | - |
| *P. aeruginosa* | *bamE* | Outer membrane | P<sub>Pa6</sub> | P<sub>Pa8</sub> | - |
| *P. aeruginosa* | *folA* | Cytoplasm | P<sub>Pa5</sub> | P<sub>Pa8</sub> | 4x |
| *P. aeruginosa* | *folP* | Cytoplasm | P<sub>Pa8</sub> | P<sub>Pa8</sub> | 2x |
| *P. aeruginosa* | *gcp* * | Cytoplasm | P<sub>Pa3</sub> | P<sub>Pa8</sub> | - |
| *P. aeruginosa* | *gyrA* | Cytoplasm | P<sub>Pa1</sub> | P<sub>Pa8</sub> | no change |
| *P. aeruginosa* | *leuS* | Cytoplasm | P<sub>Pa2</sub> | P<sub>Pa8</sub> | no change |
| *P. aeruginosa* | *lolA* | Periplasm | P<sub>Pa3</sub> | P<sub>Pa6</sub> | - |
| *P. aeruginosa* | *lolB* | Outer membrane | P<sub>Pa5</sub> | P<sub>Pa7</sub> | - |
| *P. aeruginosa* | *lppL* ** | Outer membrane | P<sub>Pa8</sub> | P<sub>Pa8</sub> | - |
| *P. aeruginosa* | *lptA* | Periplasm | P<sub>Pa1</sub> | P<sub>Pa8</sub> | - |
| *P. aeruginosa* | *lptD* | Outer membrane | P<sub>Pa6</sub> | P<sub>Pa7</sub> | 16x |
| *P. aeruginosa* | *lptE* | Outer membrane | - | - | - |
| *P. aeruginosa* | *lpxC* | Cytoplasm | P<sub>Pa4</sub> | P<sub>Pa4</sub> | 2x |
| *P. aeruginosa* | *mreC* | Periplasm | P<sub>Pa4</sub> | P<sub>Pa8</sub> | - |
| *P. aeruginosa* | *murA* | Cytoplasm | P<sub>Pa3</sub> | P<sub>Pa7</sub> | 4x |
| *P. aeruginosa* | *oprL* | Outer membrane | P<sub>Pa5</sub> | P<sub>Pa5</sub> | - |
| *P. aeruginosa* | *tolB* | Periplasm | P<sub>Pa8</sub> | P<sub>Pa8</sub> | - |
| *P. aeruginosa* | *surA* | Periplasm | - | - | - |

<sup>#</sup>The strain selected for multiplexed screen had the lowest strength promoter
\**gcp* was initially categorized as an OMP, but a more comprehensive analysis of its localization indicated it is a cytoplasmic protein.
\*\**lppL* was called an essential gene by preliminary analysis of Tn-Seq data in the PA14 strain, but further optimization of the analytical pipeline (*FiTnEss*) and analysis of multiple strains re-annotated it as a non-essential gene.

### Constructing and validating barcoded *P. aeruginosa* strains depleted for essential OMP and OMAP targets

To control expression levels of the essential genes of interest, we engineered strains using constitutive rather than inducible promoters to avoid the potential strain-to-strain variability in their dependence on the concentration of a small molecule inducer, which would complicate pooling of strains in a single screen. Meanwhile, because different levels of each essential gene product are needed for survival, we utilized a set of synthetic constitutive promoters, which were based on the *E. coli* rrnB P1 consensus promoter (called P_Ec_ in this study), whose activities span a large dynamic range.^33^ This set involved six promoter variants of P_Ec_, named P_Pa3_ to P_Pa8_ in this study, that enabled tailoring of the expression levels of each individual protein (Table S1A and Method Details).

To characterize this set of promoters in wildtype *P. aeruginosa*, we chromosomally integrated these promoters to drive *gfp* expression in *P. aeruginosa* PA14 strain and measured relative GFP expression using fluorescence (Figure 1C). The GFP levels of strains utilizing the synthetic promoter variants P_Pa3_ to P_Pa8_ ranged from 13.7% (P_Pa3_) to 65.1% (P_Pa8_) of the expression in the P_Ec_ strain. To increase the dynamic range, we created two additional, even weaker promoters, P_Pa1_ and P_Pa2_, by reducing the spacing between the Pribnow box and −35 sequence of P_Pa3_ from 17 bp to 15 bp and 14 bp, respectively. P_Pa1_ and P_Pa2_ resulted in GFP expression of 0.3% and 4% of the levels relative to P_Ec_ and had lower GFP expression than an uninduced *lac* promoter, which itself was ‘leaky’ and expressed GFP at 8.9% of P_Ec_. The set of promoter variants (P_Pa1_ to P_Pa8_) thus had a dynamic range of 0.3-65.1% of P_Ec._

To generate the desired engineered depletion strains for the 20 genes of interest, we used a two-step promoter replacement strategy. First we cloned each of the 8 promoter variants (P_Pa1_ to P_PA8_; Table S1A) in front of a copy of the gene of interest into a mini-Tn7 suicide plasmid and then integrated them into the *att*Tn7 site of PA14.^34^ This was followed by deleting the native copy of the gene by two-step allelic recombination (Fig 1B).^35,36^ Presumably, only strains in which the synthetic promoter provided sufficient expression of the essential gene of interest to confer viability could be obtained. We were able to successfully construct hypomorphic strains corresponding to 17 of the 20 desired genes, failing for *bamD*, *surA*, and *lptE*. Failure to obtain viable strains for these 3 genes after at least three attempts was interpreted as potentially implying that none of the constitutive promoters controlled expression precisely enough to allow deletion of the native copy. Interestingly, for 6 of the 17 successfully obtained genes, only a single strain corresponding to a single promoter strength could be constructed, suggesting that protein levels of these targets may be tightly controlled (Table 1). Growth rates of the hypomorph strain for each gene with the weakest promoter strength (lowest protein expression) that could be constructed were all similar to that of wildtype PA14, except for *bamE*-and *lptD*-hypomorphs which grew considerably slower (Fig. 1D).

While no significant growth defect was observed in most hypomorphs, hypomorphs with a known inhibitor were nevertheless hypersensitized to chemical inhibition, thus confirming a degree of target depletion in these strains (Table 1). For example, of the five *murA*-hypomorph strains that were successfully constructed with different promoters (P_Pa3_ to P_Pa7_), the two strains with the weakest promoters (P_Pa3_ and P_Pa4_), despite their normal growth, were hypersensitized to fosfomycin, a known MurA inhibitor. Target depletion resulted in 2 to 4-fold decreases in IC_90_ relative to wildtype PA14 (Fig. 1E). Similar results were obtained for the *lptD* hypomorph and its known inhibitor POL7080.^8,37^ The P_Pa7_/*lptD* strain that grew normally had a 2-fold decrease in IC_90_ of POL7080 relative to wildtype, while the P_Pa6_/*lptD* strain that had a growth defect had a ∼16-fold decrease in IC_90_ (Figure 1F). The significant growth defect and hypersensitization of the P_Pa6_/*lptD* strain are both consistent with a more pronounced depletion of LptD relative to the P_Pa7_/*lptD* strain, which was also confirmed by mass spectrometry proteomic measurement (Figure S1A). Thus, depletion of an essential target in *P. aeruginosa* led to increased susceptibility to a cognate inhibitor.

During strain construction we also introduced a unique 24 bp DNA barcode (Table S1C) chromosomally, immediately following the essential gene of interest at the attTn7 integration site. This single copy barcode enabled pooling of the strains, with enumeration of strain census within the pool by counting amplified barcodes using next-generation sequencing, as previously described for PROSPECT.^15^ On either side of the unique barcode were conserved restriction sites and flanking regions used for PCR amplification of the barcode. 30 nt long amplification primers consisted of a GC clamp, an 8 bp unique well or plate barcode, and a 20 bp annealing sequence to the barcode flanking regions (Figure S1A and Table S1D). In all, 96 PCR-forward primers (with ‘well barcodes’) and 96 PCR-reverse primers (with ‘plate barcodes’) were synthesized (Table S1D). Previously, in PROSPECT,^15^ to amplify the barcode, we used long 86 bp primers that contained not only unique plate and well indices but also the Illumina adaptors to allow moving directly from amplification to sequencing without additional library construction. This strategy required 384 unique, long HPLC-purified primers to provide well identity, and a similar set to provide plate identity (Table S1E). Here, we modified the library construction strategy to reduce cost by utilizing much shorter primers that contained well or plate indices but not the Illumina adaptors (Table S1D). In a second step, the Illumina adaptors were ligated to the pool of amplicons combined from all wells (Figure S1B). The unique barcode sequences were selected to have minimal overlap with each other.^38^ We also barcoded the parent wildtype PA14 strain and 4 other strains of clinically relevant pathogens, *E. coli* str CFT073, *K. pneumoniae* str RB120, *A. baummanii* str ATCC19606 and *S. aureus* str Newman, enabling rapid assessment of the spectrum of activity of active compounds in the primary screen. In total, there were 22 strains genetically engineered for inclusion in the pool (Table 1).

### Developing a multiplexed chemical screening assay

We used the pool of barcoded, engineered strains in the development of a multiplexed assay for high-throughput chemical screening in 384-well format (Figure 2A). After exposure to compounds, the final census of each mutant in each well was enumerated by lysing surviving bacteria and PCR amplifying barcodes, followed by pooling of all amplicons, ligation of Illumina adaptors for library construction, and next-generation sequencing to quantify barcodes (Method Details). A comparison of the sequencing read counts generated by the new, cheaper barcoding and amplification strategy with the previously reported PROSPECT strategy showed strong correlation between their respective outputs (Figure S1C).

The resulting library was sequenced to obtain a theoretical target depth of 100 reads per strain per well. Read counts of barcodes were demultiplexed and normalized for PCR amplification variance using internal PCR barcode controls spiked into each well. We confirmed that read counts of barcodes directly correlated to bacterial cell numbers of the corresponding strains across a dose response of cell densities (Figure 2B; Method Details). Using this sequencing readout, we optimized the inoculation size and incubation period for a pool in which all strains were inoculated with equal representation (Figure S2A; Method Details). The median Z’ factor of all strains was 0.59, with only 3 strains falling below a threshold of 0.5 (Table S2A).

Testing of known antibiotics on the strain pool compared with a DMSO control confirmed the assay’s ability to detect small molecule inhibitors (Figure 2C). As expected, treatment with the broad-spectrum antibiotic, ciprofloxacin, resulted in >4 log_2_-fold decrease in barcode reads relative to the untreated samples for all strains and species. Treatment with trimethoprim, which is active against all included species except *P. aeruginosa*, resulted in >4 log_2_-fold decrease in barcode reads for *E. coli*, *K. pneumoniae*, *A. baumannii*, and *S. aureus*, while having no effect on all *P. aeruginosa* strains except for the *P. aeruginosa folA*-hypomorph, consistent with FolA being the target of trimethoprim. Treatment with the specific anti-*P. aeruginosa* antibiotic POL7080 at high concentration resulted in significant decrease in reads for all *P. aeruginosa* strains including wildtype, but no other species. However, at lower concentrations of POL7080, the *lptD*-, *lpxC*-, and *lptA*-hypomorphs continued to show a >3 log_2_-fold reduction in reads, even while activity against wildtype PA14 and other *P. aeruginosa* hypomorphs was lost. LptD, which transports LPS across the OM, is the proposed target of POL7080 while LpxC and LptA are involved in LPS biosynthesis and LPS transport across the periplasm, respectively.

### Chemical high-throughput screening for hypomorph inhibitors

Using the multiplexed, pooled assay with 21 strains, we performed a high-throughput screen on a diverse compound library of ∼54,000 small molecules from the Broad Institute’s internal collection [∼31,000 compounds from NIH’s Molecular Libraries Probe Production Centers Network (MLPCN), ∼13,000 diversity-oriented synthetic compounds,^39^ and ∼10,000 compounds from a commercial Wuxi collection at a concentration of 67 μM].

First, to identify compounds that suppress the growth of the entire strain pool, we identified wells in which the total barcode read count was less than 10% of the median count in vehicle control wells from the same plate. This method identified two compounds as potentially pan-active, acting against all represented species. One was tested in an absorbance (OD_600nm_)-based assay, but its activity did not confirm. In addition, to identify compounds that may be active against all represented *P. aeruginosa* strains, even if not active against other species, we performed a similar calculation after subsetting the barcode read counts to *P. aeruginosa* strains only. There were no compounds, whose total count for all *P. aeruginosa* strains combined, was reduced to less than 10% of their median count in vehicle control wells.

Next, to identify hypomorphic strain specific inhibitors from the sequencing output, we developed an analysis metric called standardized log fraction (SLF). In comparison to the log-fold change metric used previously in PROSPECT^15^ which measures a strain’s sensitivity to a compound relative to DMSO control and thus reflects the activity of a compound on a strain without regard to the specificity of the compound for one or a few strains, SLF considers each strain’s growth relative to that of all the other strains in the same well, thereby taking into account any preferential compound selectivity for certain strains within the pool. SLF of each strain exposed to a given compound is obtained by calculating the fraction of the population taken up by each strain in the well, standardized relative to each strain’s fraction of the population grown in DMSO, after a log_10_ transformation (Method Details). A more negative SLF indicates increased inhibition of a strain’s fractional growth in a compound relative to that in DMSO, in the multiplexed screen. Thus, SLF incorporates both potency and specificity into a single metric, highlighting compounds that affect a small number of strains, while de-emphasizing those that affect the majority of strains.

Disregarding the two pan-active compounds, 0.6% (6,786 of 1,186,394) of all remaining compound-strain interactions had an SLF less than −3. In total, these results involve 10% (5,519 of 53,929) of the total number of screened compounds (Table S2B). When focusing on *P. aeruginosa* and its hypomorph strains alone, 0.08% (818 of 970,686) of the compound-strain interactions had SLF less than −3, representing 1% (637 of 53,929) of the total number of screened compounds (Figure S2C). To be conservative, these values were calculated by selecting the replicate with the higher or less negative SLF to ensure reproducibility of the level of inhibition. In contrast, no hits would have been identified if screening wildtype *P. aeruginosa* alone, based on total *P. aeruginosa* counts relative to those in vehicle control. Thus, multiplexed screening against a hypomorph strain pool identified hits with anti-pseudomonal activity, albeit against hypomorphs, compared to screening against wildtype *P. aeruginosa*.

To validate compound-strain activities identified by the SLF method, a subset of compounds active against *P. aeruginosa* strains were separately tested against wildtype and individual hypomorphs in a non-pooled format using an orthogonal absorbance (OD_600nm_)-based assay. Out of the 637 compounds that had SLF less than −3 against at least one *P. aeruginosa* strain, we tested 126 available compounds (160 compound-strain combinations) in the demultiplexed format. Of the 160 compound-strain combinations, 59 (36%, corresponding to 49 compounds) were confirmed to have >30% growth inhibition (Table S2C). Of note, while growth inhibition in this demultiplexed secondary assay serves to confirm a negative SLF value in the primary screen, the increased sensitivity of the multiplexed assay due to conditions of competitive growth combined with the more sensitive sequencing-based readout results in a larger number of hits compared to non-competitive growth conditions when testing a single strain in an absorbance-based assay. As expected, the confirmation rates were higher with more extreme SLF values. For example, of the 37 compound-strain interactions (29 compounds) with SLF less than −6, 25 (68%, corresponding to 21 compounds) had >30% growth inhibition (Figure 3). Additionally, the strain-specificity of the SLF metric, where only the strains with SLF less than a given value had growth inhibition, was also confirmed at higher rates with increasingly negative SLF values and increasing thresholds of growth inhibition (Tables S2C and D).

### BRD1401 activity is specific for *P. aeruginosa* strains depleted for OprL and TolB

In the screen, we identified the aminoimidazolone BRD1401 (Figures 4A and S3; Method Details) as a small molecule with significant and specific activity (SLF = −8.25) (Figure 3B) toward the *P. aeruginosa oprL*-hypomorph (Figure S4A and PA19 in Table S1G). The OprL ortholog in other gram-negative species, Pal, and its known associated protein complex, TolQRAB, have been shown to mediate linkage between the OM and PG, particularly during bacterial cell division.^40–42^ BRD1401 had no activity against any other bacterial species (Figure S5A) or against wild-type *P. aeruginosa* when tested at 250 µM, the limit of its solubility, or other *P. aeruginosa* hypomorph strains except for the *oprL* and *tolB*-hypomorphs (Figure 4B) in the demultiplexed assay.

To confirm the dependence of BRD1401’s activity on levels of OprL, we created a strain in which we could control OprL expression. We engineered a PA14 strain with an arabinose-inducible copy of *oprL*, henceforth referred to as the *oprL*-knockdown strain (*oprL*-KD; Figure S4A and PA23 in Table S1G). We confirmed depletion of OprL by 95-98% at the protein level in *oprL*-KD at an arabinose concentration of 0.063% while it was reduced by 89% in the *oprL*-hypomorph screening strain (Figures S5B and C). *oprL*-KD grew at a slower rate with 0.063% arabinose in the culture medium, growing up to 60% of the maximum cell density after 24 hours relative to 0.5% arabinose (Figure S5D). Depleting OprL even further by lowering arabinose concentrations both further impaired growth of this strain (Figure S5D), as expected for an essential gene, and increased susceptibility to BRD1401 (Figure 4C), thus confirming a chemical-genetic interaction between OprL and BRD-1401. We observed that BRD1401 was bactericidal towards *oprL*-KD at 125 μM when OprL was depleted (Figure S5E).

Because orthologs of Tol-OprL complex in other species are known to be essential for membrane integrity,^43–45^ one possible explanation for the hypersensitivity of the corresponding hypomorphs to BRD1401 is increased small molecule intracellular accumulation facilitating engagement with an intracellular target. We thus tested whether *oprL*-hypomorph has a permeability defect by measuring membrane integrity of the strain (Figure S5F). Using an AmpC activity-based assay to measure loss of OM integrity which results in leakage of the periplasmic beta-lactamase AmpC protein into the supernatant of log-phase cultures,^46^ we observed no difference in supernatant AmpC levels between *oprL*-hypomorph and wildtype PA14, suggesting that the OM of the *oprL*-hypomorph strain is intact. The lack of non-specific increases in permeability was further confirmed by the absence of significant sensitization of the *oprL*-hypomorph to a panel of standard antibiotics that require cell entry and intracellular accumulation, such as ciprofloxacin, amikacin, and doxycycline (Figure S5G). Thus, the activity of BRD1401 in this strain is not because of non-specific increased permeability of its OM. Additionally, we also observed no hypersensitization of efflux deficient *P. aeruginosa* strain, PAO397^47^ to BRD1401 (Figure S5H and Table S1G), which is further consistent with an OMP target.

### BRD1401 does not directly target OprL or TolB

Since the *tolB*-hypomorph was similarly sensitized, like the *oprL*-hypomorph, we also engineered arabinose regulated, knockdown strains of *tolB* (*tolB*-KD; PA24 in Table S1G) and its cytoplasmic membrane components, TolQRA (*tolQRA-*KD; PA25 in Table S1G). Indeed, BRD1401 activity was dependent on TolB levels but not TolQRA (Figure 4D).

Given the observed chemical-genetic interactions between BRD1401 with OprL and with TolB, we tested whether BRD1401 directly binds to either OprL or TolB by two complementary biophysical methods: Differential Scanning Fluorimetry (DSF) and Saturation-Transfer Difference Nuclear Magnetic Resonance (STD-NMR).^48,49^ Neither biophysical approach indicated a direct interaction between BRD1401 and the two proteins (Figures S6B and C). Purified OprL and TolB did bind to each other strongly in a Biolayer Interferometry readout (BLI) (Figure S6D), thus mirroring the known interaction between OprL and TolB orthologs in *E. coli*.^50,51^ The presence of BRD1401 however did not affect the off rate for OprL-TolB complex, suggesting that this molecule does not directly disrupt this protein complex (Figure S6D).

To definitively rule out OprL as the direct target of BRD1401, we leveraged a literature report of an *oprL* deletion strain (Figure S4A; PA22 in Table S1G),^52^ whose existence seemingly contradicted genome-wide negative selection studies (TnSeq) identifying it as an essential gene.^16^ While we indeed confirmed the *oprL* deletion in this published strain, henceforth referred to as Δ*oprL*-MW, by PCR (Figure S4B), whole genome re-sequencing revealed a number of putative suppressor mutations in a diverse set of pathways including drug efflux systems, putative LPS modification enzymes and PG modulating proteins that may have enabled deletion of this essential gene (Figure S4C). Meanwhile, *oprL* deletion in this strain allowed us to test whether OprL could be the target of BRD1401, as its presence would be required for BRD1401 activity. In fact, Δ*oprL*-MW was still susceptible to BRD1401 (Figure 5A), thus confirming the biochemical data suggesting that OprL cannot be the direct or sole target of BRD1401. Instead, BRD1401 must be inhibiting a protein or pathway that is genetically linked to OprL.

**Figure 5.**
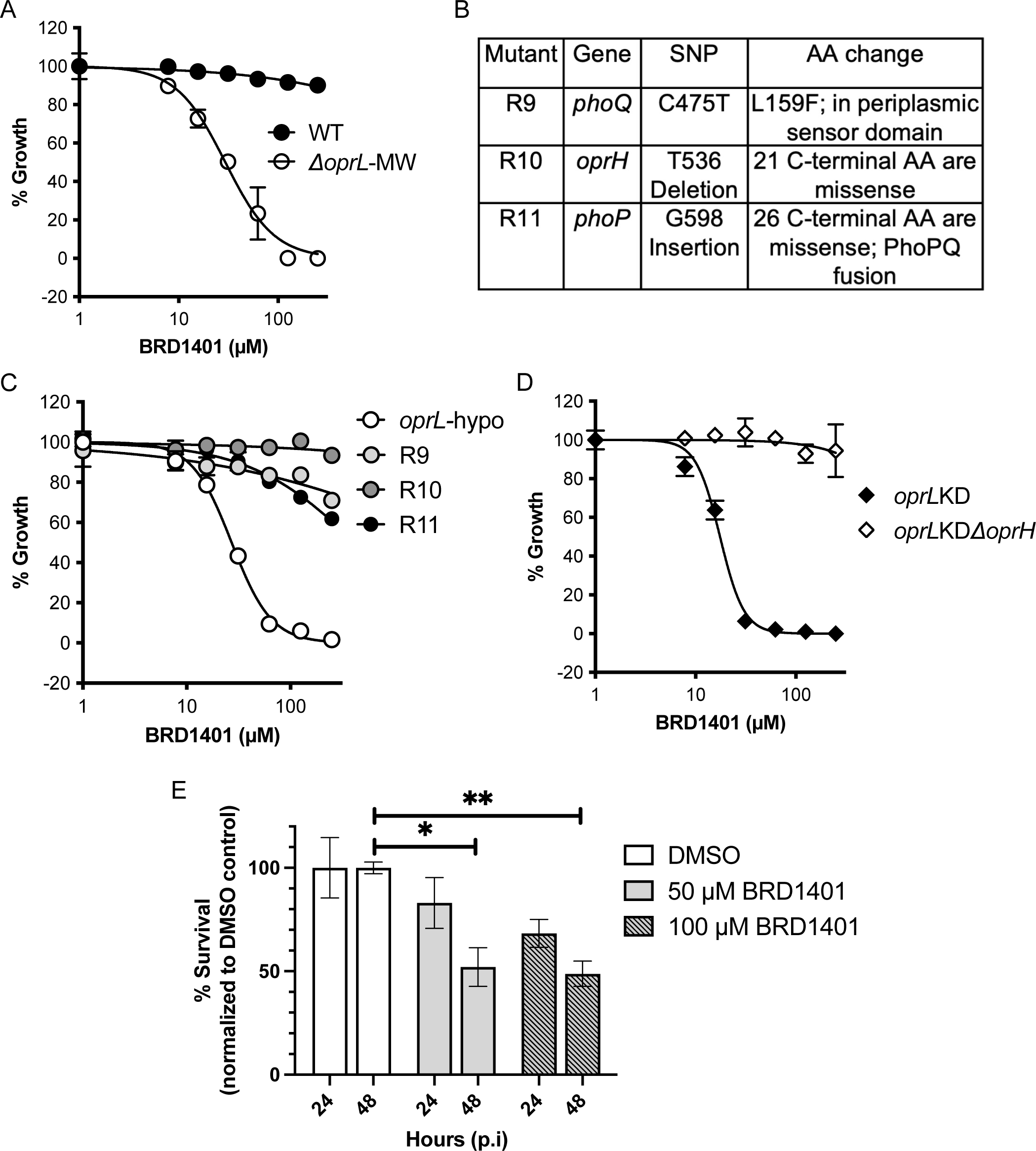
OprH is necessary for BRD1401 activity. (A) BRD1401 activity towards the Δ*oprL*-MW strain and the isogenic wildtype strain (WT) as measured by an absorbance (OD_600nm_)-based growth assay. Normalized growth, calculated in % relative to vehicle and positive controls, is plotted against BRD1401 concentration (n= 3, error bars indicate SD). (B) Table listing mutations identified in the *oprH-phoPQ* operon by whole-genome sequencing of the three *oprL*-hypomorph clones resistant to BRD1401. SNP indicates the single nucleotide polymorphism detected in each of the clones and AA indicates the amino acid residue that is mutated. (C) BRD1401 activity towards the parent *oprL*-hypomorph strain and the three resistant clones, R9, R10 and R11, measured by an absorbance-based assay. Normalized growth is plotted against BRD1401 concentration (n= 3, error bars indicate SD). All three clones show high-level resistance to BRD1401. (D) BRD1401 activity towards the *oprL*-KD and *oprL*-KDΔ*oprH* strains grown in media with 0.063% arabinose (low OprL). *oprL* expression is under the control of the P_araBAD_ promoter. Normalized growth measured by an absorbance-based assay is plotted against BRD1401 concentration (n= 3, error bars indicate SD) and shows that OprH is required for BRD1401 activity. (E) BRD1401 activity towards intracellular *P. aeruginosa*. Shown is % survival, calculated from colony forming units and normalized to DMSO vehicle control, of *P. aeruginosa* PAO1 strain recovered from bladder epithelial cells after treatment for 24 or 48 hours with indicated concentrations of BRD1401 (n= 3, error bars indicate SEM). * and ** indicate *p* < 0.05 and *p* < 0.005, respectively, calculated using an unpaired Student’s parametric *t*-test with Welch’s correction.

### Both OprH presence and OprL depletion are essential for BRD1401 activity

To gain further insight into the functional interaction between BRD1401 and OprL, we generated mutants resistant to BRD1401 in the *oprL*-hypomorph background. Initial Sanger sequencing of the manipulated genetic regions revealed no mutations while whole genome re-sequencing of three independent resistant clones revealed distinct mutations in the non-essential *oprH-phoPQ* operon (Figure 5B). PhoQ and PhoP are the membrane sensor and response regulator, respectively, of a two-component signal transduction system that regulates a downstream transcriptional program in response to magnesium sensing, including the upregulation of *oprH* (Figure S7A).^53,54^ Mutations in two of the clones fell in the PhoPQ system: mutant R9 carried a mutation in PhoQ’s periplasmic sensor domain while mutant R11 had a mutation leading to mistranslation of the C-terminus of the DNA binding domain of PhoP leading to a PhoP-PhoQ fusion protein. The third clone, R10, had a mutation in the C-terminus of OprH leading to mistranslation of the C-terminal 20 amino acids and loss of the terminal β-strand that is critical for protein folding and structure. All three mutant clones were resistant to BRD1401 with R10 displaying the most significant resistance relative to parent *oprL*-hypomorph strain (Figure 5C). These mutations suggested that resistance might be a consequence of the absence of functional (R10) or low expression levels (R9, R11) of *oprH*, with the latter reflected in transcriptional studies showing that *oprH* is down regulated in both resistant clones R9 and R11 (Figure S7B). In contrast, *oprH* is upregulated ∼2-fold in both the *oprL*-hypomorph and Δ*oprL*-MW strains relative to the wildtype PA14 strain demonstrating a correlation between *oprH* expression and BRD1401 sensitization (Figures S7C and D). This relationship is further supported by the fact that exposure of the *oprL*-hypomorph and Δ*oprL*-MW strains to BRD1401 also results in *oprH* upregulation (Figures S7C and D). Together, these results demonstrate that in addition to OprL, BRD1401 also has a chemical-genetic interaction with OprH.

OprH is a non-essential, small, OM-embedded, β-barrel protein in *P. aeruginosa* that directly interacts with LPS through its extracellular loop domains and plays a role in LPS packing in magnesium-poor environments.^18,19^ To test whether OprH is indeed required for BRD1401 activity, we deleted the non-essential *oprH* gene in *oprL*-KD and found that the mutant (PA27 in Table S1G) was completely resistant to BRD1401 (Figure 5C). Complementation of *oprH* in the deletion strain led to an *oprH* dose-dependent return in susceptibility to BRD1401 and even hyper-susceptibility with high *oprH* levels (Figure S8A). OprH is thus required for BRD1401 activity and loss of OprH function is sufficient for BRD1401 resistance. Of note, overexpression of *oprH* was unable to confer BRD1401 activity to wildtype PA14 when tested at the limit of BRD1401 solubility (Figure S8B).

Supporting BRD1401 activity requiring two simultaneous events, *oprH* expression and lowered *oprL* expression under normal growth conditions (LB medium), we found that hyperosmolarity induced by high salt, a condition that induces these corresponding expression changes in *oprH* and *oprL* increased BRD1401 activity in *oprL*-hypomorph (Figures S7E and S8C). Hyperosmolarity is also a known activator of the PhoPQ signaling system in *E. coli* and could explain induction of OprH in *P. aeruginosa* in high salt medium.^55^ Remarkably, in the presence of high salt, we observed modest (50-60% growth inhibition) but significant BRD1401 activity against wildtype *P. aeruginosa* for the first time (Figure S8D). This wildtype activity was also dependent on OprH since PA14Δ*oprH* was completely resistant (Figure S8D). Meanwhile, while over-expression of OprH had no effect on BRD1401 activity in wildtype PA14 under normal salt conditions, overexpression of OprH under high salt conditions now increased sensitization to BRD1401 (Figure S8D).

### BRD1401 reduces *P. aeruginosa* intracellular survival

The condition specific activity of BRD1401 raised the question of whether these conditions might be relevant to an infection setting, with implications for BRD1401 activity. Interestingly, a recent study reported >1000-fold up regulation of OprH after adherence of *P. aeruginosa* to epithelial cell line *ex vivo*.^56^ In contrast, after 2 hours of infection of a bladder epithelial cell line, both OprH and OprL have been reported to be downregulated in the *P. aeruginosa* strain PAO1, by 3.6-and 3-fold, respectively.^57^ When we infected the bladder epithelial cell line with the PAO1 strain, we observed a BRD1401 dose-dependent decrease in PAO1 colony forming units (CFU) at 48 hours post infection with up to 48% reduction at 100 µM of BRD1401 (Figure 5E). BRD1401 was not cytotoxic at the tested concentrations up to 48 hours (data not shown). Thus, BRD1401 has some activity against wildtype *P. aeruginosa* under specific growth conditions, including during infection.

### OprH is not a BRD1401 transporter

Since OprH is non-essential, OprH must facilitate BRD1401 activity either directly or indirectly. Although OprH has sequence homology with known porins, purified OprH in a lipid bilayer does not induce membrane conductance, indicating it would be an atypical porin.^58^ Nevertheless, we considered the possibility that OprH could act as a BRD1401 transporter required for intracellular accumulation and thus engagement of BRD1401 with an intracellular target. However, intracellular BRD1401 levels in *oprL*-KD strain or *oprL*-KD-derived strains with deletion or over-expression of *oprH* did not correlate with BRD1401 activity observed in these strains (Figures S8A and S8E), suggesting that OprH must play an alternative functional role.

### BRD1401 binds OprH to disrupt the OprH-LPS interaction

To test if BRD1401 directly binds to OprH, we purified StrepII-tagged OprH (Figure S9A) from *P. aeruginosa*, confirming protein expression and purification by intact mass-spectrometric analysis (Figure S9B). Purified OprH was folded as indicated by the shift in molecular weight after boiling^18^ and by differential scanning calorimetry (Figures S9A and C). We then utilized saturation transfer difference nuclear magnetic resonance (STD-NMR), using the aromatic protons of BRD1401 as reporters, to determine binding of BRD1401 to OprH. We included five BRD1401 analogs, three with different substitution patterns on the phenyl ring (IA-2, 1401-A, and 1401-B), one with an additional methyl group on the central imidazole ring (IA-1) and one with a carboxamide on the pyrimidine ring (1401-C) (Figure 4A). Their bactericidal activities against *oprL*-KD ranged from being similar to (1401-C), reduced (1401-A and 1401-B) or lost (IA-1 and IA-2) relative to the parent BRD1401(Figure S9E). Using a binding parameter of percent ΔSTD (Method Details), the ΔSTD signal of BRD1401 and the 5 analogs in the presence and absence of OprH were positively correlated with growth inhibitory activity in *oprL*-KD (Figures 6A, 6B, S10A, S10B and S10C). Furthermore, as binding of a small molecule to a larger molecule will enhance its transverse (T_2_) relaxation rates, we performed Carr-Purcell-Meiboom-Gill (CPMG) experiments on a subset of these analogs to monitor the effect of binding to OprH on the signal decay rate of each compound’s aromatic protons.^59^ Indeed, the measured relaxation rates for active analogs markedly differed in the presence and absence of OprH indicating binding, whereas the inactive analogs IA-1 and IA-2 showed no difference (Figures S10D and E). Overall, the NMR analyses show that BRD1401 and its active analogs bind to OprH, and that binding potency correlates with whole-cell activity.

**Figure 6.**
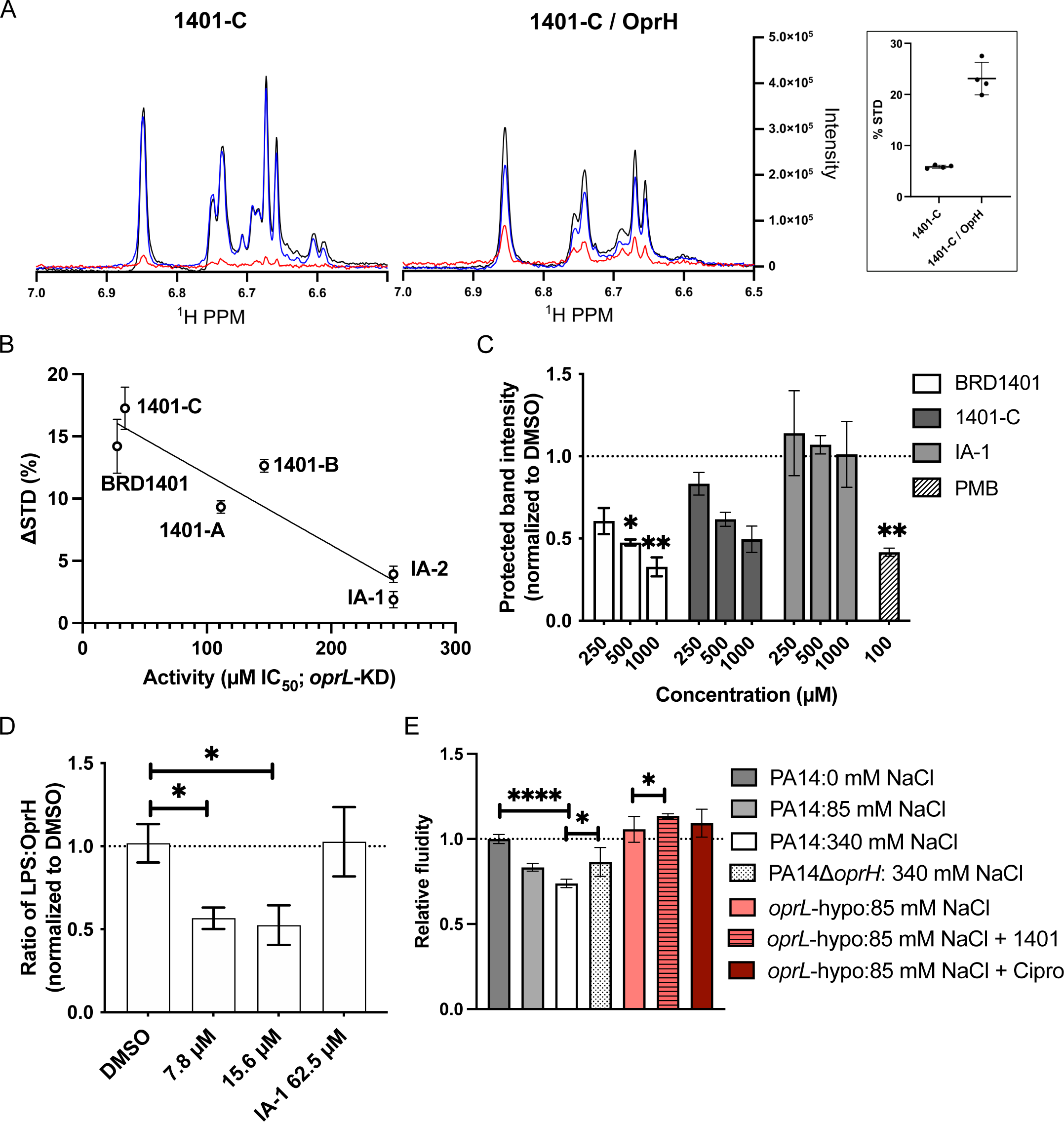
BRD1401 activity is linked to OprH binding and function. (A) BRD1401 binds to OprH. Overlaid 1H-spectra of the aromatic region chemical shifts and intensities of 500 µM 1401-C solution in dialysis buffer (left) and in the presence of 15 µM *P. aeruginosa* OprH (right; Method Details) without irradiation of OprH (black) and with irradiation of OprH (blue), inducing a decrease in 1401-C signal intensity due to a negative Ligand-Protein distance-dependent Nuclear Overhauser Effect (NOE). The difference spectrum (STD in red) between black and blue spectra corresponds to the magnitude of the NOE indicating binding dependent loss of 1401-C signal in presence of OprH. Calculated %STD for 1401-C analog with and without OprH is plotted in the box. (B) Growth inhibitory activity, measured as IC_50_, of BRD1401 and 5 analogs against *oprL*-KD strain is inversely correlated with the calculated ΔSTD binding parameters (Method Details) from STD-NMR experiments with purified *P. aeruginosa* OprH protein. Fit is simple linear regression; data represents 2 independent experiments for each analog. (C) BRD1401 decreases LPS binding to *P. aeruginosa* OprH in vitro as measured using a trypsin digestion protection assay (Method Details). Purified OprH protein was incubated with Kdo2-Lipid A with or without indicated concentrations of compounds (BRD1401 and 1401-C, which are active towards *oprL*-KD strain; inactive analog IA-1; and positive control polymyxin B (PMB) and subsequently digested with trypsin. Digested samples were boiled and run on an SDS-PAGE gel and the band intensity of the LPS-protected band at ∼14.5 kDa (Figure S10F) was quantified using Fiji (ImageJ). BRD1401 treatment reduced the band intensity, indicating decreased binding of OprH to Kdo2-Lipid A. Band intensity normalized to vehicle DMSO control (dashed line = 1) is plotted (n=2 or 3; error bar indicates SEM). * and ** indicate *p* < 0.05 and *p* < 0.01, respectively, calculated using a One sample *t*-test, in comparison to the vehicle control. (D) BRD1401 decreases LPS binding to OprH in the OM of *P. aeruginosa*. StrepII-tagged OprH was pulled down using Streptactin beads from the lysates of the *oprL*-KDΔ*oprH/pRha-StrepII-oprH* strain exposed to DMSO vehicle control, BRD1401 at 7.8 or 15.6 µM, or the inactive analog (IA-1) at 62.5 µM, followed by western blotting with anti-LPS-O10 and anti-StrepII antibodies (Method Details). Shown is the ratio of LPS band intensity relative to OprH band intensity in the elution samples, quantified by Fiji (ImageJ) and normalized to a vehicle (DMSO) control sample (n=3; error bar indicates SEM). * indicates *p* < 0.05 calculated using an unpaired Student’s parametric *t*-test with Welch’s correction. (E) Impact of BRD1401 treatment, *oprH* deletion, or *oprL* depletion on membrane fluidity in *P. aeruginosa*. Wildtype PA14, PA14Δ*oprH,* or *oprL*-hypomorph strain was grown in media with the indicated concentrations of salt (NaCl) and exposed to 0.0625 µM ciprofloxacin (Cipro) or 250 µM BRD1401 (1401) where indicated. After exposure, cultures were treated with a pyrene lipophilic dye and processed to measure relative fluidity normalized to PA14 grown with no salt, as described in Method Details (n=3-12 biological replicates; error bars indicate SEM). High salt decreases membrane fluidity, while OprH deletion, OprL depletion and BRD1401 increase membrane fluidity. * and **** indicate *p* < 0.05 and *p* < 0.0001, respectively, calculated using an unpaired Student’s parametric *t*-test with Welch’s correction.

We next sought to determine if BRD1401 binding to OprH prevents its known interaction with LPS.^18,19^ Adapting a previously described assay to detect in vitro binding of crude LPS to OprH extracellular loop 3 based on LPS binding protecting OprH from trypsin digest,^18^ we measured whether BRD1401 or its active analog, 1401-C, prevented the interaction of recombinant OprH with the LPS analog Kdo2-Lipid A. Trypsin digestion of OprH is monitored by detecting decreasing intensity of a ∼14.5 kDa protected band derived from recombinant OprH by its interaction with LPS (Figure S10F). BRD1401 and its active analog 1401-C significantly reduced Kdo2-Lipid A protection of OprH from trypsin digestion in a dose dependent manner (Figures 6C, S10G and S10H). Meanwhile, inactive analog IA-1 had minimal effect (Figures 6C and S10I). Polymyxin B served as a positive control since it is known to disrupt the Lipid A interaction with OprH (Figure 6C).

Finally, we tested whether BRD1401 disrupts OprH’s interaction with LPS in the OM of *P. aeruginosa* in whole cells.^18^ We introduced an episomal copy of StrepII-tagged OprH into the *oprL*-KDΔ*oprH* strain (PA32 in Table S1G) to enable LPS pulldown. We confirmed the tagged protein to be functional based on increased sensitization to BRD1401 upon expression of StrepII-tagged OprH in the resulting strain (Figure S11A). LPS could be detected in elution samples after pulldown with biotinylated beads from bacterial lysates with StrepII-OprH, but not untagged OprH (Figure S11B). Sub-lethal treatment of *oprL*-KDΔ*oprH/pRha-StrepII-oprH* with BRD1401 resulted in a 30-50% decrease in LPS band intensity (normalized to StrepII-OprH protein) relative to vehicle control, whereas the inactive analog IA-1 had no impact (Figures 6D and S11C). BRD1401 treatment did not affect OprH protein expression or stability as indicated by unchanged StrepII-OprH band intensities in the whole cell lysate samples (Figure S11C). Taken together, BRD1401 binds OprH to specifically disrupt LPS binding to OprH both in vitro and in bacterial cells.

### BRD1401 impacts membrane fluidity

Since BRD1401 interferes with LPS binding to OprH, we wondered if it could affect LPS organization in the OM, either to increase membrane permeability, like the cationic antimicrobial peptide colistin, or to change membrane fluidity. To examine membrane permeability, we monitored the uptake of the membrane impermeable DNA-binding dye ethidium bromide in the *oprL*-hypomorph strain after treatment with colistin or BRD1401. Unlike colistin, which caused an immediate increase in dye intake, BRD1401 had minimal effect (Figure S11D) suggesting no pore formation or dramatic destabilization of the OM that results in increased membrane permeability.

However, when we used a lipophilic pyrene dye to measure changes in membrane fluidity,^60^ we found that under the high salt conditions in which we had seen increased BRD1401 activity, there was decreased membrane fluidity in wildtype PA14, as has been reported in other species under these conditions (Figure 6E).^61^ Further, both depletion of OprL in the *oprL*-hypomorph and deletion of OprH in PA14Δ*oprH* increased membrane fluidity relative to wildtype PA14. Finally, BRD1401 further increased membrane fluidity of the *oprL*-hypomorph (Figure 6E). Taken together, these results point to a functional role for OprL and OprH in maintaining membrane fluidity, and BRD1401 disrupting LPS structure in the OM to increase fluidity, which may account for its activity in wildtype *P. aeruginosa* in high salt conditions.

### OprH directly binds to OprL

With BRD1401’s unveiling of a functional interaction between OprH and OprL, we sought to better define this interaction genetically and biochemically. First, we examined growth in *oprL*-KD when OprL is depleted as a function of OprH levels and indeed observed a significant fitness defect that was OprH dose-dependent (Figure 7A). Based on this genetic interaction, we next sought to determine whether there is a biochemical interaction between OprH and OprL. We introduced a P_araBAD_ driven FLAG-tagged OprL at the *att*Tn7 site in *oprL*-KD, along with a StrepII-tagged OprH expressed from the native locus and an untagged OprH expressed episomally under the control of a rhamnose-inducible promoter (PA39 in Table S1G). Immunoprecipitation of FLAG-tagged OprL revealed a direct, specific interaction with StrepII-tagged OprH that could be competed out by overexpression of untagged OprH. We normalized the OprL-FLAG protein band intensity to the StrepII-OprH protein band intensities in elution samples to accurately measure specific binding and observed a significant decrease in OprL-FLAG pull down by StrepII-OprH in presence of excess, untagged OprH (Figure 7B). Thus, there is a direct and specific interaction between OprL and OprH, which could explain the dependency of BRD1401 activity on both these proteins. Note, there was a mild decrease in StrepII-OprH protein levels in the lysate and thus in the eluted samples upon increasing induction of untagged OprH by rhamnose; as Strep-OprH is expressed under the control native promoter, the decrease in StrepII-OprH may suggest a transcriptional or post-translational regulation of OprH protein levels (Figure S11E). While BRD1401 disrupts the interaction between OprH and LPS, it does not disrupt the OprL-OprH interaction. In fact, BRD1401 resulted in a small increase in OprH pulled down with OprL, which could be explained either by BRD1401 increasing the affinity of OprL for OprH or by the fact that we observed a modest increase in abundance of OprL-FLAG when treated with BRD1401 which might hint at post-translational stabilization of OprL by BRD1401 (Figure S11F). Taken together, as OprL is known to associate with PG and we have shown that OprH interacts with LPS, these findings reveal that an OprH-OprL complex directly links PG and OM in *P. aeruginosa*.

**Figure 7.**
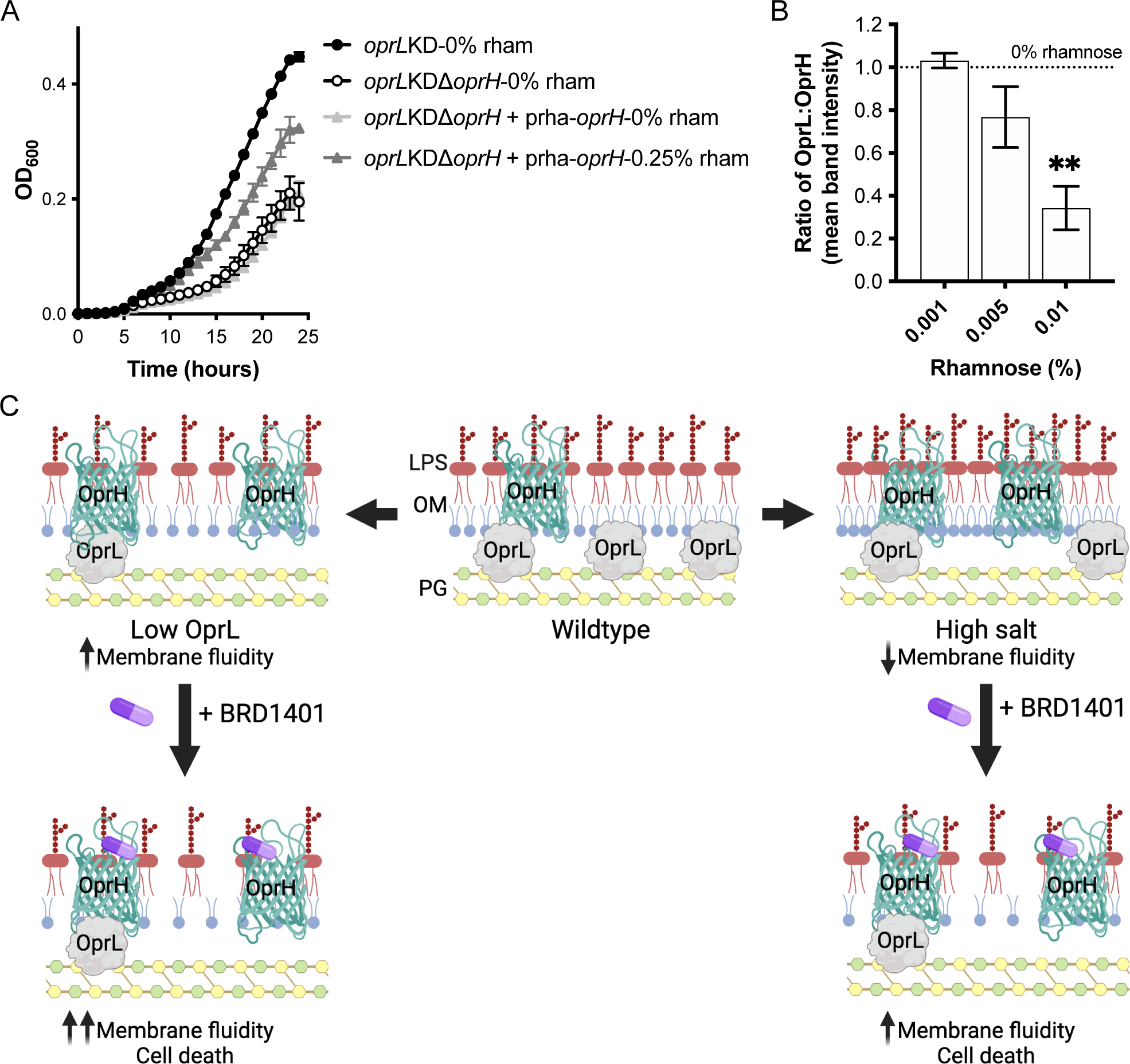
OprH binds to OprL. (A) OprH depletion impairs growth when OprL is present in low levels. Growth kinetic, measured by absorbance (OD_600nm_), of the *oprL*-KD strain, the *oprL-*KD*ΔoprH* or the *oprL-KDΔoprH/pRha-oprH* in media with 0.063% arabinose to downregulate *oprL* expression and 0% or 0.25% rhamnose (rham) to drive episomal *oprH* expression (n = 3, error bars indicate SD). (B) OprL binds specifically to OprH in *P. aeruginosa*. StrepII-OprH was pulled down using Streptactin beads from lysate made from the *oprL-FLAG-KD StrepII-oprH/pRha-oprH* strain grown in the presence of varying doses of rhamnose to drive the expression of free, untagged OprH. Following elution with excess biotin, samples underwent western blot analysis with an anti-FLAG antibody recognizing OprL-FLAG and an anti-Strep antibody recognizing StrepII-OprH, followed by quantification of the ratio of OprL-FLAG:StrepII-OprH band intensities (first normalized to StrepII-OprH band intensity and then further normalized to the control sample (0% rhamnose; dashed line = 1) using Fiji (ImageJ; Method Details). With increasing concentrations of rhamnose resulting in increasing expression of free, untagged OprH to compete with StrepII-OprH, the ratio of OprL-FLAG to StrepII-OprH decreased. * indicates *p* < 0.05 calculated using a One sample *t*-test in comparison to the 0% rhamnose control sample. (C) Proposed model for mechanism of action of BRD1401. OprL depletion increases membrane fluidity, with the addition of BRD1401 further increasing membrane fluidity and cell death. Meanwhile, in high salt conditions, membrane fluidity of wild-type *P. aeruginosa* decreases, but the addition of BRD1401 increases membrane fluidity and cell death ensues. *Created with Biorender.com*.

## DISCUSSION

A mini-PROSPECT screen with a pool of bacterial knock-down strains is a novel strategy to identify species and target-specific inhibitors that can serve as probes of protein and pathway interactions. With genome-wide negative selection studies (Tn-Seq) across *P. aeruginosa* strains under diverse conditions revealing four major essential pathways associated with the OM,^16^ we focused on the corresponding four complexes (Figure 1A) which are primarily embedded or associated with the OM and hence do not require permeation of two membranes by putative inhibitors. Three of the four complexes, Bam, Lpt and Lol, are also essential across other pathogenic gram-negative bacterial species and have been the focus of intense antibiotic discovery efforts over the past decade.^7^ mini-PROSPECT is the first attempt to identify target-specific inhibitors for these important essential OMP/OMAPs in *P. aeruginosa* by combining a whole cell phenotypic screen with a knockdown-based genetic strategy.

Relative to the genome-wide PROSPECT strategy previously reported in *M. tuberculosis,*^15^ we have adapted PROSPECT to focus on a smaller set of prioritized genes in *P. aeruginosa*. We made technical modifications in the chemical screen, specifically in strain and library construction, with the latter being simpler and more cost-effective, and developed a novel computational pipeline with the SLF metric to prioritize the strain-specificity of identified inhibitors. In the future, CRISPRi technologies are an additional alternative method for engineering of strains with gene product depletion. Importantly, in contrast to PROSPECT that can be used to predict target or mechanism of action (MOA) directly from the “genome-wide” chemical genetic interaction profiles,^15^ mini-PROSPECT does not so easily lend itself to direct target prediction because of the relatively small number of essential genes included in the multiplexed pool. This is even more true if querying a small set of targets that may interact functionally or biochemically, as the same inhibitor could have activity against several strains with interacting targets. Meanwhile, without the additional information on the susceptibility of hypomorphic strains that are not included in the screening pool, including strains with corresponding targets/pathways that may be even more susceptible than ones included in the pool, mini-PROSPECT cannot reliably predict MOA. Nevertheless, mini-PROSPECT can identify small molecules with specific activities against engineered strains that can be used to probe the biology of the hypomorphs, irrespective of whether the depleted gene product is in fact the direct target.

From this mini-PROSPECT screen, we prioritized BRD1401 for mechanistic study and target identification efforts because of its *P. aeruginosa*-specific activity and unique chemical-genetic interaction with the essential OMAP, OprL, which with its known associated protein complex, TolQRAB, mediate linkage between the OM and PG.^40,41^ Despite this specificity, genetic and biochemical studies revealed that rather than OprL, the non-essential β-barrel OMP, OprH, is the target of BRD1401 in *P. aeruginosa*. Of note, the absence of exact OprH orthologs in other gram-negative pathogenic bacteria could explain the species-specific activity of BRD1401 (Figure S5A). Homology analysis of OprH protein sequence across all protein sequences available on NCBI indicated existence of highly similar proteins only within the genus *Pseudomonas*, with OmpW from species such as *E. coli* being a distant homolog (Figure S12). OmpW is described as a putative transporter of hydrophobic molecules based on in vitro conductance and structure studies;^62^ however, OprH lacks the extracellular domains implicated in this transporter function. OprH has previously been described to stabilize LPS,^18,19^ particularly in the context of low magnesium environments,^17,53^ but not directly linked to OprL. Our studies with BRD1401 thus unveil a novel genetic and biochemical interaction between OprL and OprH, thereby providing an additional link between the PG and OM in *P. aeruginosa*.

In *E. coli,* while the OprL homolog, Pal, and the Tol complex are not essential, they do play an active role in regulating PG remodeling during cell division in *E. coli*.^63^ In contrast, this complex is essential in *P. aeruginosa*, suggesting that OprL may play an even more vital role in regulating PG homeostasis and OM stabilization, particularly during cell elongation and division. The latter could be mediated through OprL’s interaction with OprH, with OprH then binding LPS. In *E. coli,* multiple OM β-barrel proteins such as OmpF, FhuA, BtuB and FepA are known to bind LPS.^64–66^ Thus, other OMPs could be compensating for OprH absence in the *P. aeruginosa* deletion strain, even while chemical inhibition with BRD1401 initiates a more rapid disruption of LPS organization in OM, particularly in the setting of weakened OM-PG linkages mediated by OprL downregulation. The decrease in LPS organization could be the cause for the increased membrane fluidity seen in the *oprL*-hypomorph strain relative to wildtype PA14 and exacerbated by BRD1401 treatment, leading to lethality. Recent atomic force microscopy studies have shown distinct LPS-laden microdomains in the OM of *E. coli*, facilitated by OMPs.^67^ Similar high-resolution analysis of the *P. aeruginosa* OM could shed light on the molecular details of BRD1401’s effect on LPS.

BRD1401 activity in vitro was dependent on specific genetic backgrounds or growth conditions, which may be related to a strain’s dependence under certain growth conditions on membrane structure and fluidity. Interestingly, BRD1401 had activity in wildtype *P. aeruginosa* in high salt, a condition characterized by decreased membrane fluidity, with BRD1401 blunting this decrease by increasing relative fluidity under these conditions (Figure 7C). If the decrease in fluidity were part of a bacterial adaptive response to high osmolarity, possibly impacting membrane permeability, OM protein-mediated functions, or downstream signaling^61,68^, perhaps BRD1401’s interference with this response could play a role in its activity. Meanwhile, under normal growth conditions (non-hyperosmolar), BRD1401-induced increase in membrane fluidity in the *oprL-*hypomorph could be similarly disrupting membrane homeostasis by increasing membrane fluidity, even more than it is already in the hypomorph due to OprL depletion (Figure 7C); meanwhile the BRD1401-induced increase may be inconsequential in wildtype bacteria under these same conditions.

BRD1401 was also active in an ex vivo infection model of bladder epithelial cells. *P. aeruginosa* is known to invade and survive in host epithelial cells^57^ and OprH has been shown to be upregulated in the initial stages of this infection.^56^ At later stages of infection OprH and OprL are both downregulated, based on RNA-seq analysis of intracellular PAO1.^57^ Thus, BRD1401 binding to *P. aeruginosa* and activity in this model might be facilitated by initial upregulation of OprH that synergizes with downregulation of OprL at later timepoints. Other studies have also found OprH to be significantly transcriptionally upregulated in a mouse model of acute pneumonia by qRT-PCR analysis and in a cystic fibrosis-adapted strain by proteomic analysis.^69,70^ Analysis of recent transcriptomic studies with *P. aeruginosa* derived from human infection samples also shows a slight, but significant up regulation (2-fold) of PhoQ, which in turn regulates OprH, and a down regulation (1.5-fold) of OprL, relative to in vitro grown cultures.^71^ These expression changes in OprL and in PhoQ were even more apparent in sputum samples from cystic fibrosis patients compared to soft tissue infection samples.^71^ Further, PhoQ mutations that lead to dysregulation of the PhoQ regulon have been reported in *P. aeruginosa* clinical strains, particularly in the context of colistin treatment and/or resistance.^72^ OprH is expected to be upregulated in these clinical strains since it is one of the most responsive effectors of the PhoPQ signal transduction system. Thus, OprH and other PhoQ regulated genes could be involved in establishment of *P. aeruginosa* infection and subsequent mitigation of host-derived stresses on the bacterial OM.

BRD1401 emerged directly from a mini-PROSPECT screen with specificity for the *oprL*-hypomorph, without wildtype activity. In the absence of extensive medicinal chemistry efforts required to achieve wildtype activity, BRD1401 nevertheless served as a useful tool to unveil an LPS-OprH interaction and OprL-OprH interaction to link PG and OM. It has the further potential to serve as a probe of the functional role of OprH and its interactions during infection.

In conclusion, BRD1401 validated the multiplexed, hypomorph-focused screening strategy as a viable approach to identify small molecules active in killing *P. aeruginosa* with novel MOA that can serve as probes of novel biology. By using hypomorphic strains that are hypersensitized to inhibitors, this strategy dramatically expands the number and diversity of small molecular probes that can be identified, relative to very small number of molecules with potent wild-type activity. We have also demonstrated the feasibility of deciphering MOA with hypomorphic strains, even in the absence of chemistry efforts to optimize molecules for wildtype activity, and of exploring their biology. Thus, we propose mini-PROSPECT as a whole cell screening approach that can be widely applied to accelerate chemical probe and potentially antibacterial discovery against diverse pathogens of interest.

## SIGNIFICANCE

Antimicrobial resistance (AMR) poses a serious threat to efficacious treatment of infectious diseases, prompting the WHO to prioritize therapeutic intervention against pathogens that contribute most to the rise of AMR. *Pseudomonas aeruginosa*, a gram-negative, opportunistic bacterium, is one of these prioritized pathogens against which novel treatments are urgently needed. A key challenge in anti-pseudomonal drug discovery is low uptake of xenobiotics due to the relatively impermeable, double-membraned cell envelope with a vast array of efflux pumps. This has limited progression of promising inhibitors identified from target-based screens. To overcome this, we chose to target *P. aeruginosa* essential outer membrane proteins (OMPs) or OM-associated proteins (OMAPs), thus avoiding a cell permeation requirement. We first describe a chemical genetic strategy for a high-throughput multiplexed, whole-cell screening pipeline with a pool of focused genetic mutants depleted for essential OMP/OMAP targets, enabling the identification of small molecule probes or growth inhibitors with novel MOA that elude conventional screening of wildtype bacteria. Second, we show how such probes, even without a significant chemistry effort, can be used to uncover new biology with the discovery of BRD1401, a *P. aeruginosa*-specific small molecule growth inhibitor with efficacy in an epithelial cell infection model. BRD1401 targets *P. aeruginosa*’s OMP OprH to disrupt its interaction with lipopolysaccharide (LPS) and reveals a direct interaction between OprH and the essential OM lipoprotein, OprL, to link LPS and peptidoglycan in the cell wall. This is the first report of a whole-cell, multiplexed screen focused on specific OMP/OMAP targets that are essential in most gram-negative pathogens. We demonstrate how using this approach can be used to reveal novel biology, in this case, about specific OMP/OMAPs and their role in cell envelope biology.

## ACKNOWLEDGEMENTS

This work was supported by an NIH U19 grant (U19AI142780) to D.T.H, an NIH R21/R33 grant (R33AI098705) to D.T.H, a Cystic Fibrosis Canada Fellowship to B.E.P, and a generous gift from Anita and Josh Bekenstein. We are grateful for the generosity of Prof. Marvin Whiteley (Georgia Institute of Technology) for providing the Δ*oprL* strain and Emily Geddes and Prof. Paul Hergenrother (University of Illinois Urbana-Champaign) for guidance on intra-bacterial compound accumulation assays. We would also like to thank Pratyusha Mogalisetti for help with biophysical assays and Anne E. Clatworthy for guidance on molecular biology techniques. RNA-Seq libraries were constructed and sequenced at the Broad Institute of MIT and Harvard by the Microbial ‘Omics Core. The Microbial ‘Omics Core also provided guidance on experimental design and conducted preliminary analysis for all RNA-Seq data.

## AUTHOR CONTRIBUTIONS

Conceptualization, B.E.P., T.Warrier, and D.T.H.; Methodology, B.E.P., T.Warrier, and D.T.H; Formal Analysis, B.E.P., T.Warrier, J.B., T.White; Investigation, B.E.P., T.Warrier, S.B., K.P.R, T.White, X.Y., P.N., K.R., K.F., and A.G.; Resources, T.K.; Writing – Original Draft, B.E.P., T.Warrier, and J.B.; Writing – Review & Editing, B.E.P., T.Warrier, K.P.R, and D.T.H; Supervision, M.F., A.B., V.K., M.S-W., N.S., and D.T.H.; Funding Acquisition, D.T.H.

## INCLUSION AND DIVERSITY

We support a diverse, inclusive and inclusive environment to conduct scientific research.

## MATERIALS AND METHODS

### Bacterial strains

*Pseudomonas aeruginosa* strains were derived from the the PA14 lab strain^73^ and grown in LB medium (US Biological Life Sciences) at 37°C, unless otherwise stated. *E. coli* strain CFT073 (ATCC#700928), *K. pneumoniae* strain RB120 (gift from Roby P. Bhattacharyya), *A. baumannii* strain 19606 (ATCC#19606) and *S. aureus* strain Newman (ATCC#25904) were grown in LB medium at 37°C for multiplexed screen and for minimum inhibitory concentration assays.

### Reagents

Molecular biology reagents were from NEB and chemical reagents were from Sigma unless otherwise noted. Primers, barcode oligos, and gene blocks (Tables S1A, S1B, S1C and S1D) were synthesized by IDT technologies.

### BRD1401 synthesis and chemical structure validation

The structure of BRD1401 was annotated as “compound A**”** (Figure S3A) in the screening library based on the source, Chembridge (Catalog number: 7582216). The same structure was available from Vitas-M (Catalog number: STK072268) and hence, we purchased new material from both ChemBridge and Vitas-M. Both purchased samples had the identical specific activity as the screening hit. However, upon chemical synthesis (Scheme S7), “compound A” was inactive (Figure S3F). ^1^H-NMR comparison between the synthesized molecule and the active batches available from ChemBridge and Vitas-M indicated the chemical structures were distinct (Figure S3B). 2D NOESY (Nuclear Overhauser Effect Spectroscopy) of the molecule available from Vitas-M showed correlation peaks between the imidazolone methylene and two phenyl hydrogens, suggesting that the position of methylene and carbonyl should be flipped (Figure S3C). Based on this, the revised regio-isomer shown as “BRD1401 (revised)” in Figure S3A was synthesized (Scheme S8). This molecule was active specifically against the *oprL*-hypomorph and its ^1^H-NMR and 2D NOESY spectra were identical to the Vitas-M molecule (Figures S3D, E and F). Therefore, we concluded that the structure of the screening hit, BRD1401, is the “BRD1401 (revised)” shown in figure S3A, which was indeed active in all subsequent experiments. This structure can exist in two tautomeric forms, A and B, shown in figure S3G. Synthetic routes of “compound A”, “BRD1401-revised” and the 5 analogs of BRD1401 are included at the end of this section.

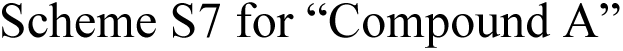
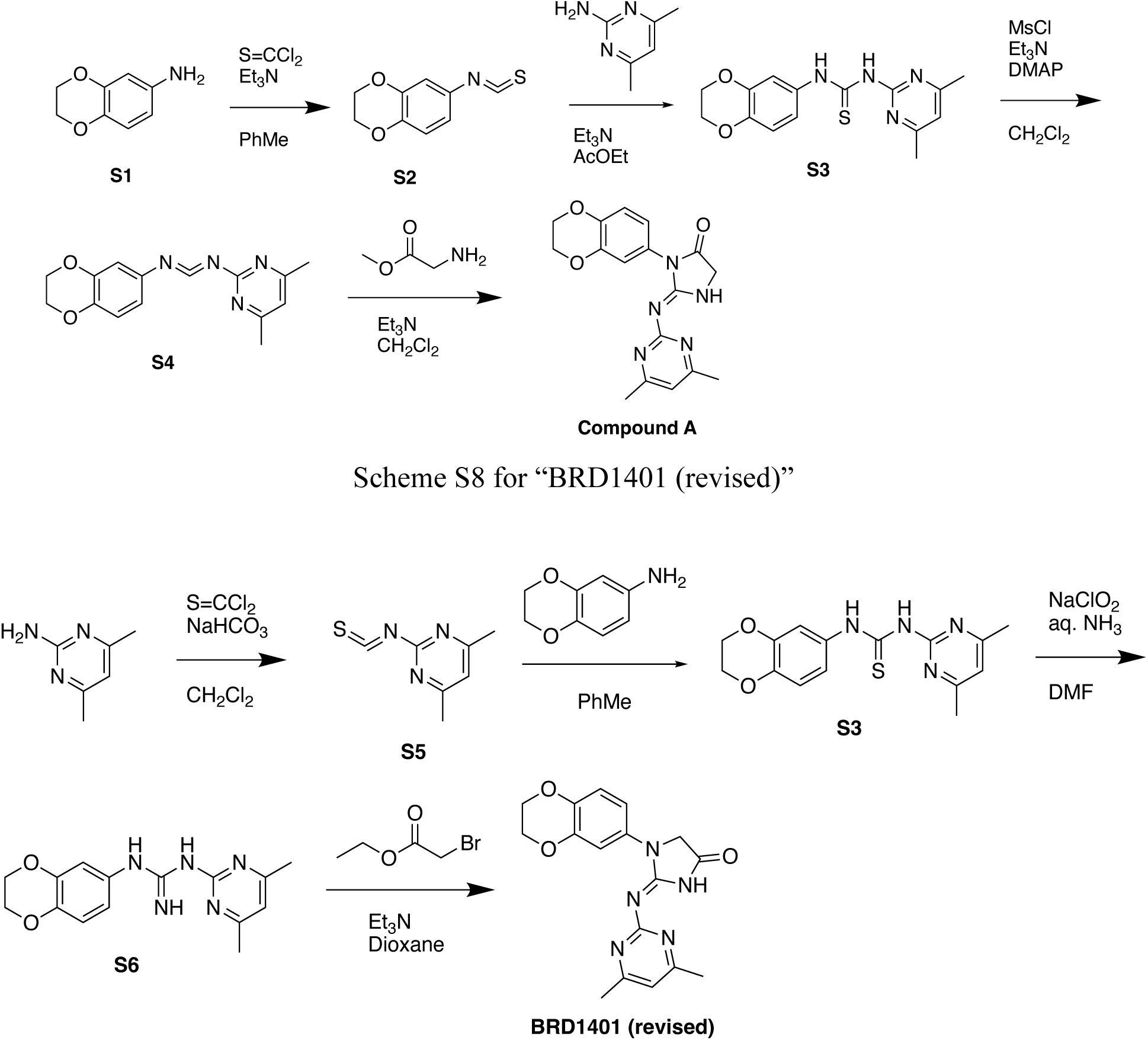

### *P. aeruginosa att*Tn7 site integration of promoter-gene combinations

All primer and plasmid information can be found in supplemental table S1. Plasmids containing promoters P_PA3-8_ were a generous gift from Robert Sauer.^33^ We modified the nomenclature from these original promoters to reflect the order of activity in *P. aeruginosa* (Table S1A). We created P_PA1-2_ by reducing the spacing between the Pribnow box and −35 sequence of P_Pa3_ from 17 bp to 15 bp and 14 bp, respectively, using site directed mutagenesis (Table S1A). To introduce the promoters and genes of interest in single copy into *P. aeruginosa*, we obtained the pUC18T-miniTn7T-Gm suicide plasmid system as a gift from Herbert Schweizer.^34^ Genes were amplified with Q5 polymerase from genomic DNA of *P. aeruginosa* strain PA14, promoters from purified plasmids, along with the mini-Tn7 plasmid using primers (Table S1B) containing 30-bp overlapping regions for Gibson assembly. Purified linear products (Qiagen) were circularized using a Gibson assembly master mix. Correct sequences were confirmed by Sanger sequencing. To introduce plasmids into *P. aeruginosa*, a four-parental mating between the host *E. coli* strain carrying the mini-Tn7 plasmid with the promoter-gene of interest, helper strains SM10 (λpir)/pTNS3 and HB101/pRK2013, and the recipient *P. aeruginosa* strain were mated and plated on LB agar plates with 10 µg/mL irgasan (Goldbio) and 30 µg/mL gentamicin as described in detail.^34,74^ Correct integration was confirmed by PCR followed by Sanger sequencing.

### *P. aeruginosa* gene deletions

A pEXG2-AmpR plasmid was derived from pEXG2-attP-Gm^75^ by digesting with NheI and EcoRV and ligating with an ampicillin resistance cassette amplified from PCR4 Blunt TOPO (Invitrogen) that was also digested with NheI and EcoRV. For each gene, a ∼1 kb kanamycin resistance cassette was amplified from pKD46^76^, and ∼1kb upstream and downstream regions were amplified with the proximal primers to the gene containing overlapping regions with the kanamycin cassette and the distal primers containing attB sequences (Table S1B). The 3 PCR amplicons were gel purified (Qiagen), added in equimolar quantities, and used in a PCR reaction to create a single ∼3 kb fragment as described.^35^ The fragment was gel purified and recombined with purified pEXG2-Amp using BP clonase II per manufacturer’s instructions (Invitrogen). Deletion plasmids were confirmed by Sanger sequencing. To knock out the gene of interest, tripartite mating between the host *E. coli* harboring a deletion plasmid for the gene, helper strain HB101/pRK2013 and the recipient *P. aeruginosa* PA14 strain was first set up. First-step recombination merodiploids were selected with 10 µg/mL irgasan (Goldbio) and 200 µg/mL carbenicillin (Invitrogen) on LB agar plates. Colonies were screened for correct merodiploids by PCR and successful clones were glycerol stocked before complement gene copies were introduced at the neutral *att*Tn7 site as described above. To perform the second recombination for substituting the native copy of the gene of interest with kanamycin-resistance cassette, cells were struck on LB agar plates with 10% sucrose and 200 µg/mL kanamycin. Colonies were screened by PCR for successful deletion and confirmed by Sanger sequencing.

### Genetically barcoding bacterial species

The parent strain PA14 was used for all *P. aeruginosa* depletion strains. The 76 bp barcode and surrounding PCR amplification sequences (Table S1C and D) were introduced at the *att-Tn7* site using the pUC18T-miniTn7T-Gm suicide plasmid system.^34^The barcoded region was introduced using the pKFT plasmid system as previously described^77^ into the *S. aureus* strain Newman in an intergenic region at position 1433140, by recombination using 1 Kb homology regions on either side of the barcode. To introduce the barcode into *E. coli*, the red recombinase expression plasmid pKD46 was first transformed into the strain CFT073, followed by transformation of a PCR fragment containing the barcode, a kanamycin resistance cassette originating from pKD4, and homology regions to integrate the barcode at position 2,689,759, as previously described.^76^ *K. pneumoniae* strain MGH48 was barcoded at the attTn7 site using the suicide plasmid delivery system pGP-miniTn7-Gm (gifted from Charles M. Dozois) as previously described.^78^ Similar to *P. aeruginosa* and *K. pneumoniae* cloning, *A. baumannii* strain 19606 was barcoded using an attTn7 integration plasmid system as previously described.^79^

### Construction of strain with arabinose-inducible *oprL* or *oprL-FLAG* expression

The P_araBAD_ promoter^80^ and downstream *oprL* or *oprL-FLAG* coding sequences were introduced into the pUC18T-miniTn7T-Gm plasmid using Gibson assembly master mix (Table S1A and B). The correct plasmid sequence was confirmed by whole plasmid sequencing and was introduced by conjugation into a PA14-*oprL* merodiploid strain created as described above. To delete the native *oprL* copy in the second recombination step to cells were struck to LB agar supplemented with 10% sucrose, 200 µg/mL kanamycin and 1% arabinose, which drove *oprL* expression from the *att*Tn7 copy. The *att*-site integration and native copy deletion were confirmed by PCR and whole genome sequencing.

### Construction of strains to modulate *oprH* and *amp*C expression

To create a Δ*oprH* strain, a pEXG2-AmpR plasmid carrying upstream and downstream sequences flanking *oprH* in PA14 and an in-frame gentamicin-resistance cassette was created using the Gibson assembly master mix (Table S1B). Merodiploid and final knock-out creation was done as described above with gentamicin-based selection. Episomal expression of native *oprH* or StrepII-tagged *oprH* or *ampC* under the control of the P_rha_ promoter^81^ was achieved by replacing the P_ara_ promoter with P_rha_ promoter in pHERD26T plasmid^80^ and inserting the gene of interest downstream using Gibson assembly master mix (Table S1A and S1B).

### Individual strain growth and broth microdilution assays

Overnight bacterial cultures were sub-cultured in LB medium and grown to mid-log phase with OD_600_ ∼0.4-0.6 at 37°C. Cultures were diluted in LB to an inoculum of 5 x 10^5^ CFU/mL based on a conversion of OD_600_ = 1 equating to 1 x 10^9^ CFU/mL for gram-negative species and 2 x 10^8^ CFU/mL for *S. aureus*. 30 or 50 µL of cells in LB were dispensed to each well of a clear-bottom 384-well culture plate (Nunc) with or without test compounds in a 2-fold dilution dose series with 0.5% DMSO or water included as vehicle controls. Plates were incubated at 37°C without shaking overnight in a humidified box or with shaking in a humidity cassette in a Spark 10M microplate reader (Tecan). OD_600_ measurements were made either as an endpoint or every 30 minutes or 1 hour with the Spark 10M microplate reader. Normalized growth was calculated as 100 x [(OD_600_ of positive control – OD_600_ of test compound)/ (OD_600_ of positive control – OD_600_ of vehicle control)] and plotted against dose response of the compound of interest.

### Mutliplex screen optimization and setup

Overnight starter cultures of all strains and species were grown individually in LB medium at 37°C for 16 h while shaking and were then sub-cultured 1:100 and grown at 37°C while shaking until OD_600_ = 0.2-0.7. To determine if sequencing reads correlated to cell density, each of the five barcoded bacterial species were grown to mid-log phase and density adjusted to an OD_600_ of 1, pooled evenly, and diluted for a range of OD_600_ 0.05-1, and Illumina libraries prepared as outlined below. To determine the optimum incubation period for the multiplexed chemical screen, barcoded strains were pooled evenly to a standard starting inoculum of 5×10^5^ CFU/mL^82^ using the conversion factors described above and grown over 24 hours with sample collection every 4 hours. All strains and species grew optimally between 0-8 hours with an increase in read counts of >2 log_10_ for most species and strains, at which point the sequencing read counts for all strains began to reach a plateau (Figure S2A). *E. coli* and *K. pneumoniae* had the highest read counts after 4 and 8 hours, consistent with their relatively quick growth rate, while *A. baummanii*, *P. aeruginosa*, and most *P. aeruginosa* depletion strains grew at equivalent rates. The two growth defective depletion strains, *bamE*-hypo and *lptD*-hypo had significantly reduced sequencing reads compared to wildtype *P. aeruginosa* PA14 strain and only increased by ∼1 log_10_ from the starting inoculum. Likewise, of the 5 species, *S. aureus* was the slowest growing strain. We selected 12 hours as the end point for the chemical screen since read counts and therefore, growth, had plateaued for all strains (Figure S2A). Ciprofloxacin hydrochloride (MP Biomedicals) at 8 µM, trimethoprim at 32 µM and POL7080 (Anaspec) at 8 nM and 64 nM were added as positive controls on each plate in 24 replicates to measure Z’ factor and detect target-specific hypersensitization. In the high-throughput chemical screen the pooled inoculum was prepared at a final concentration of 5 x 10^5^ CFU/mL, with an increased proportion of *S. aureus* and the *P. aeruginosa lptD*-and *bamE*-hypomorph strains (30X, 30X, and 15X, respectively), with the goal of obtaining approximately 100 reads per well per strain. 30 µL of the cell pool was dispensed to each well of a clear-bottom 384-well culture plate (Nunc) with or without inhibition compounds, and cells were growth at 37 °C. DMSO and ciprofloxacin hydrochloride (MP Biomedicals) at 10 µM were added as positive and negative controls on each plate in 16 replicates each. After 12 hours (or otherwise indicated in preliminary experiments), plates were sealed, heated at 80°C for 2 h, and frozen and stored at −80°C.

### Library construction

Plates were thawed, and 30 µL of 20% DMSO was added to each well, mixed, and 20 µL was transferred to a 384-well PCR plate (Eppendorf) before heating for 10 minutes at 98C. 1 µL of the lysate was transferred to a new 384-well PCR plate containing a 9 µL of a PCR mix consisting of Q5 Hot Start HiFi 2X DNA polymerase master mix, 500 nM of each the forward and reverse barcoded primers (Table S2C), and a total of 1 ng of purified gDNA from two uniquely barcoded *P. aeruginosa* wild type strains. 18 PCR cycles were performed, followed by pooling all PCR reactions and column purifying the product (Zymo Research). PCR products were 5’ phosphorylated with T4 polynucleotide kinase, followed by column purification, and the subsequent addition of a 3’ A-tail using Klenow Fragment (3’-5’ exo^-^). After an additional column purification of the prepared PCR products, an Illumina adapter was added by ligation using a Blunt/TA ligation master mix. The Illumina adapter was first constructed by heating the primers P5 adapter and P7 adapter (Table S2B) at briefly at 95°C and allowing to cool to 2°C at a rate of 1°C/minute in a thermocycler. The ligation reaction was purified using AMPureXP SPRI magnetic beads (Beckman Coulter) and PCR amplified using P5-amp and P7-amp primers for 4 cycles. The final library was purified with SPRI beads and quantified using the Agilent Tapestation DS1000 high sensitivity. Samples were sequenced using an Illumina MiSeq for preliminary experiments or an Illumina HiSeq for the 50,000-compound screen. Reads were demultiplexed and tabulated using custom scripts that can be found on [GitHub](https://github.com/broadinstitute/psa_brd1401_manuscript).. For assay development (Figure 2), reads in each well were normalized to an internal PCR control barcode sequence using the formula: Normalized reads = Raw reads/Control reads*10000. For determining Z’-factors, reads were log transformed and Z’-factors were calculated for each strain and each plate using the formula: Z’-factor = 1 – 3(α_CIP_ + α_DMSO_)/ (|μ_CIP_ – μ_DMSO_|).

### Multiplexed high-throughput screen analysis

For the multiplexed screen, 53,958 compounds were included for analysis. Compounds were excluded from analysis if they had fewer than two replicates, some of which were removed due to having no read counts corresponding to barcodes from two uniquely barcoded *P. aeruginosa* wild type gDNA that were spiked in during PCR amplification of the library, as described above. In addition, compounds in wells with known technical error were also excluded. This gave a final list of 13,399 compounds from the DOS library, 30,541 compounds from MLPCN and 9,989 compounds from the Wuxi library. Compounds that suppressed the growth of the entire strain pool were determined as wells where the total count of all strains in the well was less than 10% of the median count in DMSO wells on the same screening plate, in both replicates. For all other compounds, the sensitivity of each strain to each compound in the multiplexed screen was quantified by calculating a standardized log fraction (SLF) for a particular strain *s* and compound *c* in a given well as follows:

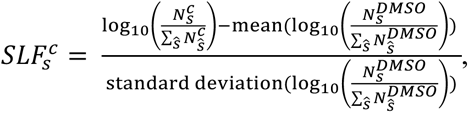

where 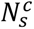, represents the read counts for a specific strain *s* in a given compound *c* in a single well. Note that a pseudocount of one was added to the read counts of each strain prior to the calculation of total well count and SLF. Also, each strain’s fractional count per well is standardized relative to its average fractional count in the twelve DMSO control wells on its same plate to account for plate-to-plate variability. Taken together, a negative SLF value indicates that the strain’s fractional growth in the compound is inhibited relative to its fractional growth in DMSO. Validation of compound activity by a non-pooled, absorbance-based assay yielded a measure of percent growth inhibition, which was calculated for each strain in each compound *c* as follows:

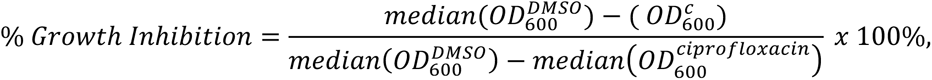

where DMSO and ciprofloxacin were used as controls. Because negative values indicate no growth inhibition, any percent growth inhibition values that were less than −20% were capped at −20%. Scripts for calculating SLF can be found on [GitHub](https://github.com/broadinstitute/psa_brd1401_manuscript).

### Proteomic quantification of OprL depletion

Triplicate mid-log cultures of wildtype PA14 and the *oprL*-hypomorph strain were collected by centrifugation at 5500g for 5 minutes, washed 3-times with phosphate buffer saline, and resuspended in 50mM NH_4_HCO_3_ + 8M Urea. 100mg of acid-washed 0.1mm glass disruption beads were added, and cells were disrupted with a TissueLyse LT (Qiagen) for 5 mins at 50Hz, followed by centrifugation for 20 minutes at 16,000g. Supernatant was collected, and protein concentration of each sample was determined by Qubit fluorometry (Invitrogen). 10μg of each sample was separated by SDS-PAGE using a 10% Bis-Tris NuPAGE gel (Invitrogen) with the MES buffer system. The 1 cM mobility region was excised and processed by in-gel digestion with trypsin using a ProGest robot (Digilab). One fifth of each digested sample was analyzed by nano UPLC-PRM/MS with a Proxeon EASY-nLC 1000 HPLC system interfaced to a ThermoFisher Q Exactive HF mass spectrometer. Peptides were loaded on a trapping column and eluted over a 75μm x 50cm analytical column (Thermo Fisher P/N ES-803) at 300 nL/min using a 1hr reverse phase gradient; both columns were packed with PepMap RSLC C18, 2 μm resin (Thermo Scientific). The mass spectrometer was operated in Parallel Reaction Monitoring (PRM) mode with the Orbitrap operating at 60,000 FWHM and 17,500 FWHM for MS and MS/MS respectively. Data were collected for the target peptides and analysis was performed using the XCalibur QualBrowser software (ThermoFisher).

### OM permeability assay

The OM permeability of *P. aeruginosa* depletion strains was assessed using a chromogenic substrate, nitrocefin (EMD Millipore), and its beta-lactamase activator, AmpC, from *P. aeruginosa* based on the assay developed by Angus *et al*.^83^ *P. aeruginosa* strains with an episomal copy of *ampC* gene under rhamnose-inducible (P_rha_) promoter were grown overnight at 37 °C in LB broth and then sub-cultured with 0.1% rhamnose to induce AmpC expression. When OD_600_ reached 0.5-0.6, the culture was spun down, and the pellet was gently resuspended in 10 mM Na-HEPES at pH 7.0 with 5 mM MgCl_2_ (assay buffer). The OD_600nm_ was measured and adjusted to 1.0. 400 μL of the adjusted cell culture was dispensed in a 96 well deep-well block plate, where each well contained 0.1% rhamnose. The plate was incubated at 37°C under shaking conditions for 1 hour to allow equilibration of *ampC* expression. 200 μL of the incubated samples were spun at 6,000 g for 10 minutes. 50 μL of the supernatant (SN), which contains extracellular AmpC, was transferred over to a storage plate on ice. The bacterial pellet was resuspended in equal volume of assay buffer with 10 μM colistin (Adipogen) and incubated at 37 °C for 90 minutes to lyse bacteria. The lysate (L), which contains intracellular AmpC, was spun down and 50 μL of the supernatant was collected and transferred to the chilled storage plate. 20 μL of the previously collected supernatant and the lysate samples was transferred to a 96-well flat-bottomed falcon plate (Corning) and 0.1 mg/mL nitrocefin in assay buffer was added to the samples for a final nitrocefin concentration of 0.087 mg/mL. Absorbance at 495nm was measured for 12 hours at 10-minute intervals in a Spark 10M microplate reader (Tecan). A standard curve of purified AmpC (Creative Biomart) was used to calibrate and quantify the presence of beta-lactamase in various tested strains. The concentration for standard curve ranged from 0.125 μg/mL to 0.000488 μg/mL. Relative OM permeability was quantified with the following formula: OM permeability = (Absorbance of SN) / (Absorbance of SN + Absorbance of L).

### Protein purification

*P. aeruginosa* TolB and OprL were expressed and purified using *E. coli* expression systems. An N-terminal 6X-His tag followed by a TEV protease cleavage site and then an AVI-tag preceded the TolB sequence lacking the signal peptide region (22-432 AA) in a pET-based expression vector. After induction, cell pellet was lysed by sonication in 20 mM Tris at pH 8.0 with 0.5 mM PMSF and 500 mM NaCl. The lysate was centrifuged to remove cell debris, the supernatant was loaded on a Ni-NTA column and washed with 20 mM Tris at pH 8.0 with 500 mM NaCl and 20 mM or 50 mM imidazole sequentially. Protein was eluted with 20 mM Tris at pH 8.0 with 500 mM NaCl and 300 mM imidazole. The His-tag was then cleaved with a His-tagged TEV protease and the protease removed using another round of Ni-NTA column run. The flow through was then incubated with His-tagged BirA protein to add biotin tag followed by another round of Ni-NTA column to remove the BirA protein. The final biotinylated TolB protein was run through a S75 (Cytiva) size exclusion chromatography column in 20 mM Tris at pH 8.0 with 500 mM NaCl. Protein purity was assessed on an SDS gel and confirmed by mass spectrometry (Figure S6A). OprL was purified using the same strategy with minor modifications. An OprL sequence without signal sequence (50-168 AA) and with an N-terminal 6XHis-TEV cleavage site-AVI tag was purified as a biotinylated protein in 20 mM Tris at pH 8.0 with 200 mM NaCl following the above protocol. Mass spectrometry and sequence analysis revealed a point mutation at T137Q in OprL. StrepII-OprH was purified from the *ΔoprH/pRha-StrepII-oprH P. aeruginosa* strain (PA30 in Table S1G) in which the native copy of OprH was deleted and a StrepII-tagged OprH was episomally over-expressed under the control of a rhamnose-inducible promoter. Log phase culture of this strain was diluted and induced with 0.125% rhamnose and grown up to an OD_600_ of 1-1.2 in LB at 37°C with shaking. The culture was spun down at 6000g for 20 minutes at 4°C and pellet suspended in 50 mM Tris-HCl at pH 8.0, 150 mM NaCl, 0.5 mg/mL lysozyme (Sigma) and protease inhibitor cocktail (Roche). Cells were lysed with the Microfluidizer LM20 (Microfluidics) cell disruptor and lysate was incubated with 1% N-Dodecyl-β-D-maltoside (Anatrace) for 1 1hour at 4°C with rocking. The lysate was centrifuged at 10,000 g for 20 minutes to remove cell debris. Supernatant was added to Strep-Tactin®XT 4Flow®resin (IBA Life Sciences) and incubated for 1 hour at 4°C with rocking. The beads were spun down, supernatant discarded and washed in 5X volume of 50 mM Tris-HCl at pH 8.0, 150 mM NaCl, and 1% N-Dodecyl-β-D-maltoside. Elution was done with 3X bead volume of BXT buffer (IBA Life Sciences) with 1% N-Dodecyl-β-D-maltoside. Elute was then run on the Superdex®200 increase 10/300 GL (Cytiva) column in 20mM HEPES at pH 8.0, 50mM KCl, and 0.05% N-Dodecyl-β-D-maltoside. The fractions were then concentrated on the Strep-Tactin®XT 4Flow®resin using the method described above and eluted in 3X volume of the SEC buffer with 50 mM biotin. The sample was then dialyzed thrice for 6 hours each against 50 mM HEPES at pH 7.4, 150mM NaCl, 1 mM EDTA and 0.05% N-Dodecyl-β-D-maltoside using a 35 KDa MWCO dialysis cassette (ThermoScientific). The protein samples were then snap-frozen and stored at −80 °C. Samples were run on a 12%TGX (Biorad) gels with and without boiling at 100°C for 10 minutes. Protein purity and mass was assessed with intact mass spectrometry (Waters BioAccord LC-MS) and cooperative unfolding was detected using a differential scanning calorimeter to estimate the melting temperature in DDM micelles. 900 µg of purified StrepII-OprH was diluted to 400 µL with storage buffer (20 mM HEPES 7.4, 150 mM NaCl, 50 mM Biotin, 0.05% DDM), loaded into sample plates (500 µL V-bottom round 96-w plates; VWR) and sealed with Silicone covers. The temperature ramp for each injected sample was 10-110C at 0.5°C /min followed by a standard cleaning cycle using Contrad 70 and water injections (Malvern Instruments). Collected data was analyzed using the instrument’s software by subtracting the buffer only blanks.

### DSF

Recombinant proteins (5-30 µg) were incubated with 200 µM BRD-1401 or DMSO (2%) in 1x PBS. SYPRO™ Orange Protein Gel Stain (ThermoFisher S6650) was added to each reaction for a final concentration of 2x. The dye signal was monitored in a Roche Lightcycler 480 Real-Time PCR System as a function of a temperature ramp (25-95°C; 0.1°/s) and the Tm was calculated using the instrument’s software.

### NMR

^1^H-NMR data was recorded on a Bruker 600MHz spectrometer equipped with a cryogenic QCIF probe and an automatic sample handling system. STD-NMR and CPMG experiments were conducted at 280K and processed using the spectrometer’s TopSpin 3.6 software. Intensities of the most prominent aromatic proton peaks were used to calculate %STD = 100 x [(I_ref_ – I_sat_) / I_ref_] where *ref* and *sat* are the reference and saturated spectra, respectively. Carr-Purcell-Meiboom-Gill (CPMG)-T2 experiments were recorded as a series of ^1^H-NMR spectra with 1-600 CPMG blocks of 1ms duration. Normalized peak intensities were fitted to an exponential decay function to calculate the transverse relaxation rate R_2_. Samples typically contained 15 µM protein and 500 µM compounds in NMR buffer (50 mM HEPES pH 7.4, 150 mM NaCl, 0.05% DDM in a deuterated background). All molecules tested in this study (except analog IA-1) exhibited mild STD in buffer at 500 µM as calculated by the above formula under standard experimental conditions. To normalize for this, we subtracted the background signal for each molecule from the calculated STD in the presence of protein (ΔSTD = STD_protein_ – STD_buffer_).

### Biolayer Interferometry

Biotinylated Avi-tagged TolB was diluted in binding buffer (50mM HEPES pH 7.4, 150mM NaCl, 0.05% Tween-20) and immobilized on Streptavidin biosensors using an Octet Red 384 instrument (ForteBio). The Sensors were subsequently blocked with 100 µM Biotin and immersed in binding buffer to reach a stable baseline (T=0). Samples containing 10 µM OprL or 10 µM OprL + 500 µM BRD-1401 were analyzed in duplicates; the dissociation phase were fit to a single exponential function to deduce the off rate for each condition.

### Whole genome resequencing and SNP identification

Genomic DNA was extracted from stationary phase culture using DNeasy® DNA extraction kit (Qiagen) as per the manufacturer’s instructions. DNA was quantified with Qubit dsDNA HS Kit (Invitrogen) and diluted to 0.4 ng/µL. Libraries were prepared with the Nextera XT DNA library preparation kit (Illumina) as per standard protocol. Briefly, 1 ng of genomic DNA was fragmented and tagged with adaptor sequences using the TD and ATM buffers. The DNA is then amplified using i7 and i5 primers and NPM buffer using a limited-cycle PCR. Prepared libraries were cleaned up with AMPure XP beads (Beckman Coulter) and quantified with Qubit™ dsDNA HS Kit (Invitrogen). All samples were pooled to a final concentration of 5-10 ng/µL and sequenced on a NextSeq 500 (Illumina). The demultiplexed data was assembled and SNPs detected with Pilon software using reference genome sequences.^84^

### Transcriptomics with RNA sequencing (RNA-seq)

Bacterial cultures were inoculated from glycerol stocks, grown overnight with appropriate selection antibiotics at 37°C with shaking and sub-cultured at 37°C with shaking until log-phase growth. Cultures were diluted to an OD_600_ of 0.2-0.4. LB medium with appropriate drugs such as BRD1401 or inducers such as arabinose was prepared at 2X final concentration and dispensed into 384 well plates (Nunc). Diluted bacterial suspensions were added on top to achieve a final bacterial density of 0.1-0.2 OD_600nm_ and incubated at 37°C. At desired timepoints the bacterial suspension was added to 1X lysis buffer (BLUE buffer; Microgem) with RNAgem (Microgem) in 96w PCR plates and incubated at 72°C for 10 minutes to lyse bacteria. Samples were immediately transferred to and stored at −80°C until further processing. RNA was extracted with Direct-Zol 96 well plate kit (Zymo Research) and eluted into DNase/RNase-free water. Illumina cDNA libraries were generated using a modified version of the RNAtag-seq protocol.^85^ Briefly, 250ng of extracted RNA was fragmented, dephosphorylated, and ligated to DNA adapters carrying 5’-AN_8_-3’ barcodes of known sequence with a 5’ phosphate and a 3’ blocking group. Barcoded RNAs were pooled and depleted of rRNA using the Pan-Bacteria riboPOOL depletion kit (siTOOLs Biotech, Galen Laboratories). Pools of barcoded RNAs were converted to Illumina cDNA libraries in 2 main steps: (i) reverse transcription of the RNA using a primer designed to the constant region of the barcoded adaptor with addition of an adapter to the 3’ end of the cDNA by template switching using SMARTScribe (Clontech) as described;^86^ (ii) PCR amplification using primers whose 5’ ends target the constant regions of the 3’ or 5’ adaptors and whose 3’ ends contain the full Illumina P5 or P7 sequences. cDNA libraries were sequenced on the Illumina [NovaSeq SP 100] platform to generate paired end reads. Sequencing reads from each sample in a pool were demultiplexed based on their associated barcode sequence using custom scripts. Up to 1 mismatch in the barcode was allowed provided it did not make assignment of the read to a different barcode possible. Barcode sequences were removed from the first read as were terminal G’s from the second read that may have been added by SMARTScribe during template switching. Reads were aligned to NC_0088463.1 (UCBPP-PA14) using BWA^87^ and read counts were assigned to genes and other genomic features using custom scripts. The dataset was analyzed using the DESeq2 (1.28.1) package in R to obtain fold changes in gene expression across multiple conditions.^88^ Two separate RNA-seq datasets were generated and their accession numbers on the NCBI-Geo repository are GSE252756 (dataset 1) and GSE254108 (dataset 2). In the first experiment linked to dataset 1, strains were exposed to 125 µM of BRD1401 or 0.5% DMSO vehicle control for 90 minutes while in the second experiment linked to dataset 2, strains were exposed to 256 µM of BRD1401 or 0.5% DMSO vehicle control for 120 minutes. Read counts for each gene after each treatment is tabulated in Tables S5A (dataset 1) and S5B (dataset 2).

### Epithelial cell infection assay

Human bladder epithelial cell line 5637 (RRID: CVCL_0126) were cultured and maintained in RPMI medium (Gibco; ThermoFisher Scientific) supplemented with 10% heat inactivated FBS (Sigma) at 37°C with 5% CO_2_. Bladder epithelial cells were infected with *P. aeruginosa* wildtype PAO1 strain as previously described.^57^ Briefly, ∼100,000 human bladder epithelial cells (RRID: CVCL_0126) were seeded in 24-well plates overnight in RPMI (Gibco; ThermoFisher Scientific) supplemented with 10% heat inactivated FBS. Log-phase *P. aeruginosa* PAO1 grown in LB was used to infect at a multiplicity of infection of 10. Cells were co-incubated with bacteria for 2 hours, media removed and fresh media containing 300 μg/ml gentamicin was added for an additional 2 hours to kill extracellular bacteria. Dimethyl sulfoxide vehicle control or 50 or 100 μM BRD1401 was then added to wells (t = 3 h). At each timepoint (t = 3, 24, 48 h), epithelial cells were washed twice with phosphate buffered saline, lysed with 0.1% Triton-X100 and the lysate bacteria plated on LB agar plates. Agar plates were incubated at 37°C overnight, and colony forming units (CFU) enumerated manually. CFU normalized to the vehicle control was plotted as %survival.

### Intra-bacterial compound accumulation assay

Bacterial cultures were inoculated from glycerol stocks into LB, grown overnight with appropriate selection antibiotics and inducers at 37°C with shaking. These cultures were diluted to an OD_600nm_ of 0.05 in LB and grown at 37°C with shaking to log phase. Arabinose was added at 0.063% to deplete OprL in strains with an arabinose-inducible copy of OprL under the control of the P_araBAD_ promoter, while rhamnose was added at 0.125% or 0% for strains with an episomal copy of OprH under the control of rhamnose-inducible P_rhaBAD_ promoter. Cultures were processed, exposed to compounds and analyzed as described by Geddes *et al*.^89^ Briefly, 50 mL of culture at an OD of ∼0.55 was spun down at 5000g for 10 minutes and the supernatant was discarded. The bacterial pellet was washed once with phosphate buffer saline, spun down and suspended in 2.2 mL of phosphate buffer saline. 0.875 mL of this suspension was aliquoted into a 96w deep well block, acclimatized to 37°C for 10-20 minutes and then exposed to 100 µM of BRD1401 or 50 µM of ciprofloxacin for 10 or 40 minutes at 37°C. 0.8 mL of this bacterial was layered on 0.7 mL of a frozen mix of AR20 silicone oil (9 parts) and high temperature silicone oil (1 part) and spun at 13,000g for 2 minutes. Supernatant was removed and the bacterial pellet suspended in 0.2 mL of ultrapure water (ThermoFisher). Lysis was achieved by repeated freeze-thaw cycles followed by a spin at 13,000g for 2 minutes and a wash with methanol (ThermoFisher) to collect the supernatant fraction as lysate. This lysate was run on the Acquity liquid chromatography system (Waters) followed by the Triple Quad^TM^ 4500 MS/MS (SCIEX) to detect compounds. Data analysis was performed using Analyst software (SCIEX) and quantification was based on a standard curve generated with the same system.

### Trypsin digestion

3 µg of purified OprH (16 µM) was pre-incubated with Kdo2-LipidA (Avanti Polar Lipid) and each of the the compounds in 50mM Tris-HCl at pH 7.5 with 150mM and 4% DMSO (vehicle control) under 37°C for 30min. 42 µM trypsin from the bovine pancreas (Sigma) was added to the mixture and incubated for 5 additional hours at37°C. The samples were boiled or not boiled with SDS-PAGE sample buffer (BIO-RAD) at 100°C for 15min, then run in a 15% Tris-glycine SDS-PAGE gel for 42 minutes at 150V until the dye head reached the bottom. Gel was fixed with 40% methanol and 10% acetic acid for 30 minutes, stained with Bio-Safe Coomassie G-250 (BIO-RAD) for 30 minutes and then de-stained with water. Gels were imaged on Gbox Chemi XT4 (Syngene) and mean band intensity quantified with Fiji (ImageJ).^90^

### LPS and protein detection

Bacterial cultures were inoculated from glycerol stocks, grown overnight with appropriate selection antibiotics and inducers at 37°C with shaking and sub-cultured at 37°C with inducer molecules to obtain a log-phase culture. Rhamnose was added to the sub-culture at 0.25% for strains with StrepII-tagged OprH under the control of rhamnose-inducible P_rhaBAD_ promoter. To measure impact of BRD1401 on LPS-OprH interaction in the *oprL*-KD strain (PA32 in Table S1G), log phase cultures were diluted to an OD of 0.0005 and incubated overnight at 37°C with shaking with 0.063% arabinose, 0.25% rhamnose and BRD1401 until an OD of 0.4-0.6 was reached. Cultures were spun down at 5000g for 10 minutes and supernatant discarded. Cell pellets were lysed by sonication in 50 mM Tris-HCl at pH 7.5, 150 mM NaCl, 1 mM EDTA, 0.5 mg/mL lysozyme, 5 µg/mL DNAse and protease inhibitors (Roche). Lysate was incubated with 1% N-Dodecyl-β-D-maltoside for 1 hour at 4°C with mixing and then spun down at 20,000g for 20 minutes at 4°C to remove cell debris. Supernatant was incubated with Strep-Tactin®XT 4Flow®resin resin for 15-30 minutes, washed with 10X volume of 100 mM Tris-HCl at pH 8.0, 150 mM NaCl and 1 mM EDTA and eluted with 3X volume of BXT buffer (IBA Life Sciences). Samples were run on 4-20% TGX gels (Biorad) and blotted with the LPS antibody against *P. aeruginosa* PA14 antigen O10 (MEDIMABS). Blots were stripped and probed with rabbit anti-StrepII antibody (Abcam) to detect Strep-tagged OprH. Images were quantified with Fiji (ImageJ) software.^90^ LPS band intensity was normalized to StrepII-OprH protein band intensity for each sample and expressed as a ratio. This ratio was normalized to vehicle (0.5% DMSO) treated sample and plotted.

### Co-immunoprecipitation

Bacterial strains with FLAG-tagged OprL and StrepII-tagged OprH or untagged OprH (PA36 and PA39 in Table S1G) were used. To measure specific binding of OprL-FLAG to StrepII-OprH, PA39 was inoculated with 0.5% arabinose to induce FLAG-tagged OprL and then sub-cultured with 0.5% arabinose and varying rhamnose concentrations to induce expression of untagged OprH until an OD_600nm_ of 0.5-0.8. Cultures were then spun down at 5000g for 10 minutes and supernatant discarded. Cell pellets were suspended in 50 mM Tris-HCl at pH 8.0, 150 mM NaCl, 1mM EDTA, 0.5 mg/mL lysozyme, 5 µg/mL DNAse and protease inhibitors (Roche) and lysed by sonication. Lysate was incubated with 0.05% N-Dodecyl-β-D-maltoside (Anatrace) for 1 hour at 4°C with rocking and then spun down at 20,000g for 20 minutes at 4°C to remove cell debris. Supernatant was incubated with Strep-Tactin®XT 4Flow®resin (IBA Life Sciences) resin for 30 minutes at 4°C, washed with 10X volume of 50 mM Tris-HCl at pH 8.0, 150 mM NaCl, 1mM EDTA and 2 mM biotin) and eluted with 3X volume of BXT buffer (IBA Life Sciences). Samples were run on 12% TGX gels (Biorad) and blotted with the anti-FLAG M2 antibody (Sigma) and rabbit anti-StrepII antibody to detect FLAG-tagged OprL and StrepII-tagged OprH, respectively. The protein band intensities were quantified with Fiji and the OprL-FLAG band intensity in the elution samples, first normalized to StrepII-OprH band intensity in the same sample and then further normalized to the control sample (0% rhamnose) was plotted. To measure impact of BRD1401 exposure, PA36 was inoculated with 0.5% arabinose to induce OprL-FLAG and then sub-cultured until log-phase with 0.063% arabinose to downregulate OprL-FLAG expression and 0.13% rhamnose to induce StrepII-OprH expression. It was then diluted to an OD_600nm_ of 0.001 and exposed overnight to varying doses of BRD1401 or DMSO vehicle control. StrepII-OprH was pulled down as described above and the protein band intensities in elution samples quantified with ImageJ (Fiji). The ratio of OprL-FLAG to StrepII-OprH was calculated as described above with normalization to vehicle DMSO control and plotted.

### Ethidium bromide accumulation

Membrane permeability was quantified by measuring uptake of ethidium bromide whose fluorescence emission at 585 nm increases upon binding intracellular DNA. Log phase culture of *oprL*-hypomorph was spun down and suspended in phosphate buffered saline. Bacterial cell density was then adjusted to OD_600nm_ of 1.0 and 40 µL of this bacterial suspension was mixed with 10 µL of 50 µM (10X final concentration) of ethidium bromide solution in a 96 well back-walled plate. Ethidium bromide uptake was allowed with shaking at 37°C, and uptake was measured by fluorescence emission of DNA-bound ethidium bromide at 585 nm after excitation at 530 nm with the Spark 10M microplate reader (Tecan). After 10-15 minutes 50 µL of colistin or BRD1401 at 2X the final concentration was added in a dose response to the wells and fluorescence read in 1-minute intervals for an hour. Relative Fluorescence Units (RFU) is plotted against time.

### Membrane fluidity

Membrane fluidity of *P. aeruginosa* strains was measured using the Membrane Fluidity kit (Abcam) as described previously.^60^ Lateral diffusion of lipophilic pyrene probe, included in the kit, in cell membrane leads to formation of transient homodimers or excimers whose emission shifts to higher wavelengths relative to the monomeric state. Thus, ratio of excimers to monomers, detected by differential fluorescence emission, is directly correlated to membrane fluidity. Log phase cultures of wildtype PA14, PA14*ΔoprH* and *oprL*-hypomorph strains grown in media with varying salt concentrations or doses of BRD1401 or ciprofloxacin were spun down and bacterial pellet suspended in phosphate buffered saline. To the bacterial suspension diluted to a density of 0.8 (OD_600nm_), 10 µM of the pyrenedecanoic acid, 0.08% F-127 and 1 mM EDTA was added. Samples were rocked for 30 minutes at room temperature to allow probe incorporation. Then samples were washed with 3-fold excess phosphate buffered saline and fluorescence emission measured at 470 and 405 nm after excitation at 350 nm with the Spark 10M microplate reader (Tecan). Ratio of relative fluorescence units at 470 nm versus 405 nm was normalized to the wildtype PA14 strain grown in LB medium with no salt (sodium chloride).

### Phylogenetic analysis of OprH homologs

The OprH sequence from PA14 (PA14_49200) was used as the query protein for a [PSI-BLAST](https://blast.ncbi.nlm.nih.gov) search. We performed 5 iterations using default parameters, except we increased the number of alignments view to 5000. After 5 iterations, the distant homolog OmpW from *E. coli* and other bacterial species appeared as hits. The tree was produced using NCBI BLAST pairwise alignment feature after the 5^th^ iteration and the [iTOL tree viewer](https://itol.embl.de/).

## Experimental Procedure for “Compound A” and “BRD1401 (revised)” synthesis

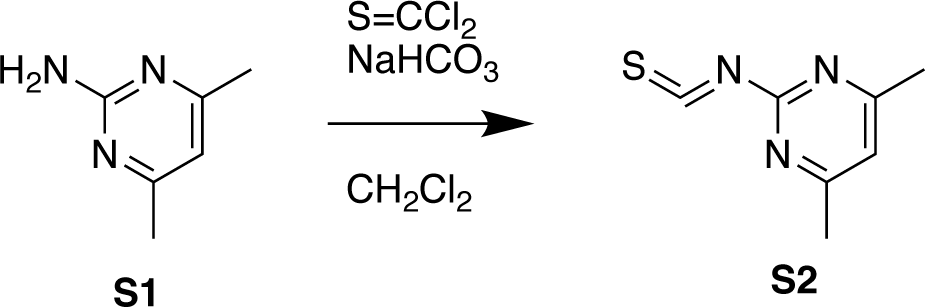

### 2-Isothiocyanato-4,6-dimethyl-pyrimidine S2

To a mixture of 4,6-dimethylpyrimidin-2-amine **S1** (15.0 g, 121 mmol) and NaHCO_3_ (25.5 g, 304 mmol, 11.8 mL) in DCM (300 mL) was added thiocarbonyl dichloride (14.0 g, 121 mmol, 9.34 mL) at 0 °C. Then the reaction mixture was stirred at 40 °C for 6 hrs. On completion, the reaction mixture was filtered, and the filtrate was concentrated in vacuo to get a residue. The residue was purified by column chromatography (Petroleum ether: ethyl acetate = 20: 1) to give the title compound **S2** (7.00 g, 34% yield) as a brown oil. ^1^H NMR (400MHz, CDCl_3_) δ = 6.93 (s, 1H), 2.47 (s, 6H).

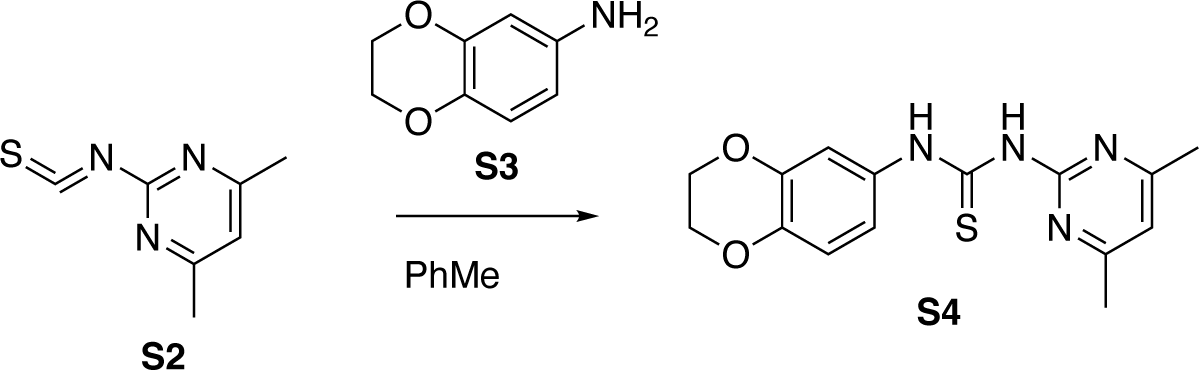

### 1-(2,3-Dihydro-1,4-benzodioxin-6-yl)-3-(4,6-dimethylpyrimidin-2-yl)thiourea S4

A mixture of 2-isothiocyanato-4,6-dimethyl-pyrimidine **S2** (7.00 g, 42.3 mmol) and 2,3-dihydro-1,4-benzodioxin-6-amine **S3** (6.40 g, 42.3 mmol) in toluene (70 mL) was stirred at 25 °C for 1 hr, and a white solid was precipitated. On completion, the reaction mixture was filtered and the solid was washed with MeOH (10 mL) to give the title compound **S4** (12.0 g, 89% yield) as a white solid. ^1^H NMR (400MHz, DMSO-*d_6_*) δ = 10.62 (s, 1H), 7.41 (d, *J* = 2.0 Hz, 1H), 7.04 (dd, *J* = 2.0, 8.8 Hz, 1H), 6.99 (s, 1H), 6.87 (d, *J* = 8.8 Hz, 1H), 4.25 (s, 4H), 2.41 (s, 6H).

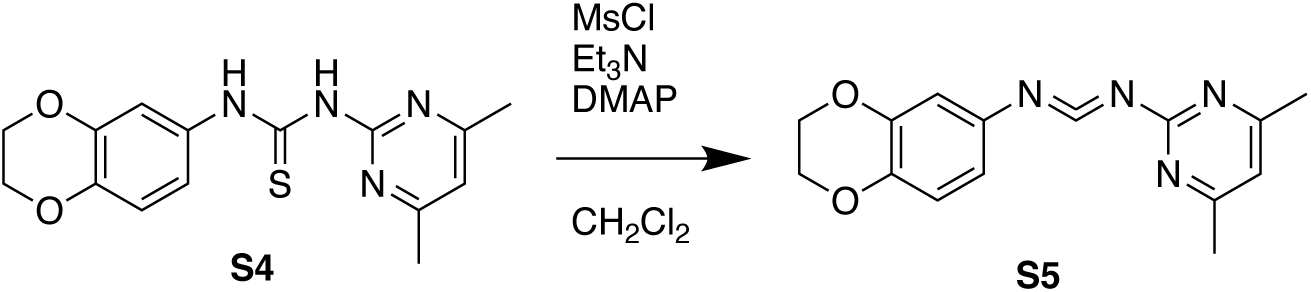

### *N*-(2,3-dihydrobenzo[*b*][1,4]dioxin-6-yl)-*N*-(4,6-dimethylpyrimidin-2-yl)methanediimine S5

To a solution of thiourea compound **S4** (400 mg, 1.3 mmol, 1 eq) in DCM (10 mL) was added TEA (384 mg, 3.8 mmol, 528 uL, 3 eq), DMAP (20 mg, 163 umol, 0.1 eq) and MsCl (290 mg, 2.5 mmol, 196 uL, 2 eq). The solution was stirred at 25 °C for 1 hr. TLC (PE/EtOAc = 3/1, Product (R_f_) = 0.50) indicated compound **S4** was consumed completely and one new spot formed. The crude product **S5** was used into the next step without any work up and further purification.

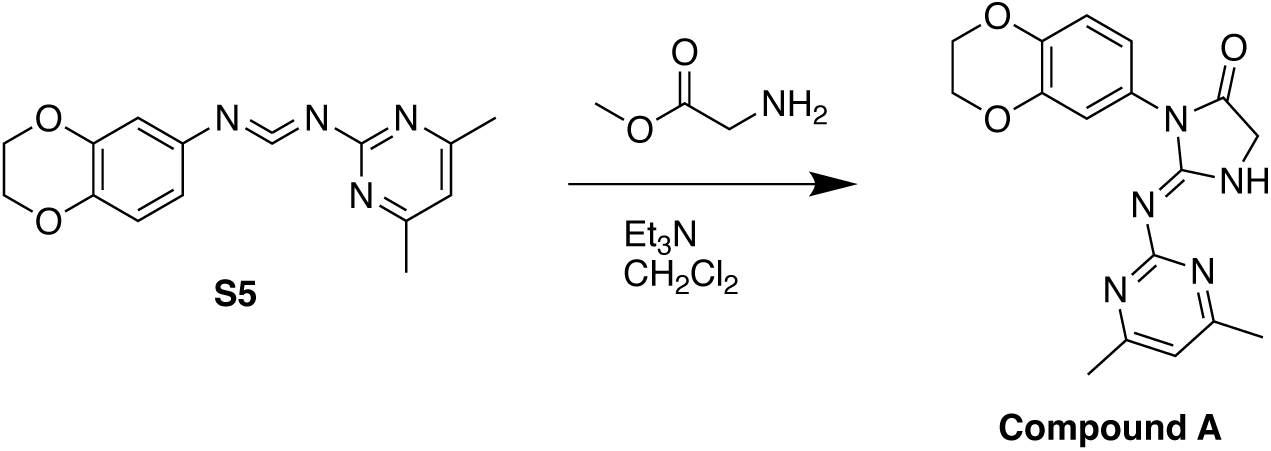

### 3-(2,3-dihydrobenzo[b][1,4]dioxin-6-yl)-2-((4,6-dimethylpyrimidin-2-yl)amino)-3,5-dihydro-4*H*-imidazol-4-one Compound A

To a solution of compound S5 (350 mg, 1.24 mmol, 1 *eq*) in DCM (10 mL) was added methyl 2-aminoacetate (166 mg, 1.86 mmol, 1.5 *eq*) and TEA (376 mg, 3.72 mmol, 518 uL, 3 *eq*). The solution was stirred at 25 °C for 6 hr. TLC (PE/EtOAc = 3/1, Product (R_f_) = 0.50) and LCMS (ET14092-83-P1A1) and HPLC (ET14092-83-P1B1) showed compound S5 was consumed completely. The reaction solution was diluted with water 50 mL and extracted with DCM 100 mL (50 mL x 2). The combined organic layers were washed with brine 60 mL, dried over Na_2_SO_4_, filtered and concentrated under reduced pressure to give a solid. The solid was purified by prep-TLC (SiO_2_, PE/EtOAc = 3/1) and prep-HPLC (column: Phenomenex Synergi C18 100×21.2mm, 4um; mobile phase: [water(0.1%TFA)-ACN];B%: 1%-33%,10min). Compound A (80 mg, 232 umol, 18.7% yield, 98.2% purity) was obtained as a white solid.

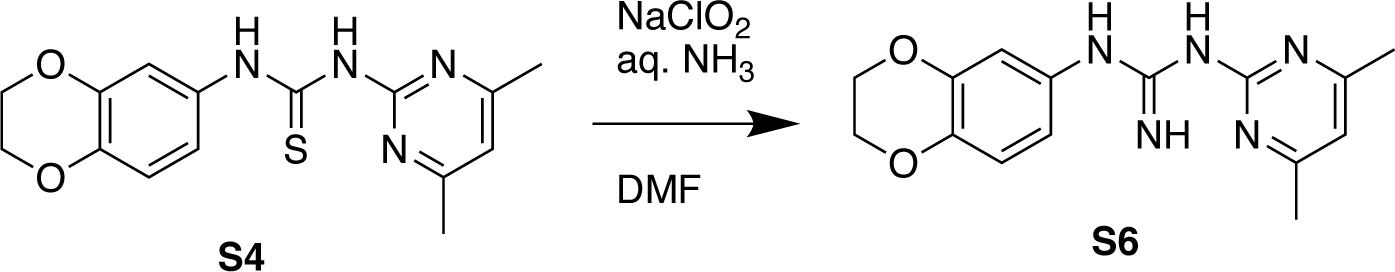

### 1-(2,3-Dihydro-1,4-benzodioxin-6-yl)-2-(4,6-dimethylpyrimidin-2-yl)guanidine S6

To a solution of 1-(2,3-dihydro-1,4-benzodioxin-6-yl)-3-(4,6-dimethylpyrimidin-2-yl)thiourea **S4** (2.00 g, 6.32 mmol) in DMF (40 mL) was added aqueous ammonia (2.46 g, 18.9 mmol, 2.71 mL, 27% purity) and sodium chlorite (1.72 g, 18.9 mmol). The mixture was stirred at 80 °C for 5 hrs. On completion, the reaction mixture was concentrated in vacuo. The getting residue was diluted with water (50 mL) and extracted with DCM (3 X 45 mL). The organic layer was dried over anhydrous Na_2_SO_4_, filtered and concentrated in vacuo. The getting residue was purified by column chromatography (dichloromethane: methane = 20: 1) to give the title compound **S6** (600 mg, 31% yield) as a light gray solid. ^1^H NMR (400MHz, DMSO-*d_6_*) δ = 7.95 (br s, 3H), 7.16 (s, 1H), 6.75 (s, 2H), 6.61 (s, 1H), 4.29 - 4.14 (m, 4H), 2.28 (s, 6H).

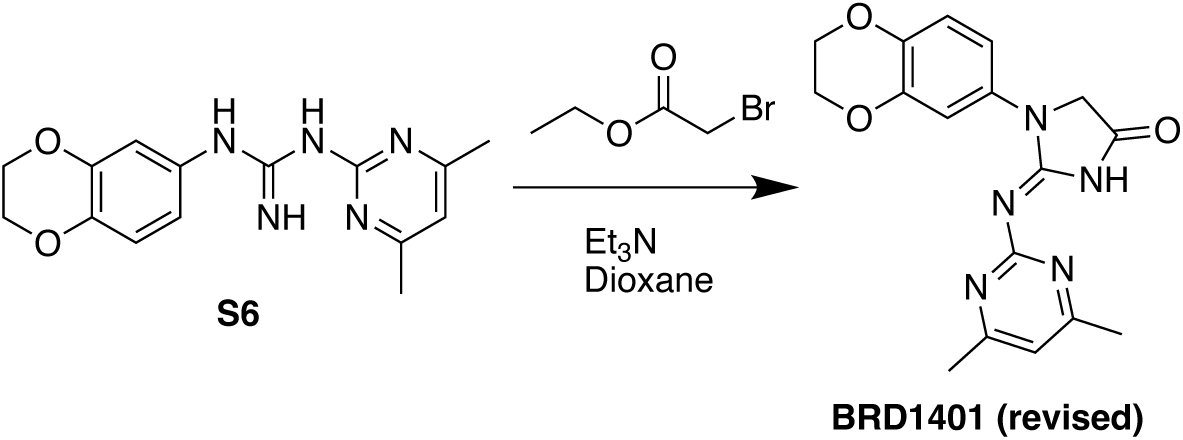

### (2E)-1-(2,3-dihydro-1,4-benzodioxin-6-yl)-2-(4,6-dimethylpyrimidin-2-yl)imino-imidazo-lidin-4-one (revised BRD1401)

To a solution of 1-(2,3-dihydro-1,4-benzodioxin-6-yl)-2-(4,6-dimethylpyrimidin-2-yl)guanidine **S6** (300 mg, 1.00 mmol) in dioxane (15 mL) was added ethyl 2-bromoacetate (251 mg, 1.50 mmol) and TEA (253 mg, 2.51 mmol, 348 uL). The reaction was warmed to 90 °C and stirred for 6 hrs. On completion, the reaction mixture was concentrated in vacuo. The getting residue was diluted with ice water (25 mL) and extracted with dichloromethane (3 × 15 mL). The organic layer was dried over anhydrous Na_2_SO_4_, filtered and concentrated in vacuo to get a residue. The residue was purified by Prep-HPLC (column: Phenomenex Gemini C18 250×50 mm, 10 um; mobile phase: [water(10mM NH_4_HCO_3_)-ACN]) to give the title compound **revised BRD1401** (116 mg, 33% yield) as a white solid. LCMS (M+1)^+^: 340.2. ^1^H NMR (400MHz, DMSO-*d_6_*) δ = 11.44 (br s, 1H), 7.45 (d, *J* = 2.4 Hz, 1H), 7.19 (dd, *J* = 2.4, 8.8 Hz, 1H), 6.87 (d, *J* = 8.8 Hz, 1H), 6.78 (s, 1H), 4.47 (s, 2H), 4.31 - 4.20 (m, 4H), 2.34 (s, 6H).

## Synthesis of analog IA-1

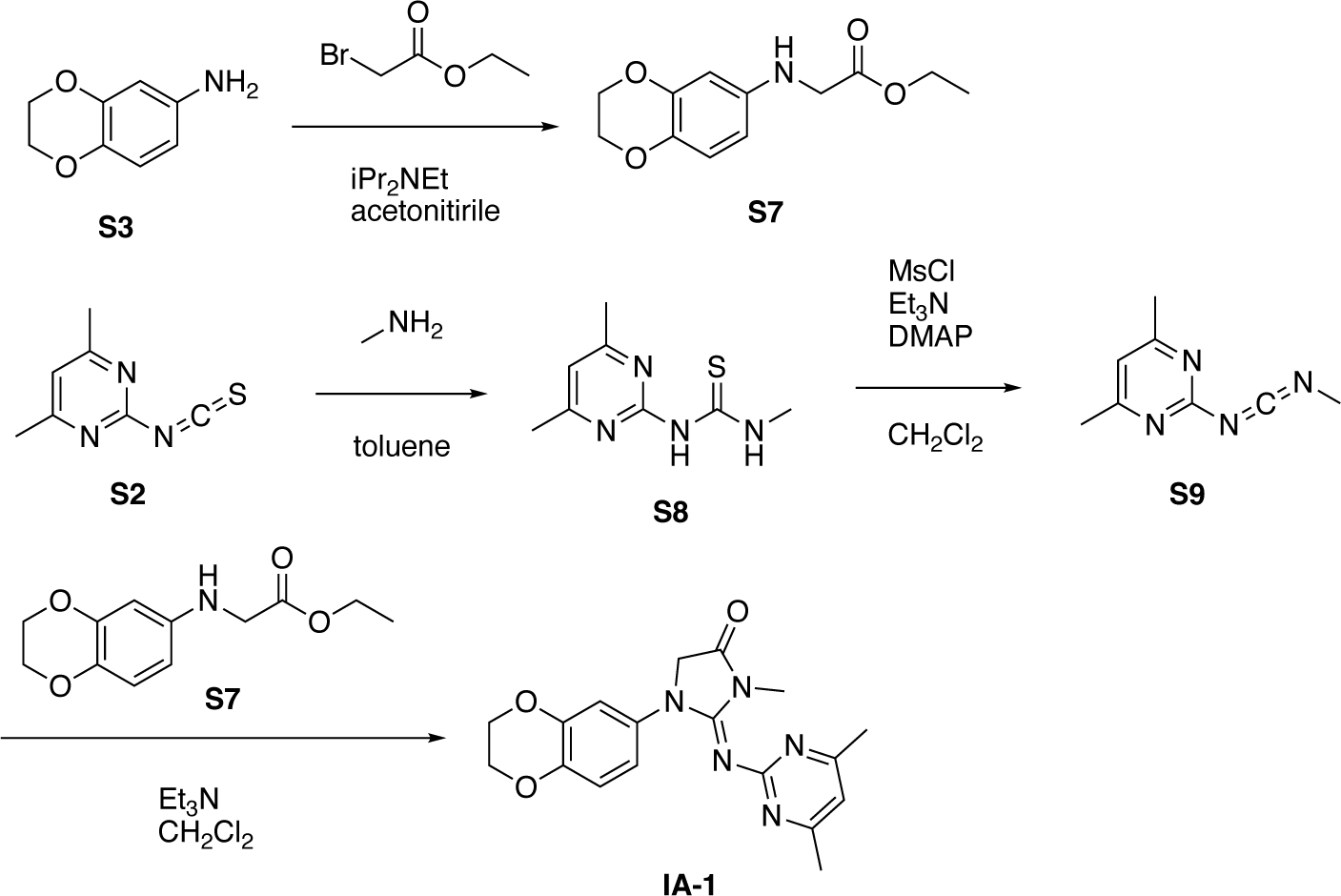

### Ethyl 2-(2,3-dihydro-1,4-benzodioxin-6-ylamino)acetate (S7)

To a solution of 2,3-dihydro-1,4-benzodioxin-6-amine **S3** (2.00 g, 13.2 mmol) and DIEA (2.56 g, 19.8 mmol) in acetonitrile (40 mL) was added ethyl 2-bromoacetate (2.21 g, 13.2 mmol) at 60 °C dropwise. Then the reaction mixture was stirred at 60 °C for 12 hrs. The reaction was concentrated in vacuo to give a residue. The residue was purified by silica gel column chromatography (Petroleum: ethyl acetate = 10: 1) to give the title compound **S7** (2.00 g, 63% yield) as a yellow gum. ^1^H NMR (400MHz, CDCl_3_) δ = 6.77 - 6.70 (m, 1H), 6.22 - 6.15 (m, 2H), 4.29 - 4.17 (m, 6H), 4.05 (br s, 1H), 3.84 (s, 2H), 1.31 (t, *J* = 7.2 Hz, 3H).

### 1-(4,6-Dimethylpyrimidin-2-yl)-3-methyl-thiourea (S8)

A mixture of 2-isothiocyanato-4,6-dimethyl-pyrimidine **S2** (1.40 g, 8.47 mmol), methylamine (2 M, 4.66 mL) in toluene (10 mL) was stirred at 110 °C for 12 hrs. The reaction was cooled to room temperature and a white solid was isolated. The solid was filtered and washed with (petroleum ether: ethyl acetate = 1:1, 10 mL) to give the title compound **S8** (1.10 g, 66% yield) as a white solid. ^1^H NMR (300MHz, DMSO-*d_6_*) δ = 11.26 (br s, 1H), 10.29 (br s, 1H), 6.92 (s, 1H), 3.10 (d, *J* = 4.4 Hz, 3H), 2.39 (s, 6H).

### *N*’-(4,6-dimethylpyrimidin-2-yl)-N-methyl-methanediimine (S9)

To a solution of 1-(4,6-dimethylpyrimidin-2-yl)-3-methyl-thiourea **S8** (300 mg, 1.53 mmol) in dioxane (15 mL) was added DMAP (18.6 mg, 152 umol) and TEA (464 mg, 4.59 mmol). And MsCl (350 mg, 3.06 mmol) was added dropwise. Then the mixture was stirred at 25 °C for 30 mins. The reaction mixture was filtered and the filtrate was concentrated in vacuo to give the title compound **S9** (240 mg, 96% yield), which was used in to the next step directly. LCMS (M+18+1)^+^: 181.2.

### (2E)-3-(2,3-dihydro-1,4-benzodioxin-6-yl)-2-(4,6-dimethylpyrimidin-2-yl)imino-1-methyl-imidazolidin-4-one (IA-1)

To a solution of *N*’-(4,6-dimethylpyrimidin-2-yl)-N-methyl-methanediimine **S9** (240 mg, 1.48 mmol) in dioxane (15 mL) was added triethylamine (449 mg, 4.44 mmol) and ethyl 2-(2,3-dihydro-1,4-benzodioxin-6-ylamino)acetate **S7** (351 mg, 1.48 mmol). The reaction mixture was stirred at 90 °C for 5 hrs. The reaction mixture was concentrated in vacuo to give a residue. The residue was purified by Prep-HPLC (column: Phenomenex Gemini 150×25mm, 10um; mobile phase: [water(0.04%NH_3_aq+ 10mM NH_4_HCO_3_)-ACN]) to give the title compound **IA-1** (49.3 mg, 9.2% yield) a white solid. LCMS (M+1)^+^: 353.9. ^1^H NMR (400MHz, DMSO-*d_6_*) δ = 6.60 - 6.51 (m, 3H), 6.48 (s, 1H), 4.44 (s, 2H), 4.16 - 4.08 (m, 4H), 3.01 (s, 3H), 2.09 (s, 6H).

## Synthesis of analog IA-2

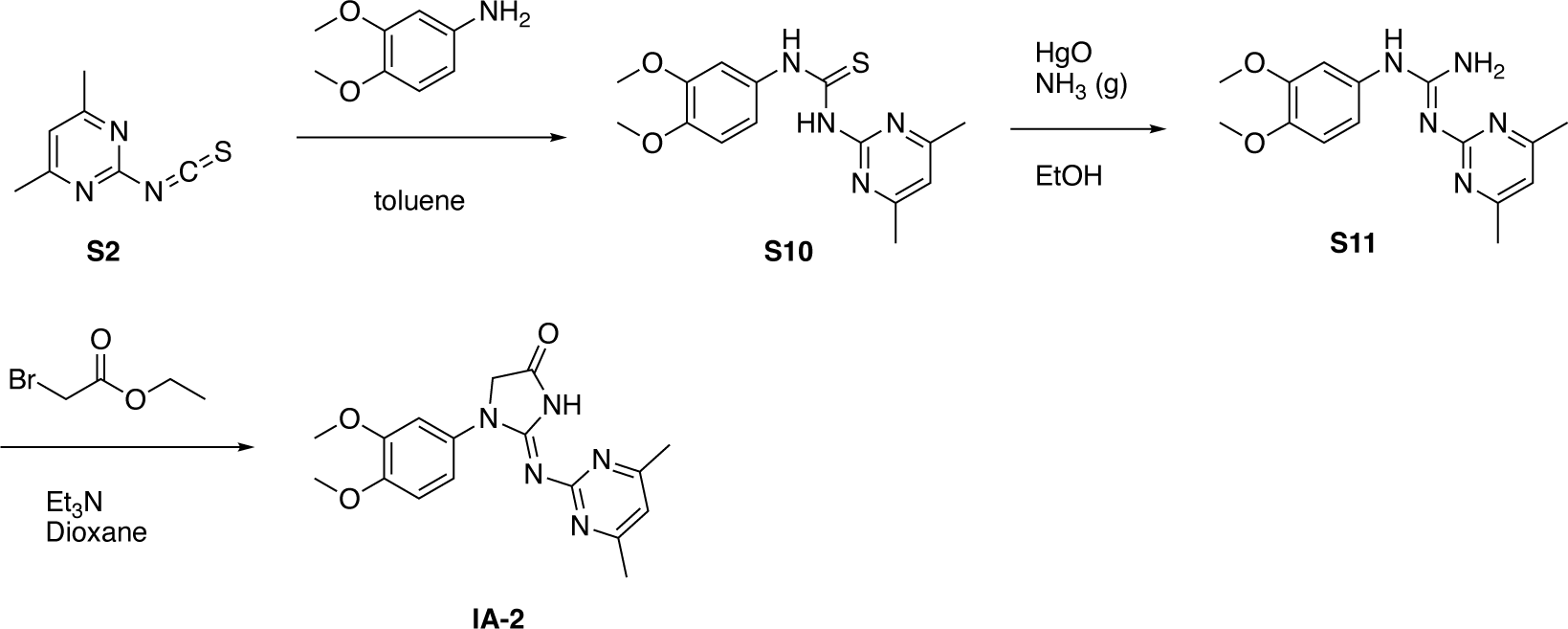

### 1-(3,4-Dimethoxyphenyl)-3-(4,6-dimethylpyrimidin-2-yl)thiourea (S10)

To a solution of 2-isothiocyanato-4,6-dimethyl-pyrimidine **S2** (2.00 g, 12.11 mmol) in toluene (30 mL) was added 3,4-dimethoxyaniline (2.04 g, 13.3 mmol). The reaction mixture was stirred at 25 °C for 1 hr. The reaction mixture was concentrated in vacuo to give a residue. The residue was triturated with methanol (20 mL), filtered and the filter cake was dried in vacuo to give the title compound **S10** (3.20 g, 83% yield) as a purple solid, which was used into the next step without further purification. ^1^H NMR (400MHz, DMSO-*d_6_*) δ = 13.50 (s, 1H), 10.60 (s, 1H), 7.49 (d, *J* = 2.4 Hz, 1H), 7.15 (dd, *J* = 2.4, 8.4 Hz, 1H), 6.98 (s, 1H), 6.96 (d, *J* = 8.4 Hz, 1H), 3.77 (s, 3H), 3.77 (s, 3H), 2.42 (s, 6H).

### 1-(3,4-Dimethoxyphenyl)-2-(4,6-dimethylpyrimidin-2-yl)guanidine (S11)

Ammonia gas was bubbled to EtOH (10 mL) to give a NH_3_-EtOH solution and 1-(3,4-dimethoxyphenyl)-3-(4,6-dimethylpyrimidin-2-yl)thiourea **S10** (800 mg, 2.51 mmol) was dissolved in above solution. Then HgO (2.72 g, 12.5 mmol) was added. The reaction mixture was stirred at 25 °C for 15 hrs. The reaction mixture was filtered and the filtrate was concentrated in vacuo to give a residue. The residue was purified by silica gel column chromatography (dichloromethane: methanol = 20: 1) to give the title compound **S11** (320 mg, 42% yield) as a light purple solid. LCMS (M+1)^+^: 302.3.

### 1-(3,4-Dimethoxyphenyl)-2-(4,6-dimethylpyrimidin-2-yl)imino-imidazolidin-4 -one (IA-2)

To a solution of 1-(3,4-dimethoxyphenyl)-2-(4,6-dimethylpyrimidin-2-yl)guanidine **S11** (160 mg, 530 umol) in dioxane (15 mL) was added ethyl 2-bromoacetate (266 mg, 1.59 mmol) and triethylamine (214 mg, 2.12 mmol). The reaction mixture was stirred at 90 °C for 12 hrs. On completion, the reaction mixture was concentrated in vacuo. The residue was diluted with ice water (25 mL) and extracted with dichloromethane (3 X 15 mL). The organic layer was dried over anhydrous Na_2_SO_4_, filtered, and concentrated in vacuo to get a residue. The residue was purified by Prep-HPLC (column: Phenomenex Gemini 150×25mm, 10um; mobile phase: [water (10mM NH_4_HCO_3_)-acetonitrile]) to give the title compound **IA-2** (15.5 mg, 8.4% yield) as a white solid. LCMS (M+1)^+^: 342.3. ^1^H NMR (400MHz, DMSO-*d_6_*) δ = 11.42 (br s, 1H), 7.45 (d, *J* = 2.0 Hz, 1H), 7.28 (dd, *J* = 2.0, 8.8 Hz, 1H), 6.98 (d, *J* = 8.8 Hz, 1H), 6.78 (s, 1H), 4.53 (s, 2H), 3.76 (s, 3H), 3.75 (s, 3H), 2.33 (s, 6H).

## Synthesis of analog 1401-A

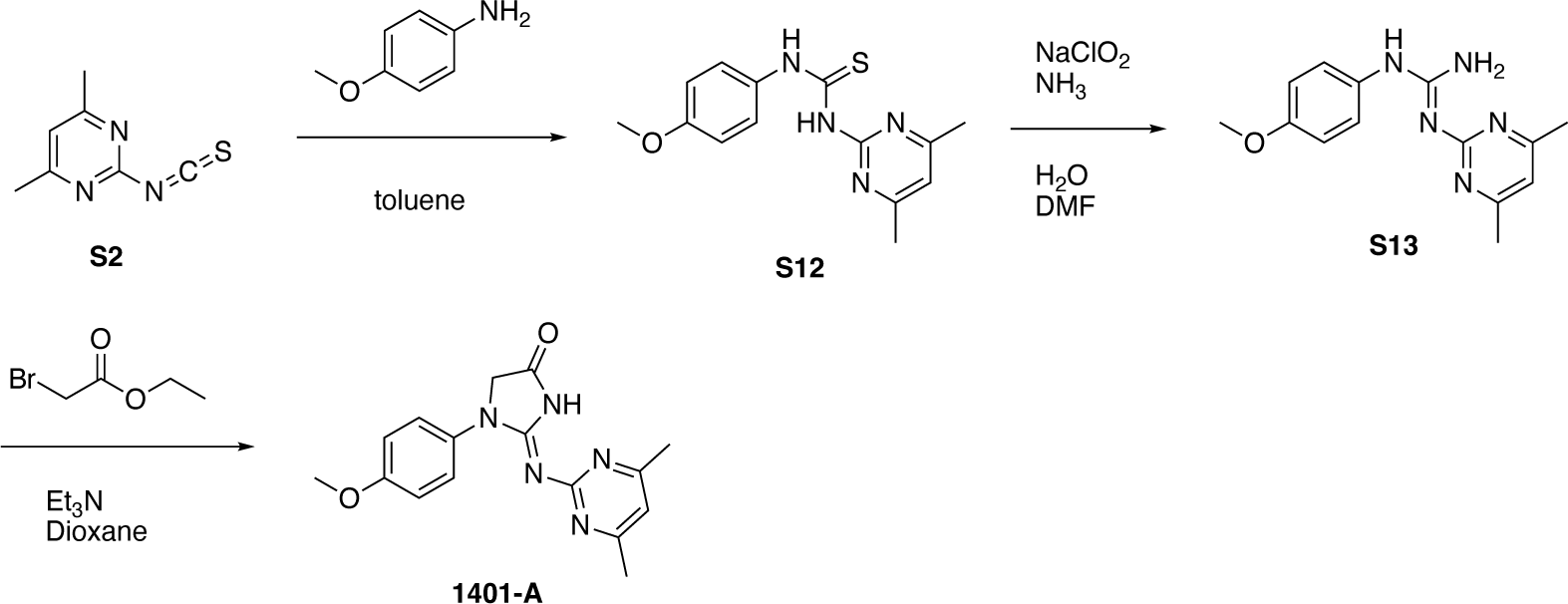

### 1-(4,6-Dimethylpyrimidin-2-yl)-3-(4-methoxyphenyl)thiourea (S12)

A mixture of 2-isothiocyanato-4,6-dimethyl-pyrimidine (2.00 g, 12.1 mmol) and 4-methoxyaniline (1.64 g, 13.3 mmol) in toluene (5 mL) was stirred at 90 °C for 12 hrs. The reaction mixture was filtered and the filter cake was washed with ethyl acetate (20 mL) to give the title compound **S12** (3.00 g, 86% yield) as a gray solid. ^1^H NMR (400 MHz, CDCl_3_) δ = 10.60 (s, 1H), 7.59 (d, *J* = 8.8 Hz, 2H), 7.03 - 6.91 (m, 3H), 3.77 (s, 3H), 2.42 (s, 6H).

### (*E*)-2-(4,6-dimethylpyrimidin-2-yl)-1-(4-methoxyphenyl)guanidine (S13)

A mixture of 1-(4,6-dimethylpyrimidin-2-yl)-3-(4-methoxyphenyl)thiourea **S12** (600 mg, 2.08 mmol), aqueous NH_3_ (30% purity, 6 mL) and NaClO_2_ (376 mg, 4.16 mmol) in DMF (10 mL) was stirred at 80 °C for 3 hrs. The reaction mixture was poured into ice-water (40 mL) and the mixture was filtered. The filter cake was dried in vacuo to give the title compound **S13** (200 mg, 35% yield) as yellow solid. LCMS (M+1)^+^: 272.1.

### (*E*)-2-((4,6-dimethylpyrimidin-2-yl)imino)-1-(4-methoxyphenyl)imidazolidin-4-one (1401-A)

A mixture of 2-(4,6-dimethylpyrimidin-2-yl)-1-(4-methoxyphenyl)guanidine **S13** (100 mg, 368 umol), ethyl 2-bromoacetate (184 mg, 1.11 mmol) and triethylamine (186 mg, 1.84 mmol) in dioxane (5 mL) was stirred at 90 °C for 5 hrs. The reaction mixture was concentrated in vacuo. The residue was purified by Prep-HPLC (column: Phenomenex Gemini 150×25mm, 10um; mobile phase: [water(0.04%NH_3_+10mM NH_4_HCO_3_)-ACN]; B%: 20%-50%,10min) to give the title compound **1401-A** (41.3 mg, 35% yield) as a white solid. LCMS (M+1)^+^: 312.2. ^1^H NMR (400 MHz, DMSO-*d_6_*) δ = 11.38 (br s, 1H), 7.71 (d, *J* = 9.2 Hz, 2H), 6.98 (d, *J* = 9.2 Hz, 2H), 6.78 (s, 1H), 4.50 (s, 2H), 3.77 (s, 3H), 2.34 (s, 6H).

## Synthesis of analog 1401-B

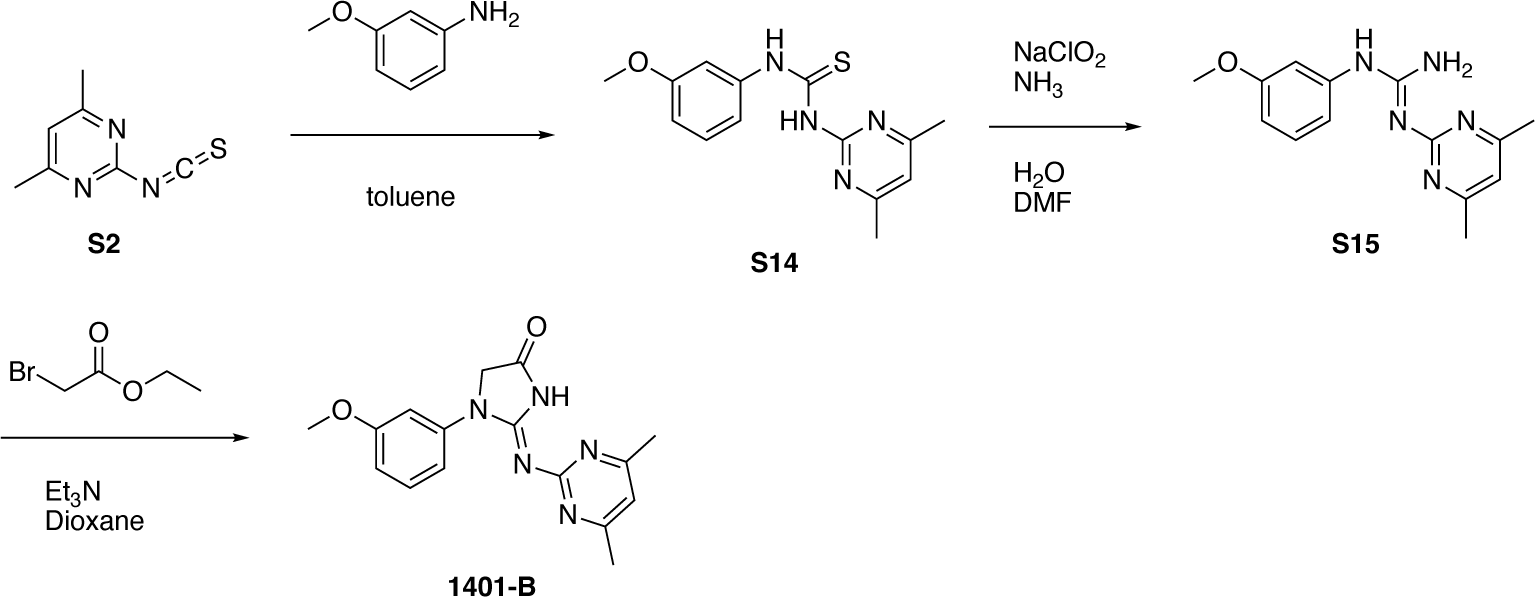

### 1-(4,6-dimethylpyrimidin-2-yl)-3-(3-methoxyphenyl)thiourea (S14)

To a solution of 2-isothiocyanato-4,6-dimethyl-pyrimidine **S2** (2.00 g, 12.1 mmol) in toluene (30 mL) was added 3-methoxyaniline (1.64 g, 13.3 mmol). The reaction mixture was stirred at 25 °C for 1 hr. The reaction mixture was concentrated in vacuo. The residue was diluted with methanol (20 mL) and filtered. The filter cake was dried in vacuo to give the title compound **S14** (2.70 g, 77% yield) as a white solid, which was used into the next step without further purification. ^1^H NMR (400MHz, DMSO-*d_6_*) δ = 13.72 (s, 1H), 10.69 (s, 1H), 7.60 (t, *J* = 2.0 Hz, 1H), 7.35 - 7.28 (m, 1H), 7.25 - 7.18 (m, 1H), 6.98 (s, 1H), 6.84 - 6.77 (m, 1H), 3.77 (s, 3H), 2.42 (s, 6H).

### 2-(4,6-Dimethylpyrimidin-2-yl)-1-(3-methoxyphenyl)guanidine (S15)

To a solution of 1-(4,6-dimethylpyrimidin-2-yl)-3-(3-methoxyphenyl)thiourea **S14** (500 mg, 1.73 mmol) in DMF (5 mL) and H_2_O (10 mL) was added aqueous NH_3_ (1.13 g, 8.67 mmol, 1.24 mL, 27% purity) and sodium chlorite (784 mg, 8.67 mmol). The reaction mixture was stirred at 80 °C for 15 hrs. The reaction mixture was concentrated in vacuo. The residue was diluted with water (50 mL) and extracted with DCM (3 X 45 mL). The organic layer was dried over anhydrous Na_2_SO_4_, filtered and concentrated in vacuo to give a residue. The residue was purified by silica gel column chromatography (dichloromethane: methanol = 20: 1) to give the title compound **S15** (170 mg, 36% yield) as a light brown solid. ^1^H NMR (400MHz, CDCl_3_) δ = 7.23 −7.18 (m, 1H), 6.70 - 6.59 (m, 3H), 6.54 (s, 1H), 3.73 (s, 3H), 2.31 (s, 6H).

### 3 - 2-(4,6-Dimethylpyrimidin-2-yl)imino-1-(3-methoxyphenyl)imidazolidin-4-one (1401-B)

To a solution of 2-(4,6-dimethylpyrimidin-2-yl)-1-(3-methoxyphenyl)guanidine **S15** (130 mg, 479 umol) in dioxane (15 mL) was added ethyl 2-bromoacetate (240 mg, 1.44 mmol) and triethylamine (193 mg, 1.92 mmol). The reaction mixture was stirred at 90 °C for 12 hrs. The reaction mixture was concentrated in vacuo to get a residue. The residue was diluted with ice-water (25 mL) and extracted with DCM (3 X 15 mL). The organic layer was dried over anhydrous Na_2_SO_4_, filtered and concentrated in vacuo to give a residue. The residue was purified by Prep-HPLC (column: Phenomenex Gemini 150×25mm, 10um; mobile phase: [water(10mM NH_4_HCO_3_)-ACN]) to give the title compound **1401-B** (56.1 mg, 36% yield) as a white solid. LCMS (M+1)^+^: 312.1. ^1^H NMR (400MHz, DMSO-*d_6_*) δ = 10.19 (br s, 1H), 7.55 (t, *J* = 2.4 Hz, 1H), 7.42 (dd, *J* = 2.0, 8.0 Hz, 1H), 7.34 - 7.24 (m, 1H), 6.81 (s, 1H), 6.74 (dd, *J* = 2.4, 8.0 Hz, 1H), 4.52 (s, 2H), 3.78 (s, 3H), 2.35 (s, 6H).

## Synthesis of analog 1401-C

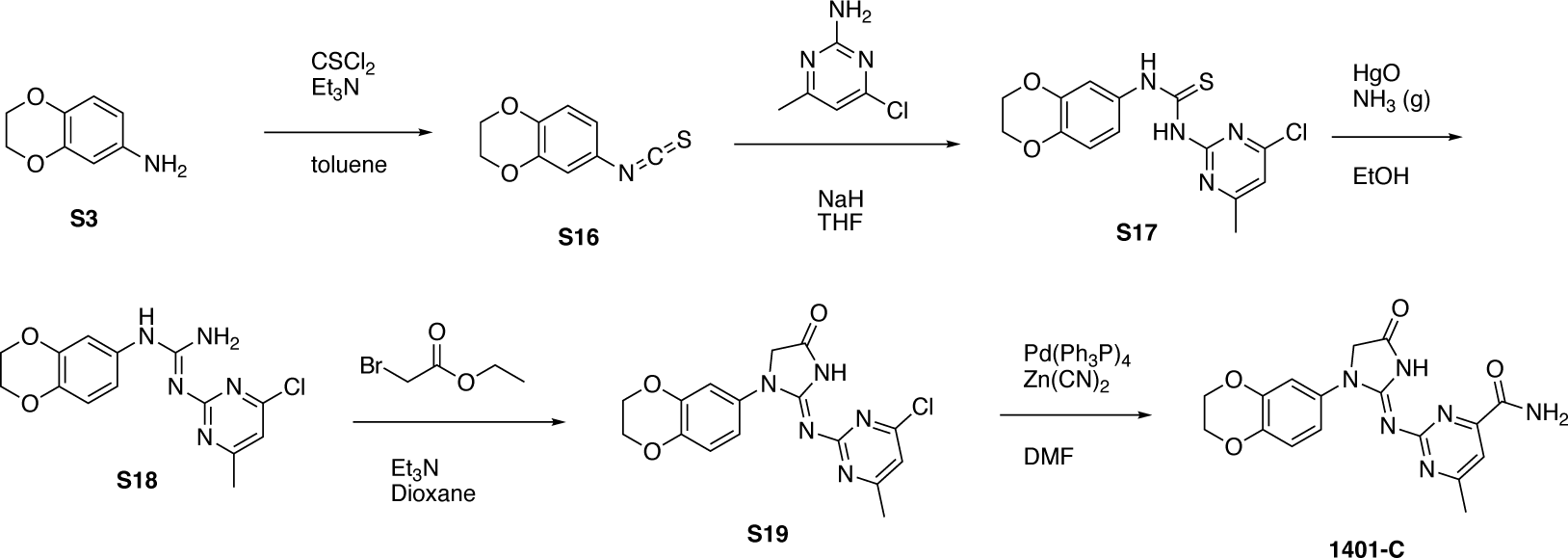

### 1 - 6-Isothiocyanato-2,3-dihydrobenzo[b][1,4]dioxine (S16)

To a solution of 2,3-dihydro-1,4-benzodioxin-6-amine **S3** (14.0 g, 92.6 mmol) and TEA (23.4 g, 231 mmol) in toluene (100 mL) was added a solution of thiocarbonyl dichloride (10.6 g, 92.6 mmol) in toluene (50 mL) at 0 °C, then the reaction mixture was stirred at 80 °C for 30 minutes. The reaction mixture was filtered and the filtrate was concentrated in vacuo. The residue was purified by silica gel column chromatography (PE: EA = 100: 1) to give the title compound **S16** (6.00 g, 34% yield) as a white solid. ^1^H NMR (400 MHz, CDCl_3_) δ = 6.83 - 6.79 (m, 1H), 6.78 - 6.70 (m, 2H), 4.28 - 4.25 (m, 4H).

### 1-(4-Chloro-6-methylpyrimidin-2-yl)-3-(2,3-dihydrobenzo[b][1,4]dioxin-6-yl)thiourea (S17)

To a mixture of NaH (2.09 g, 52.2 mmol, 60% purity) in THF (130 mL) was added 4-chloro-6-methyl-pyrimidin-2-amine (5.00 g, 34.8 mmol) at 0 °C. The reaction mixture was stirred at 0 °C for 0.5 hour, then 6-isothiocyanato-2,3-dihydro-1,4-benzodioxine **S16** (8.07 g, 41.7 mmol) was added. The reaction mixture was stirred at 50 °C for 1 hour. The reaction mixture was quenched with water (50 mL) at 0 °C and extracted with ethyl acetate (3 × 20 mL). The combined organic layer was dried over Na_2_SO_4_, filtered and concentrated in vacuo to give a white solid. The solid was triturated with EtOH (5 mL) to give the title compound **S17** (8.00 g, 68% yield) as a white solid. ^1^H NMR (400 MHz, DMSO-*d*_6_) δ = 12.94 (br s, 1H), 11.14 (s, 1H), 7.37 (d, *J* = 2.4 Hz, 1H), 7.28 (s, 1H), 7.03 (dd, *J* = 2.4, 8.8 Hz, 1H), 6.93 - 6.84 (m, 1H), 4.26 - 4.22 (m, 4H), 2.48 (s, 3H).

### 1-(4-Chloro-6-methylpyrimidin-2-yl)-3-(2,3-dihydrobenzo[b][1,4]dioxin-6-yl)guanidine (S18)

A mixture of 1-(4-chloro-6-methyl-pyrimidin-2-yl)-3-(2,3-dihydro-1,4-benzodioxin-6- yl)thiourea **S17** (8.00 g, 23.7 mmol), HgO (21.3 g, 98.5 mmol) and NH_3_/EtOH (80 mL) in a sealed tube was stirred at 15 °C for 12 hours. The reaction was filtered and the filter cake was washed with DCM (3 × 500 ml). The combined organic layer was concentrated in vacuo to give the title compound **S18** (5.00 g, 60% yield) as a yellow solid. LCMS (M+1)^+^: 320.1.

### (*E*)-2-((4-chloro-6-methylpyrimidin-2-yl)imino)-1-(2,3-dihydrobenzo[b][1,4]dioxin-6-yl) imidazolidin-4-one (S19)

A mixture of 2-(4-chloro-6-methyl-pyrimidin-2-yl)-1-(2,3-dihydro-1,4-benzodioxin-6-yl)guanidine **S18** (3.60 g, 11.2 mmol) and ethyl 2-bromoacetate (1.88 g, 11.2 mmol) in dioxane (30 mL) was stirred at 110 °C for 0.5 hour under microwave. The reaction was concentrated in vacuo and the residue was purified by reverse phase chromatography to give the title compound **S19** (1.00 g, 19% yield) as a yellow solid. LCMS (M+1)^+^: 360.1.

### (*E*)-2-((1-(2,3-dihydrobenzo[b][1,4]dioxin-6-yl)-4-oxoimidazolidin-2-ylidene)amino)-6-methylpyrimidine-4-carboxamide (1401-C)

A mixture of (2*E*)-2-(4-chloro-6-methyl-pyrimidin-2-yl)imino-1-(2,3-dihydro-1,4-benzodioxin-6-yl) imidazolidin-4-one **S19** (200 mg, 555 umol), Pd(PPh_3_)_4_ (64.2 mg, 55.5 umol) and Zn(CN)_2_ (130 mg, 1.11 mmol) in DMF (15 mL) was stirred at 130 °C for 3 hours. The reaction was filtered, and the filtrate was concentrated in vacuo. The residue was purified by Prep-HPLC (column: Phenomenex luna C18 150×25mm, 10um; mobile phase: [water (0.225% formic acid)-ACN]; B%: 14%-34%, 7.8min) to give the title compound **1401-C** (9.00 mg, 5% yield) as an off-white solid. LCMS (M+1)^+^: 369.2. ^1^H NMR (400 MHz, DMSO-*d*_6_) δ = 8.23 (br s, 1H), 7.42 (d, *J* = 2.4 Hz, 1H), 7.14 (dd, *J* = 2.4, 8.8 Hz, 1H), 6.86 (d, *J* = 8.8 Hz, 1H), 6.22 (s, 1H), 4.38 (s, 2H), 4.26 - 4.21 (m, 4H), 2.25 (s, 3H).

## QUANTIFICATION AND STATISTICAL ANALYSIS

Quantification of trypsin digestion was done by measuring mean band intensity of the LPS-protected band of OprH (∼14.5 kDa; Figure S10F) using Fiji (Image J) after Coomassie staining of SDS-PAGE gels (Method Details). After background subtraction the intensity was normalized to vehicle treated sample and plotted using GraphPad Prism version 10. Mean band intensity of protein bands of interest (LPS or OprL-FLAG) detected by western blotting was measured using Fiji (ImageJ software) with background subtraction. This value was then normalized to the band intensity of the loading control, which was StrepII-OprH in the pull-down experiments. This ratio was then normalized to the vehicle or control treatment sample and plotted using GraphPad Prism version 10. Statistical significance for the epithelial cell infection, LPS pull-down with StrepII-OprH, membrane fluidity and the intra-bacterial accumulation assays were calculated with the unpaired Student’s parametric *t*-test with Welch’s correction in GraphPad Prism version 10. Statistical significance for the trypsin digestion and the OprL pull-down with StrepII-OprH assays were calculated with the One sample *t*-test in comparison to the control (mean = 1) in GraphPad Prism version 10.

