## Supplemental Figures and Supplemental Table S2 for "“Multiplexed screen identifies a *Pseudomonas aeruginosa*-specific small molecule targeting the outer membrane protein OprH and its interaction with LPS”"

**SUPPLEMENTAL INFORMATION**

Supplemental figures S1 to S12

Supplemental table S1 (excel): Promoters, primers, plasmids and strains used in this study.

Supplemental table S2: Multiplex screen analysis

Supplemental table S3 (excel): Gene expression data (read counts) from the two RNA-Seq datasets reported in this study.

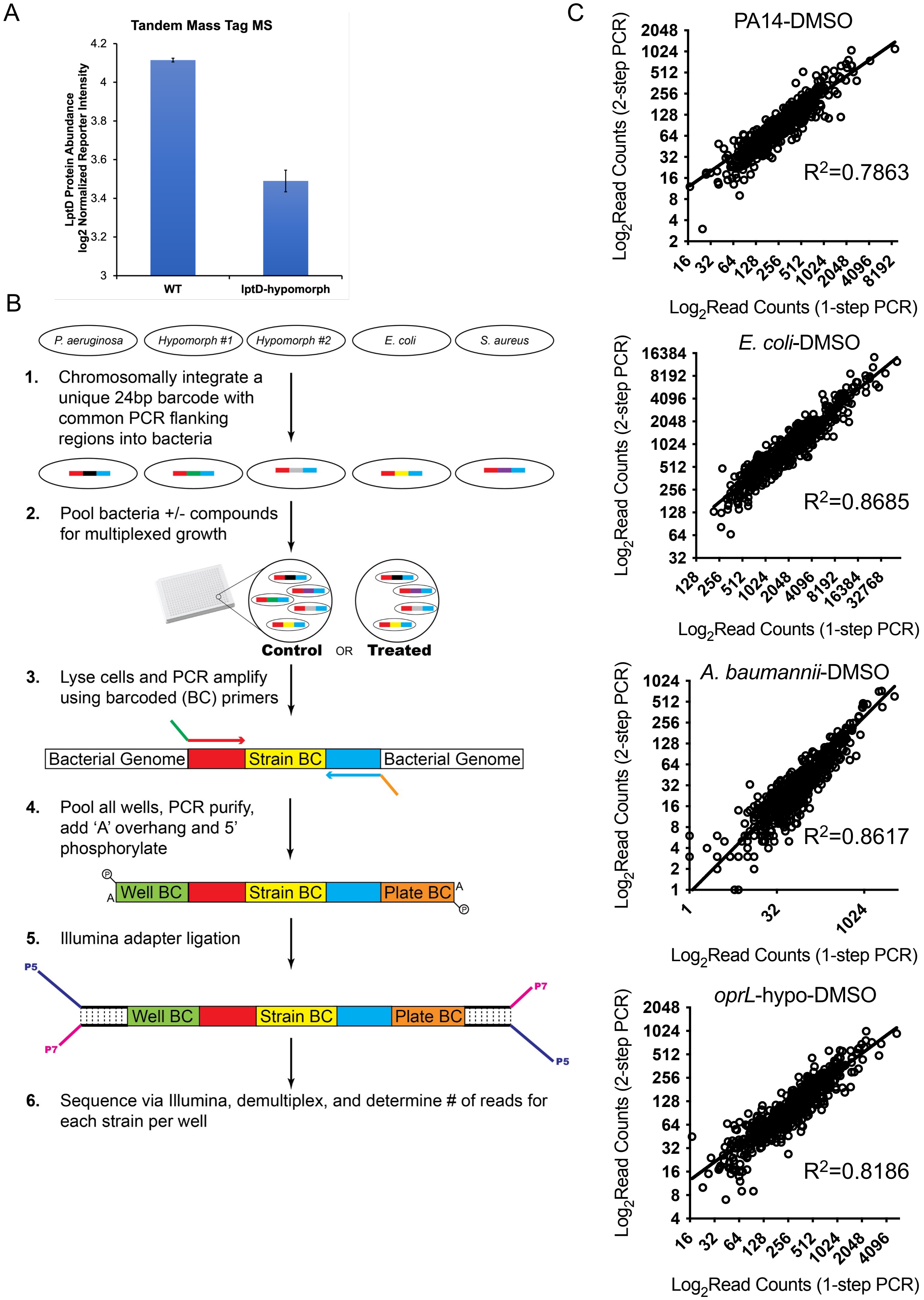
A
Tandem Mass Tag MS
LptD Protein Abundance
log2 Normalized Reporter Intensity
4.2
4
3.8
3.6
3.4
3.2
3
WT
lptD-hypomorph
B
P. aeruginosa
Hypomorph #1
Hypomorph #2
E. coli
S. aureus
1. Chromosomally integrate a unique 24bp barcode with common PCR flanking regions into bacteria
2. Pool bacteria +/- compounds for multiplexed growth
Control
OR
Treated
3. Lyse cells and PCR amplify using barcoded (BC) primers
Bacterial Genome
Strain BC
Bacterial Genome
4. Pool all wells, PCR purify, add 'A' overhang and 5' phosphorylate
Well BC
Strain BC
Plate BC
5. Illumina adapter ligation
P5
P7
Well BC
Strain BC
Plate BC
P7
P5
6. Sequence via Illumina, demultiplex, and determine # of reads for each strain per well
C
PA14-DMSO
$R^2$=0.7863
$Log_2$Read Counts (2-step PCR)
$Log_2$Read Counts (1-step PCR)
E. coli-DMSO
$R^2$=0.8685
$Log_2$Read Counts (2-step PCR)
$Log_2$Read Counts (1-step PCR)
A. baumannii-DMSO
$R^2$=0.8617
$Log_2$Read Counts (2-step PCR)
$Log_2$Read Counts (1-step PCR)
oprL-hypo-DMSO
$R^2$=0.8186
$Log_2$Read Counts (2-step PCR)
$Log_2$Read Counts (1-step PCR)

**Figure S1.** A) Quantitative proteomics with tandem mass tags (TMT) mass spectrometry is used to measure the amount of LptD protein in wildtype PA14 and the *lptD*-hypomorph strains. LptD was depleted by ~50% in the *lptD*-hypomorph strain compared to wildtype strain (n = 3, error bars indicate SEM). B) Schematic of the molecular biology protocol employed for adaptor ligation-based library construction strategy. C) Comparison of the two library construction methods. $Log_2$-transformed read counts generated from vehicle (DMSO)-treated wells by Illumina adaptor ligation-based library construction method (2-step PCR) is plotted against the read counts generated by the traditional, single-step PCR library construction method with long primers (1-step PCR). The multiplexed pool was grown for 12 hours before samples were processed. Data from wildtype strains of 3 gram-negative species, *P. aeruginosa*, *E. coli*, and *A. baumannii*, and 1 *P. aeruginosa* hypomorph strain, *oprL*-hypomorph, is shown. High correlation is observed between the read counts obtained from libraries generated by the two methods.

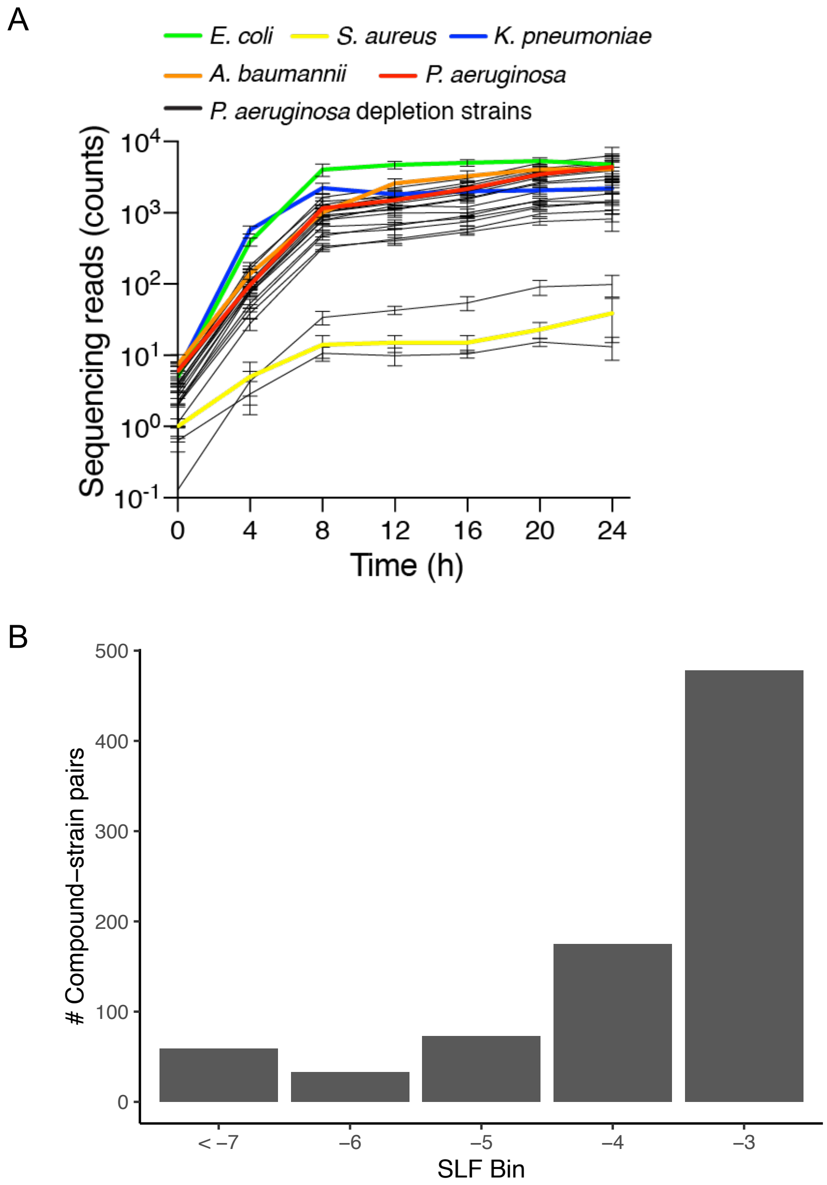


**Figure S2. Development of the multiplexed assay and analysis of compound activity detected by the multiplexed screen.** A) Growth over time is determined using mean sequencing read counts of hypomorph (depletion) strains (black lines) and wildtype species (colored lines) in a multiplexed pool. Starting total cell density of an evenly mixed pool was $OD_{600nm} = 5 \times 10^{-4}$, which is approximately $5 \times 10^5$ CFU/mL. Measurements were made at 4-hour intervals between 0 and 24 h (n = 12; error bars indicate SD). B) Shown is the distribution of SLF values for compound-strain pairs with an SLF value of less than –3, representing 0.09% of the total number of compound-strain pairs for *P. aeruginosa* strains in the screen. The number of compound-strain interactions in the multiplexed screen with a *P. aeruginosa* strain is plotted against the sensitivity indicated by the SLF bin, where the SLF is between the listed bin value and the bin value to its left (e.g., bin "–5" indicates $-6 \leq SLF < -5$).

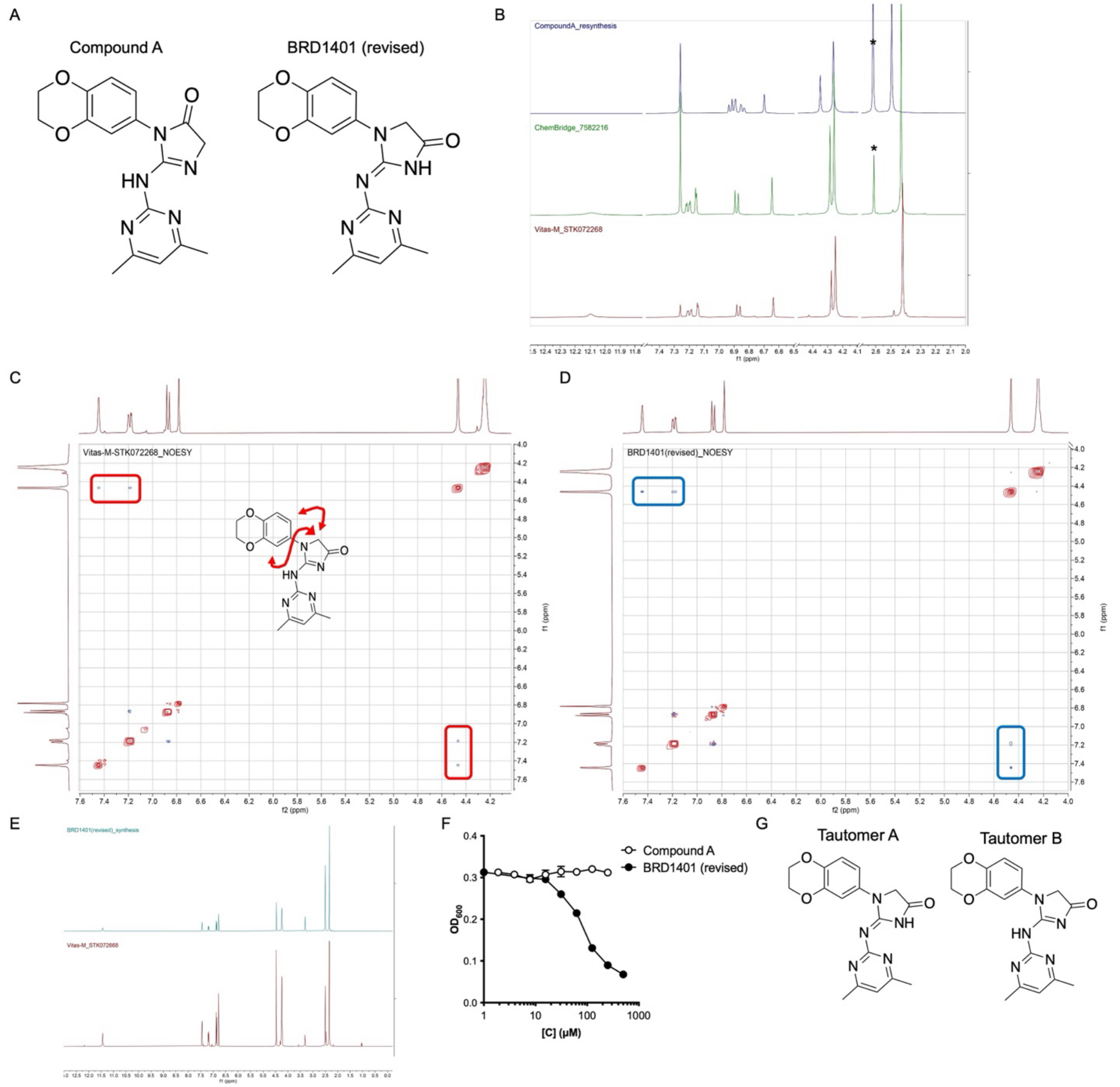


**Figure S3. Elucidation of BRD1401 structure.** A) Structure of BRD1401 hit compound based on library annotation, "compound A", and after structure elucidation, "BRD1401 (revised)", with NMR studies. B) $^{1}H$-NMR of synthesized "compound A" and the two molecules available from ChemBridge and Vitas-M in $CDCl_3$. The NMR spectra of the compounds from ChemBridge and Vitas-M are superimposable, while that of compound A is distinct. * indicates a peak derived from DMSO in which stocks of "compound A" and ChemBridge compound were made. C) 2D NOESY of STK072268 from Vitas-M. Correlation peaks between imidazolone methylene peak and two phenyl hydrogens are highlighted in the red boxes and the corresponding positions in the structure highlighted with red arrows. D) 2D NOESY of synthesized "BRD1401 (revised)". Correlation peaks identical to the ones observed in (C) are highlighted in the blue boxes. E) $^{1}H$-NMR of synthesized "BRD1401 (revised)"and the Vitas-M molecule (STK072268) in DMSO-$d_6$. The two 2D NOESY and $^{1}H$-NMR spectra of "BRD1401 (revised)"and the Vitas-M molecule (STK072268) are superimposable and hence their chemical structures are identical. F) Activity of synthesized “compound A” and “BRD1401 (revised)” toward *oprL*-hypomorph measured by an absorbance ($OD_{600nm}$)-based growth over a period of 16 hours in presence of a dose range of BRD1401. $OD_{600nm}$ versus BRD1401 concentration is plotted (n=2; error bars indicate SD). G) Structures of the two tautomeric forms of BRD1401. Tautomer A structure has an exocyclic CN double bond while tautomer B has an endocyclic CN double bond. Tautomer A is reported in figure 4A.

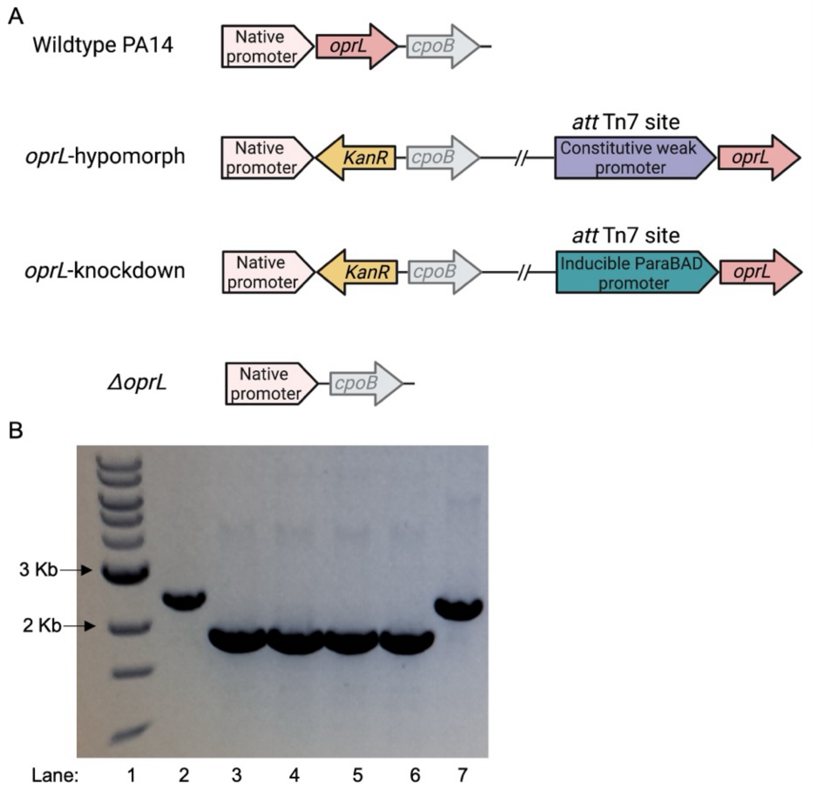


C

| Clone | PA14# | Gene/annotation | SNP | AA change | Functional implication |
|---|---|---|---|---|---|
| B | PA14_70270 | *algC* | T1481A | V494E | Unknown |
| | Intergenic between PA14_16280 & PA14_16290 | Upstream of *armR* | A(-59)G | NA | Unknown |
| | PA14_51710 | *oprL* | - | - | Deletion of gene |
| | PA14_23400 | *PA14_23400* | C947G | S316* | Premature stop, hence truncated protein |
| D | PA14_70270 | *algC* | T1481A | V494E | Unknown |
| | Intergenic between PA14_16280 & PA14_16290 | Upstream of *armR* | A(-59)G | NA | Unknown |
| | PA14_51710 | *oprL* | - | - | Deletion of gene |
| | PA14_67790 | *envC* | A274- | - | Mistranslation after K93, leading to premature stop |
| | PA14_23470 | *wbpM* | G1468T | V490F | Unknown |

**Figure S4.** A) Schematic of the *oprL* operon and the *att*Tn7 site in wildtype PA14, *oprL*-hypomorph (PA19 in Table S1G), *oprL*-KD (PA23 in Table S1G) and Δ*oprL*-MW (PA22 in Table S1G) strains used in this study. B) PCR confirmation of *oprL* knock-out in Δ*oprL*-MW strain. Lane 1) 1Kb Ladder 2) PA14 wildtype control strain from Marvin Whiteley's laboratory 3) Δ*oprL*-MW clone A 4) Δ*oprL*-MW clone B 5) Δ*oprL*-MW clone C 6) Δ*oprL*-MW clone D 7) PA14 wildtype strain (PA01 in Table S1G). Expected band size in wildtype strain with intact *oprL* gene is 2432 bp while in Δ*oprL* strain is 1925 bp. B) Table of mutations identified in the two clones, B & D, of the Δ*oprL*-MW strain. Common mutations in the two clones are highlighted in red.

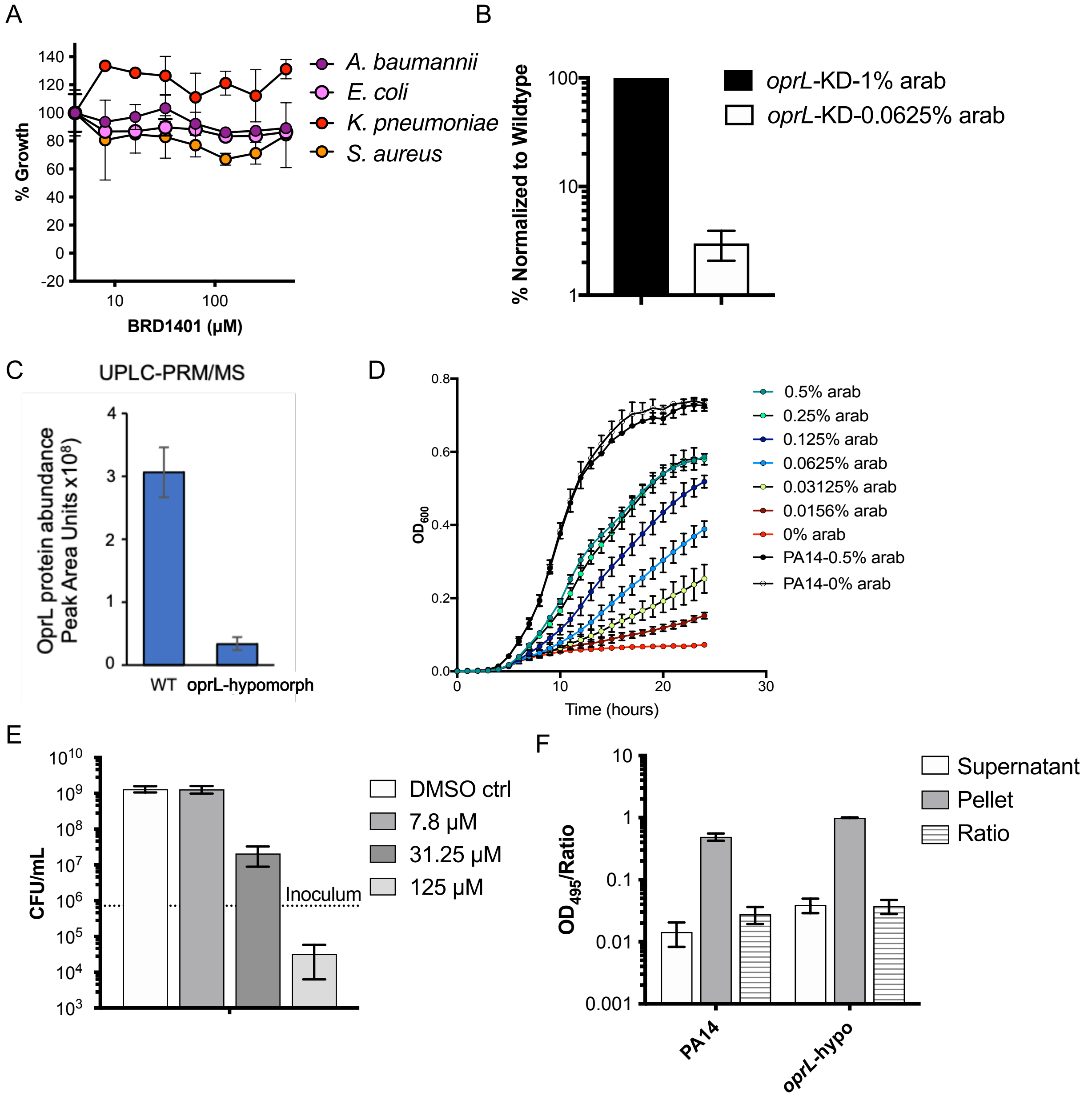


G

| Antibiotic | MIC (μM ) | |
|---|---|---|
| | Wildtype (PA14) | *oprL*-hypomorph |
| Ciprofloxacin | 0.25 | 0.125 |
| Amikacin | 1.25 | 2.5 |
| Doxycycline | 50 | 25 |
| Trimethoprim | 250 | 125 |

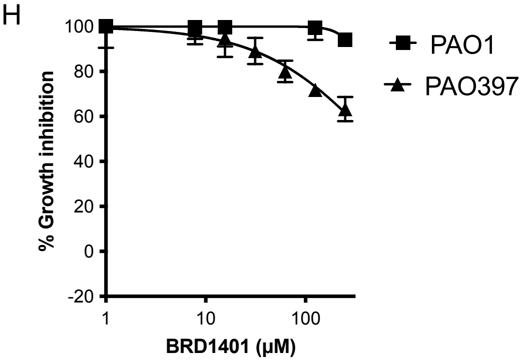


**Figure S5. Characterization of BRD1401 activity and of the *oprL*-KD and *oprL*-hypo strains.** A) BRD1401 activity towards bacterial species other than *P.aeruginosa* that were included in the multiplexed screen after exposure to a dose range of BRD1401 over 16 hours. Normalized growth, calculated in % relative to positive and negative controls, which was measured by an absorbance ($OD_{600nm}$)-based assay, is plotted against BRD1401concentration (n=2; error bars indicate SD). B) Label-free proteomics (see Method Details) was used to quantify OprL peptide levels in the *oprL*-KD or wildtype PA14 strains grown to mid-log phase in media with high (1%) or low (0.0625%) levels of arabinose (arab). Total peptide counts were normalized to amounts detected in the wildtype strain exposed to identical concentrations of arabinose (n = 3, error bars indicate SD). C) Label-free quantitative proteomics is used to measure the amount of OprL protein in wildtype PA14 and the *oprL*-hypomorph strains. OprL was depleted by 89% in the *oprL*-hypomorph strain compared to wildtype strain (n = 3, error bars indicate SEM). D) Growth kinetic of the *oprL*-KD strain or wildtype PA14 strain measured by absorbance at 600nm ($OD_{600}$) over a period of 24 hours in media with varying concentrations of arabinose (arab) (n = 3, error bars indicate SD). OprL expression is under the control of the arabinose inducible $P_{araBAD}$ promoter in the *oprL*-KD strain. F) Bactericidal activity of BRD1401 towards the *oprL*-KD strain was measured by enumerating colony forming units (CFU/mL) after exposure to varying concentration of BRD1401 or vehicle control (DMSO) for 17h in media with low (0.0625%) or high (0.5%) arabinose. 0.0625% arabinose led to down regulation of *oprL* expression (n = 3, error bars indicate SD). G) AmpC activity, as detected by absorbance at 495nm ($OD_{495}$), in the supernatant or bacterial pellet of wildtype PA14 or the *oprL*-hypomorph strains grown to log phase. The ratio of $OD_{495}$ in supernatant to the sum of $OD_{495}$ detected in the supernatant and bacterial pellet is also plotted (n = 3, error bars indicate SD). This ratio is a measure of the normalized amount of AmpC detected in the supernatant and directly correlates with outer membrane permeabilization. H) Table of minimum inhibitory concentrations (MIC) of antibiotics with known anti-pseudomonal activity against the *oprL*-hypomorph strain and wildtype PA14 strain as measured from the absorbance-based growth assay. I) BRD1401 activity towards the efflux deficient PAO397 strain in PAO1 background and the parent PAO1 strain measured by the absorbance-based assay. Normalized growth calculated in % is plotted against BRD1401 concentration (n = 3, error bars indicate SD).

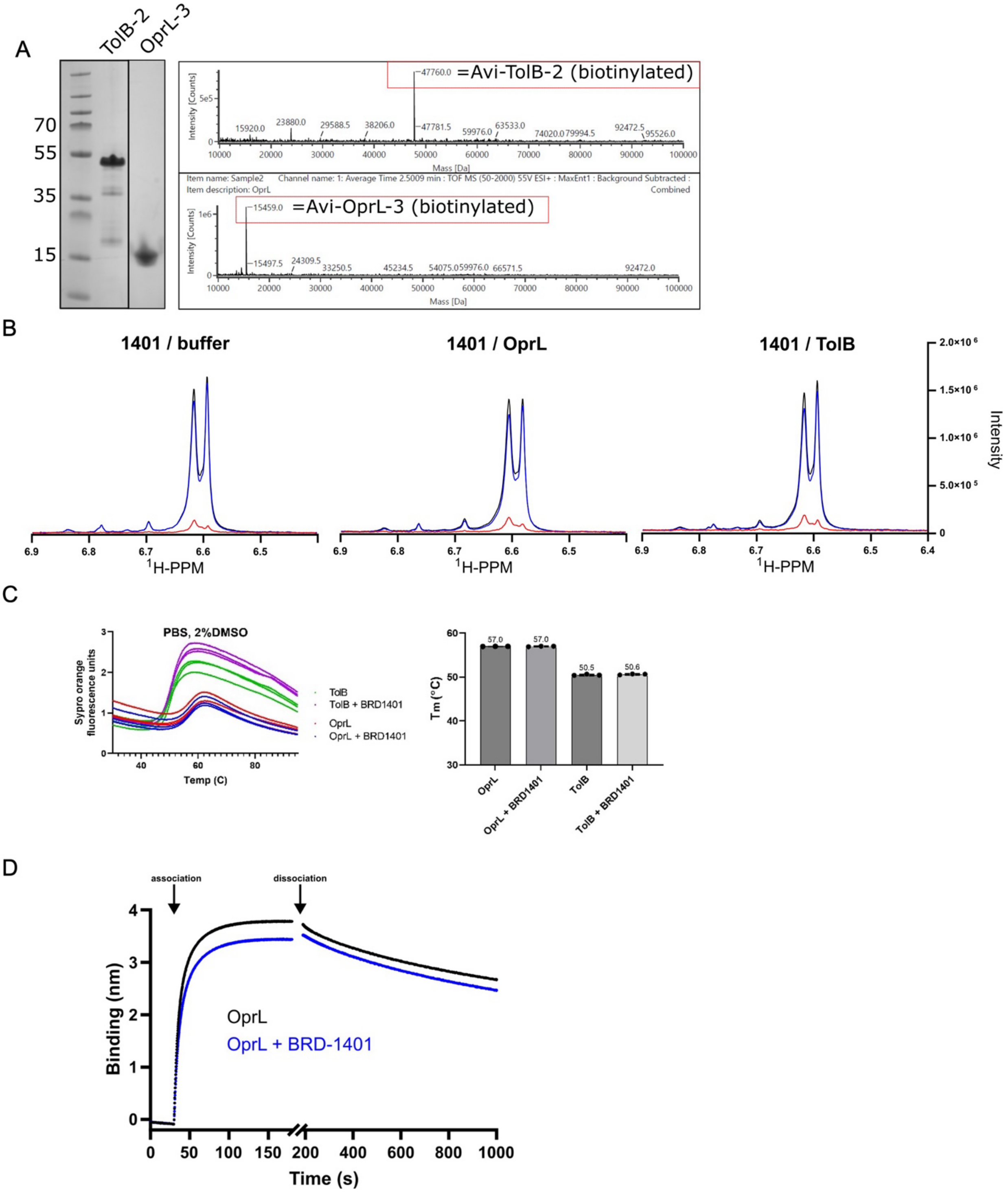


**Figure S6. Biophysical studies with BRD1401 and TolB-OprL proteins.** A) Mass spectrometry and gel-based qualitative analysis of purified TolB and OprL proteins. A clear band at ~47 kDa corresponding to Avi-tagged TolB and a band at ~15kDa corresponding to Avi-tagged OprL are visible in the gel. Peaks with high intensity corresponding to the protein masses are also detected by intact mass-spectrometry of the purified samples. B) STD-NMR of BRD1401 with OprL and TolB proteins in PBS buffer. Overlaid 1H-spectra on the left show the aromatic region chemical shifts and intensities of 500 μM BRD1401 solution in PBS (see Method Details); these serve as a reference set. Spectra on the right correspond to 500 μM 1401 in the presence of 15 μM *P. aeruginosa* OprL or TolB: without irradiation of the protein (black), with irradiation of the protein (blue), and the difference spectrum (STD in red) between black and blue spectra corresponds to the magnitude of the Ligand-Protein distance-dependent Nuclear Overhauser Effect (NOE). There is no difference in the red spectra in the absence or presence of proteins indicating no binding to BRD1401. C) DSF evaluation of the melting temperature of TolB and OprL in the presence and absence of BRD1401 is plotted. BRD1401 has no impact on the melting temperature of the two proteins suggesting it does not bind to either protein. D) Biolayer Interferometry (BLI) binding analysis. TolB (at 200 nM) was immobilized on BLI sensors in binding buffer and OprL (10 μM) with and without 500 μM BRD-1401 were used as ligands to determine the association and dissociation binding kinetics. Fitting the dissociation phase yielded similar off rates in the presence 1.30 x $10^{-3}$ $s^{-1}$ (±3.22 x $10^{-6}$) or absence 1.37 x $10^{-3}$ $s^{-1}$ (±3.54 x $10^{-6}$) of BRD-1401 indicating BRD1401does not impact the TolB-OprL complex.

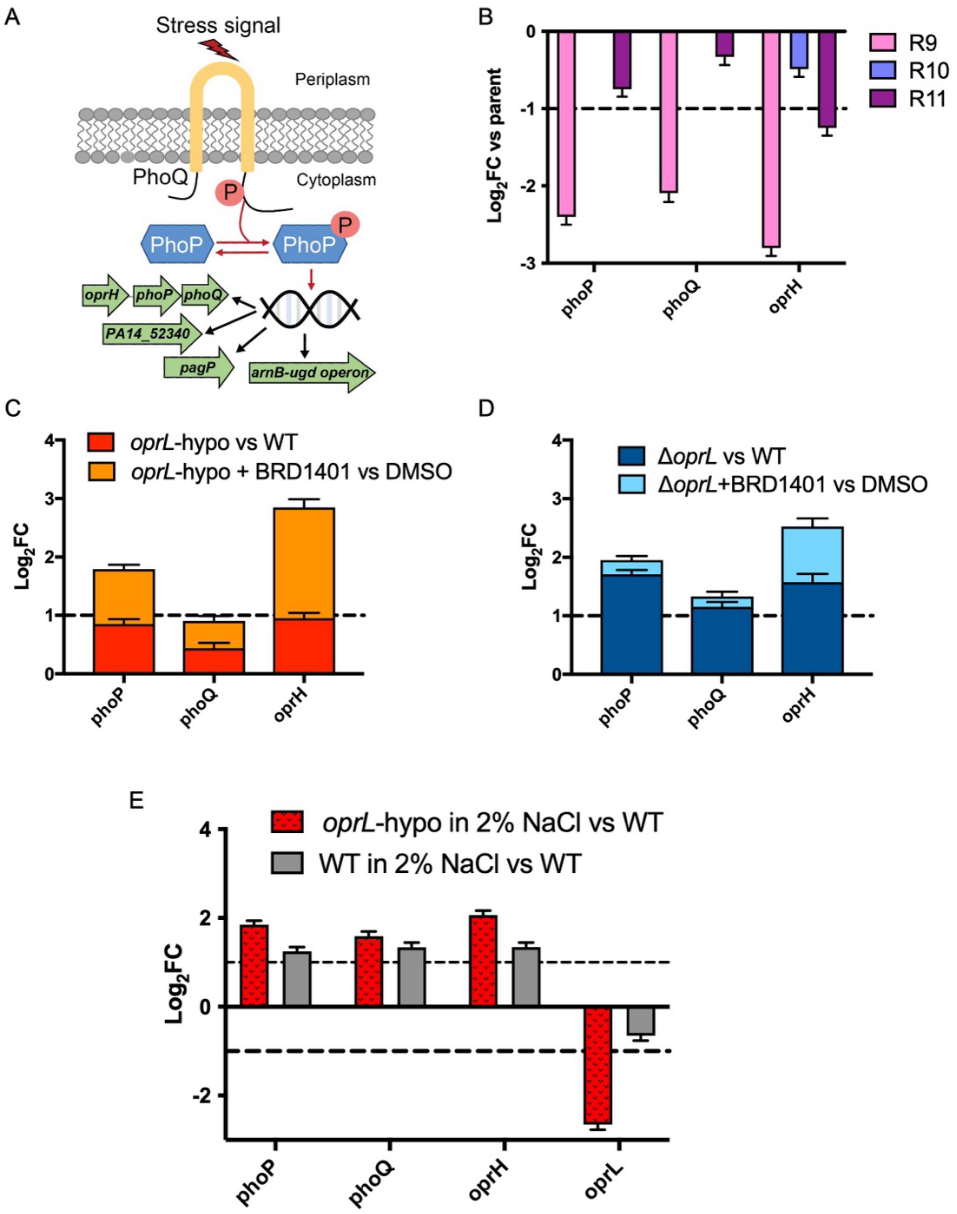


**Figure S7. Gene expression analysis of *oprH-phoPQ* operon and *oprL*.** A) Schematic of PhoPQ two component signaling transduction system in *P. aeruginosa*. Periplasmic domain of PhoQ senses stress signals that leads to autophosphorylation of the cytoplasmic domain and subsequent reversible phosphorylation and activation of the PhoP transcription factor. PhoP then upregulates ~20 genes including the *arnB-ugd* operon, *oprH-phoPQ* operon, *pagP* and *PA14_52340*.[53] B), C), D) and E) mRNA was isolated and processed for whole genome RNA sequencing to measure gene expression after genetic manipulation and/or compound exposure under various growth conditions. Relative gene expression was calculated and plotted as log2-transformed fold change ($Log_2FC$) relative to a parent strain or vehicle treatment using the DESeq pipeline. B) Fold change in expression of *phoP*, *phoQ* and *oprH* in the BRD1401-resistant clones, R9, R10 and R11, relative to the parent *oprL*-hypomorph strain (n = 3, error bars indicate SEM). B) Fold change in expression of *phoP*, *phoQ* and *oprH* in the *oprL*-hypomorph strain relative to wildtype PA14 strain or in the *oprL*-hypomorph strain exposed to 256 μM of BRD1401 for 120 min relative to DMSO vehicle control (n = 3, error bars indicate SEM). C) Fold change in expression of *phoP*, *phoQ* and *oprH* in the Δ*oprL* strain relative to wildtype PA14 strain or in the Δ*oprL* strain exposed to 256 μM of BRD1401 for 120 min relative to DMSO vehicle control (n = 3, error bars indicate SEM). D) Fold change in expression of *phoP*, *phoQ*, *oprH* and *oprL* in the *oprL*-hypomorph or wildtype PA14 strains grown in media with high salt (2% NaCl) relative to wildtype strain grown with normal (0.5% NaCl) salt for 90 minutes (n = 3, error bars indicate SEM).

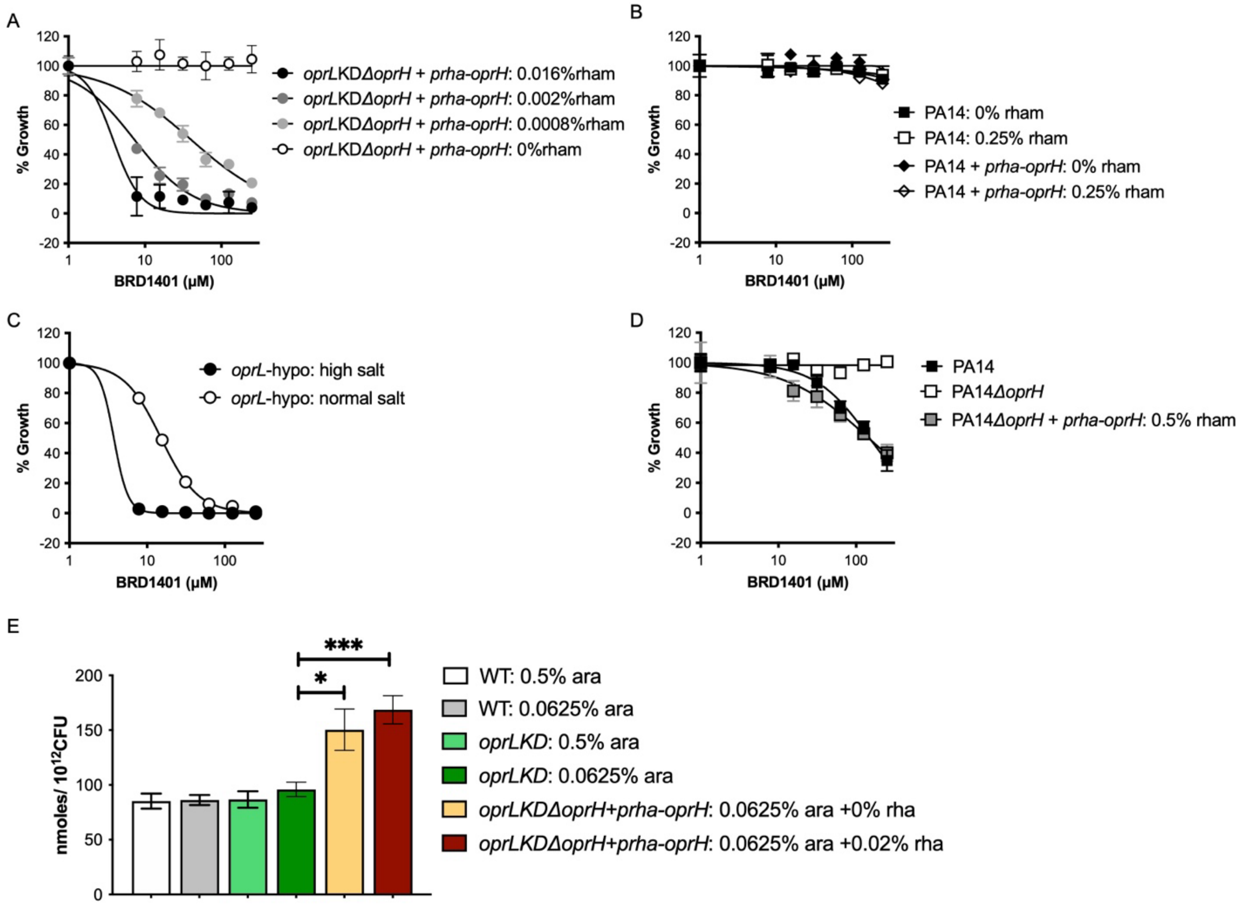


**Figure S8. BRD1401 activity and accumulation in wildtype, *oprL*-KD or *oprL*-hypomorph strains in various growth conditions.** A) Impact of OprH over-expression on BRD1401 activity. Normalized growth, calculated in % relative to vehicle (0.5% DMSO) and positive controls, measured from an absorbance ($OD_{600nm}$)-based growth assay of the *oprL*-KDΔ*oprH*/*pRha-oprH* strain (PA31 in Table S1G) is plotted against BRD1401 concentration. The strain was grown in media with 0.063% arabinose that downregulates OprL expression and varying concentrations of BRD1401 and rhamnose (rham) that drives episomal over-expression of *oprH* through the $P_{rhaBAD}$ promoter (n= 3, error bars indicate SD). Increased expression of *oprH* led to heightened sensitization to BRD1401. B) Activity of BRD1401 towards the wildtype PA14 and a PA14-derived strain over-expressing OprH (*[pRha-oprH]*; PA28 in Table S1G) measured as normalized growth with the absorbance-based assay is plotted against BRD1401 concentration (n= 3, error bars indicate SD). Episomal over-expression of *oprH* was induced by addition of 0.25% rhamnose (rham) or not induced with 0% rhamnose in standard LB medium. C) BRD1401 activity towards the *oprL*-hypomorph strain in media with either normal (0.5%; normal salt) or high (2%; high salt) levels of NaCl. Normalized growth in %, measured with the absorbance-based assay, is plotted against BRD1401 concentration (n= 3, error bars indicate SD). D) BRD1401 activity towards wildtype PA14, PA14Δ*oprH* (PA26 in Table S1G) and *[pRha-oprH]* (PA28 in Table S1G) strains grown in LB with high salt (2% sodium chloride). Episomal over-expression of *oprH* was induced by addition of 0.5% rhamnose (rham) or not induced with 0% rhamnose in the *[pRha-oprH]* strain. Normalized growth inhibition in % measured with the absorbance-based assay, is plotted against BRD1401 concentration (n= 3, error bars indicate SD). E) Shown is intra-bacterial concentration of BRD1401 in nanomoles, normalized to bacterial number enumerated as CFU, in wildtype PA14

(WT), *oprL*-KD or *oprL*-KDΔ*oprH*/*pRha-oprH* (PA31 in Table S1G) strains exposed to 100 µM BRD1401 for 10 minutes in PBS at 37ºC. The strains were grown with variable rhamnose (rham) that controls OprH expression levels in the *oprL*-KDΔ*oprH*/*pRha-oprH* strain and arabinose (ara) that controls OprL expression in the *oprL*-KD-derived strains. 0.063% arabinose was used to downregulate *oprL* expression and thus sensitize to BRD1401, while 0.5% arabinose increased *oprL* expression to wildtype levels. Bacterial pellets were collected, lysed, and extracted into a methanol-water mixture, which was run on an LC-MS/MS to quantify compound accumulation (n= 6-12; error bars indicate SEM). BRD1401 intra-bacterial concentration increased with both deletion and over-expression of *oprH* showing there is no correlation between its activity and accumulation. *, *** indicate $p$-value < 0.05 and $p$-value < 0.001, respectively, calculated using an unpaired Student's parametric $t$-test with Welch's correction.

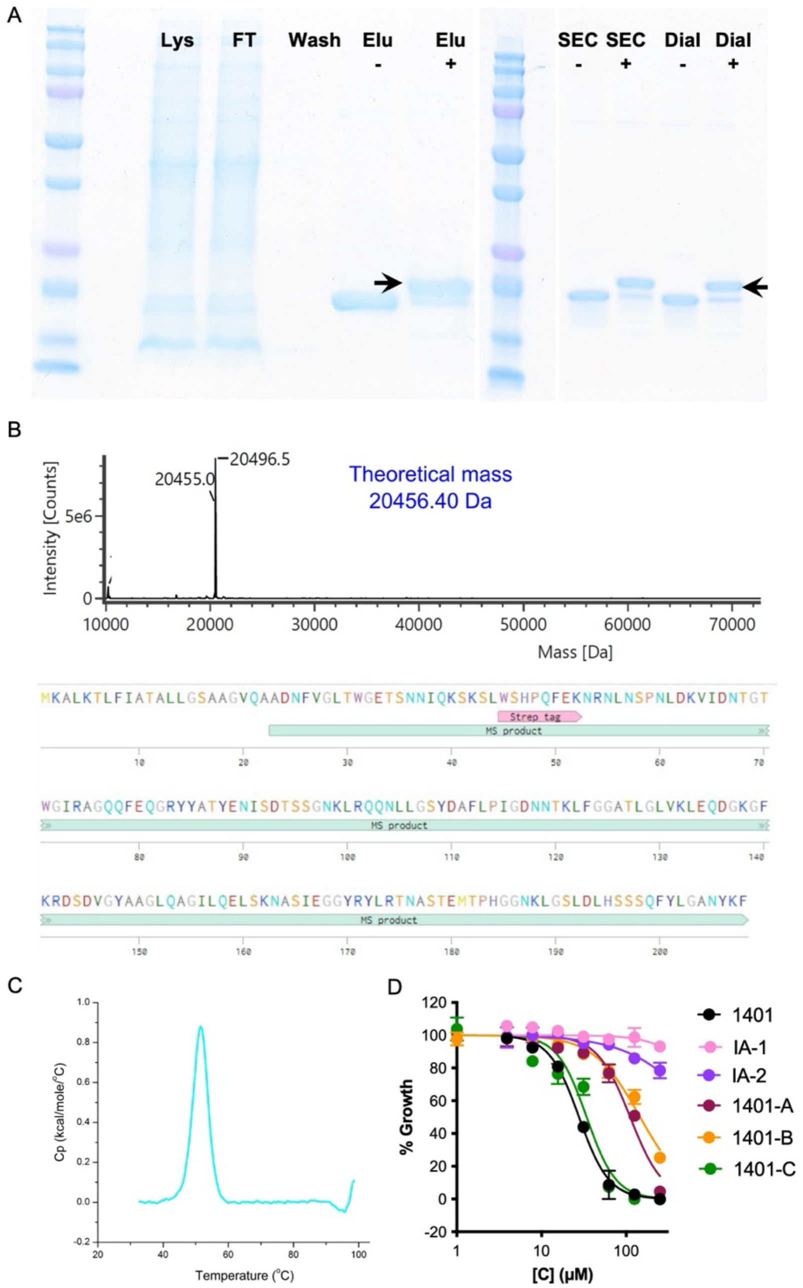


**Figure S9. A) B) and C) Expression and purification of StrepII-tagged OprH from *P. aeruginosa* and D) Activity of BRD1401 analogs.** A) SDS-PAGE gel of samples collected during purification of StrepII-OprH from *P. aeruginosa* (PA30 in Table S1G) using Streptactin beads is shown on the left; lysate (Lys), flow-through (FT), wash, elution samples with (Elu+) and without (Elu–) boiling. The eluted sample was further purified on a Size Exclusion Chromatography (SEC) column, relevant fractions pooled and then dialyzed to obtain pure protein. The SDS-PAGE gel of pooled SEC fraction with (SEC+) and without (SEC–) boiling and dialyzed protein with (Dial+) and without (Dial–) boiling is shown (see Method Details). The upward shift in StrepII-OprH protein mobility upon boiling indicates folded protein[18] and is highlighted with the black arrowhead. B) Intact mass spectrometry analysis of purified StrepII-OprH from *P. aeruginosa* and its amino acid sequence is shown. The predicted mass for a processed (cleaved signal peptide)

sequence is 20456.4 Da and is annotated as MS product in the amino acid sequence. The higher mass likely reflects an acetonitrile adduct. C) Differential Scanning Calorimetry trace of purified StrepII-OprH protein shows an unfolding transition peak at 51ºC. D) Activity of BRD1401 (1401) and 5 analogs (structures shown in figure 4A) towards the *oprL*-KD strain in media with 0.063% arabinose and a dose response of the tested compounds. Normalized growth, calculated in % relative to vehicle (0.5% DMSO) and positive controls, measured with an absorbance ($OD_{600nm}$)-based assay is plotted against compound concentration (n= 3, error bars indicate SD).

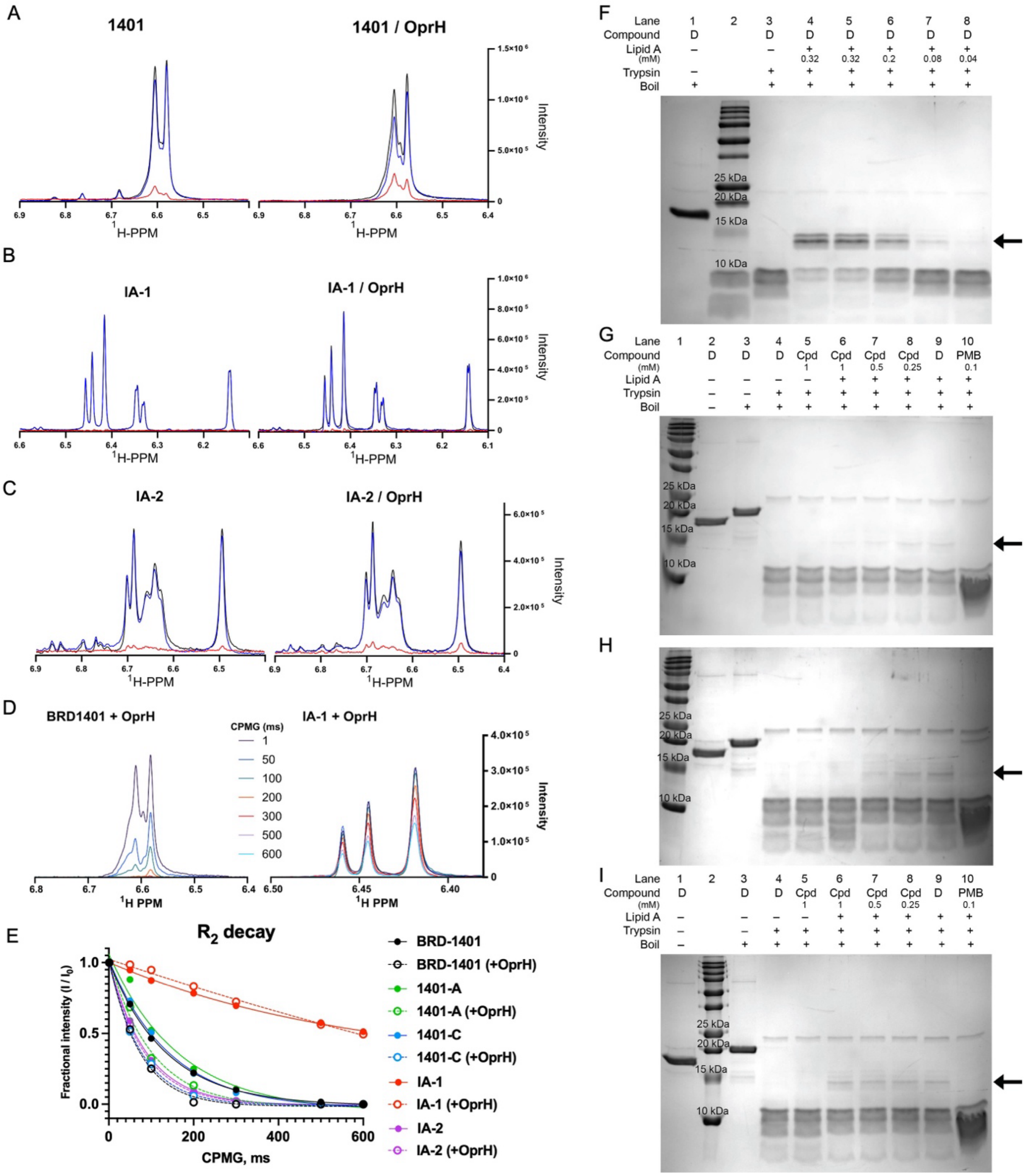


**Figure S10. BRD1401 binds OprH.** A), B) and C) STD-NMR of BRD1401 and two inactive analogs ($IC_{50}$ > 250 µM against *oprL*-KD strain; Figure S9D), IA-1 and IA-2, with purified *P. aeruginosa* OprH protein. Overlaid 1H-spectra of the aromatic region chemical shifts and intensities of 500 µM the compound solution in dialysis buffer (left) and in the presence of 15 µM *P. aeruginosa* OprH (right; See Method Details) without irradiation of OprH (black) and with irradiation of OprH (blue) and the difference spectrum (STD in red) A) BRD1401 and inactive analogs B) IA-1 and C) IA-2 with and without OprH. Enhancement of the STD spectrum in the presence of OprH suggests binding to OprH, which is clearly observed for BRD1401 and not for

IA-1 and IA-2. D) $^1$H-NMR $T_2$-CPMG spectra (x-axis in PPM) of BRD1401 and the inactive analog IA-1 with relaxation delays colored in navy (1ms) to turquoise (600ms). Attenuation of the compound signal (y-axis) with increasing delay times is a consequence of a small molecule binding to OprH. E) $R_2$ relaxation plots representing fractional intensities relative to 1ms spectra for BRD1401 and its analogs with (open circles) and without (closed circles) OprH. The rate is quantified by exponential decay fit for each compound and condition. Faster signal decay rates in the presence of protein suggest binding to OprH. Data extracted from 1 representative experiment as shown in panel D. F), G), H) and I) Impact of BRD1401 on *in vitro* binding of Kdo2-Lipid A to OprH. F) SDS-PAGE gel-based detection of a Lipid A-protected OprH protein fragment after incubation of 16 μM of OprH with (+) or without (–) a dose response of Kdo2-Lipid A and subsequently digested with 42 μM of trypsin (+) or not digested (–). Samples were boiled and run on an SDS-PAGE gel to detect the LPS-protected band at ~14.5 kDa, which is highlighted with the arrow. Impact of BRD1401 (B), the active analog, 1401-C (C), or the inactive analog, IA-1 (D) on Kdo2-Lipid A-mediated protection of OprH from trypsin digestion. Detection of the Lipid A-protected ~14.5 kDa sized OprH protein fragment (arrow) after incubation of OprH with (+) or without (–) 80 μM of Kdo2-Lipid A and a dose response of each of the compounds and subsequently digested with trypsin (+) or not digested (–). Samples were boiled (+) or not (-) and run on an SDS-PAGE gel. Representative gel from 2 independent experiments is shown. The intensity of the 14.5 kDa band is reduced at the higher concentrations of BRD1401 and 1401-C indicating disruption of the OprH-Lipid A interaction by these compounds while IA-1 has no effect. Cpd refers to tested compound, D refers to DMSO vehicle control and PMB refers to polymyxin B, the positive control.

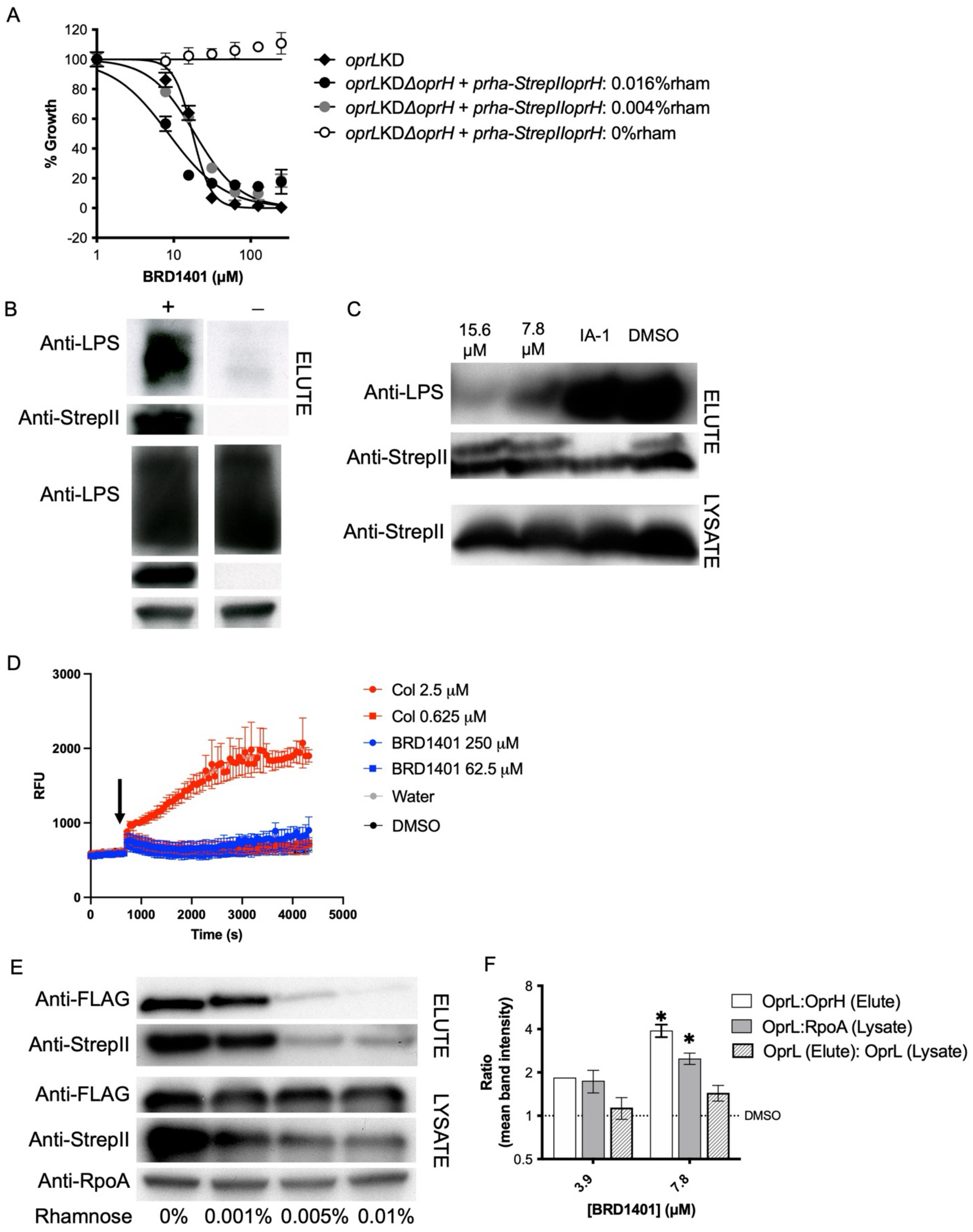


**Figure S11.** A) BRD1401 activity towards the *oprL*-KD or *oprL*-KDΔ*oprH/pRha-StrepII-oprH* (PA32 in Table S1G) strains in media with 0.063% arabinose that downregulates OprL expression. Rhamnose (rham) was added in varying amounts to drive episomal expression of *strepII-oprH* in

PA32. Normalized growth, calculated in %, measured from an absorbance ($OD_{600nm}$)-based growth assay is plotted against BRD1401 concentration (n= 3, error bars indicate SD). Over-expression of StrepII-OprH in the *oprL*-KDΔ*oprH* strain conferred BRD1401sensitization. B) LPS is pulled down by StrepII-tagged OprH from *P. aeruginosa* lysate. LPS and StrepII-OprH were detected by western blotting either after cell lysis (LYSATE) or after pull-down from lysates with Streptactin beads (ELUTE), using anti-LPS(O10) and anti-StrepII antibodies, respectively. Representative blot of samples from *oprL*-KDΔ*oprH/pRha-StrepII-oprH* (+; PA32 in Table S1G) and the control *oprL*-KDΔ*oprH/pRha-oprH* (–; PA31 in Table S1G) is shown. OprL expression was induced with 0.5% arabinose while StrepII-OprH or untagged OprH was driven with 0.13% rhamnose. Anti-RpoA antibody was used as a loading control in the lysate samples. C) BRD1401 reduced LPS binding to OprH but did not affect StrepII-OprH protein levels. Representative western blot of elution samples after pull-down of lysates (top) or the lysate samples directly (bottom) prepared from PA32 strain after exposure to the vehicle control (0.5% DMSO), BRD1401 at 7.8 or 15.6 μM or the inactive analog, IA-1, at 62.5 μM is shown. The strain was grown in media with 0.063% arabinose, which downregulates OprL, while StrepII-OprH was induced with 0.13% rhamnose. Antibodies are identical to (B). D) Ethidium bromide uptake assay to monitor OM permeabilization after compound exposure. *oprL*-hypomorph strain was equilibrated with 5 μM of ethidium bromide in phosphate buffered saline and then exposed (indicated by arrow) to 2 doses of colistin (Col) or BRD1401for the indicated time. Relative fluorescence units (RFU; $\lambda_{em}$= 585nm and $\lambda_{ex}$= 535nm), a direct measure of ethidium bromide uptake and binding to DNA, is plotted against time (n=3; error bars indicate SD). Colistin treatment led to significant permeabilization at the high concentration while BRD1401 had no effect. E) OprL binding to OprH is specific. Western blot detection of OprL-FLAG and StrepII-OprH in the *oprL-FLAG-KDStrepII-oprH/pRha-oprH* (PA39 in Table S1G) strain after cell lysis (LYSATE) and after incubation of lysate with Streptactin beads followed by elution (ELUTE) with excess biotin (One representative blot is shown). The strain was exposed to varying doses of rhamnose in the medium to drive the expression of untagged OprH, while OprL-FLAG expression was induced with 0.5% arabinose and StrepII-OprH expression was driven by the native promoter. OprL-FLAG was detected with an anti-FLAG antibody while the anti-StrepII and anti-RpoA antibodies are identical to (B). StrepII-OprH protein band intensity is reduced at higher rhamnose doses indicating transcriptional or post-translational regulation of StrepII-OprH because of over-expression of untagged OprH. Hence, ratio of OprL-FLAG to StrepII-OprH band intensities in the elution samples was calculated to measure specificity of this interaction. F) Impact of BRD1401 treatment on co-immunoprecipitation of OprL-FLAG with StrepII-OprH in the *oprL-FLAG*-KDΔ*oprH/pRha-StrepII-oprH* strain (PA36 in Table S1G). The strain was grown to log phase with indicated concentrations of BRD1401 and 0.063% arabinose and then lysed for pull-down using Streptactin beads. Western blotting was performed with antibodies described in (E) and protein band intensity was quantified with ImageJ. Ratio of the mean intensity of the OprL-FLAG band relative to the StrepII-OprH band in elution samples or of the OprL-FLAG band relative to the RpoA loading control band in lysate or of normalized OprL-FLAG in elution versus lysate samples is plotted (normalized to the DMSO vehicle control; dashed line = 1) (n=3-4; error bars indicate SEM). * indicates $p$-value < 0.05 calculated using a One sample $t$-test in comparison to the vehicle control.

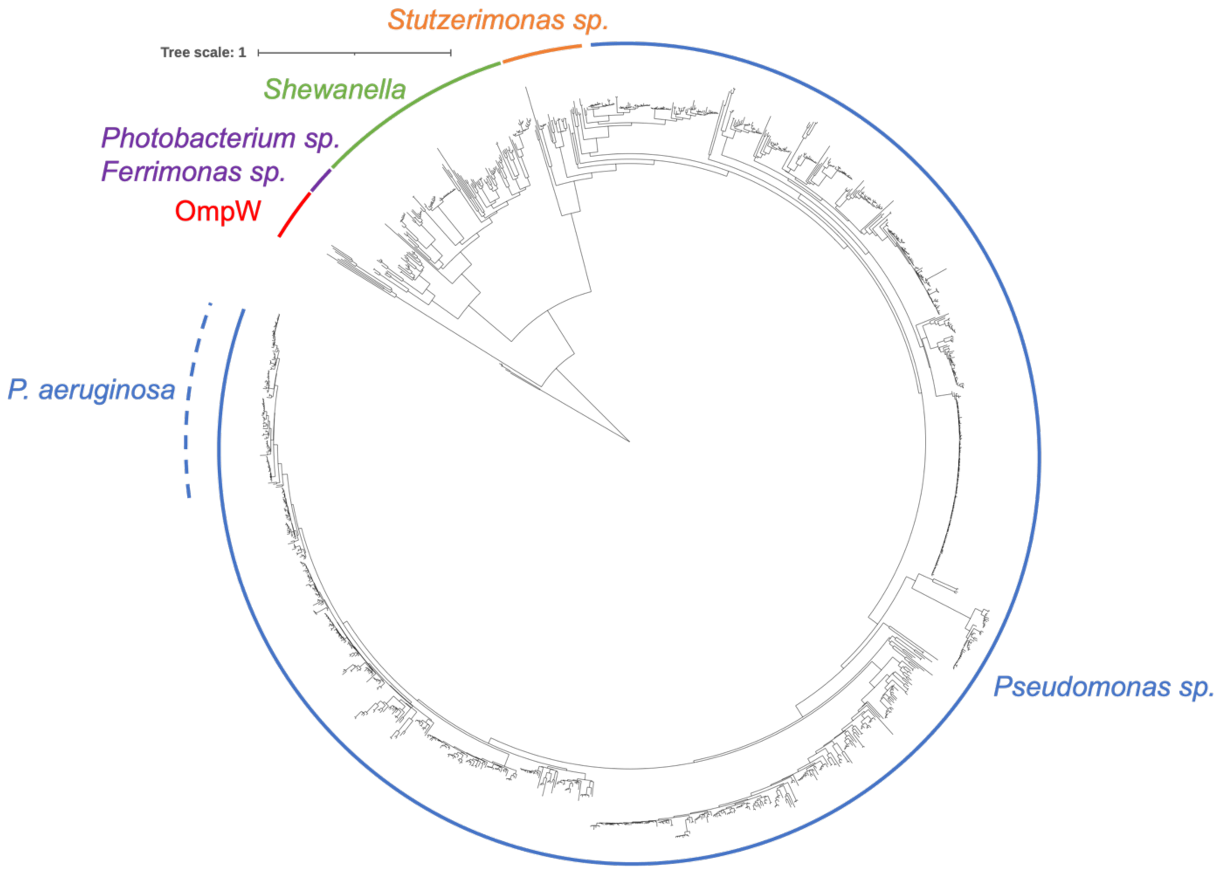


**Figure S12. Phylogenetic tree of OprH homologs.** The OprH sequence from PA14 (PA14_49200) was used as the query protein for a PSI-BLAST search and the resulting hit sequences were used to construct a phylogenetic tree as described in Method Details. The distant homolog OmpW (red) from *E. coli* and other bacterial species appear as hits, albeit with only ~14% identity. *Pseudomonas* species (blue) was the predominant genus in the search including *P. aeruginosa* (blue, dashed), *P. putida*, and *P. fluorescens*, among others. *Stutzerimonas sp.* (orange), *Shewanella* (green), and *Photobacterium sp.* and *Ferrimonas sp.* (purple) comprised the remaining species.

**Table S2. Multiplexed screen analysis**

**Table S2A.** Table of Z' factors for each individual strain included in the pilot multiplexed screen

| Species | Name | Z'-factor |
|---|---|---|
| *S. aureus* | Newman | 0.22* |
| *E. coli* | CFT073 | 0.464 |
| *K. pneumoniae* | MGH48 | 0.642 |
| *A. baumannii* | ATCC19606 | 0.443 |
| *P. aeruginosa* | PA14 | 0.611 |
| *P. aeruginosa* | *bamA*-hypomorph | 0.681 |
| *P. aeruginosa* | *bamE*-hypomorph | 0.578 |
| *P. aeruginosa* | *folA*-hypomorph | 0.553 |
| *P. aeruginosa* | *folP*-hypomorph | 0.552 |
| *P. aeruginosa* | *gcp*-hypomorph | 0.549 |
| *P. aeruginosa* | *gyrA*-hypomorph | 0.594 |
| *P. aeruginosa* | *leuS*-hypomorph | 0.613 |
| *P. aeruginosa* | *lolA*-hypomorph | 0.630 |
| *P. aeruginosa* | *lolB*-hypomorph | 0.598 |
| *P. aeruginosa* | *lppL*-hypomorph | 0.604 |
| *P. aeruginosa* | *lptA*-hypomorph | 0.632 |
| *P. aeruginosa* | *lptD*-hypomorph | 0.205 |
| *P. aeruginosa* | *lpxC*-hypomorph | 0.647 |
| *P. aeruginosa* | *mreC*-hypomorph | 0.567 |
| *P. aeruginosa* | *murA*-hypomorph | 0.554 |
| *P. aeruginosa* | *oprL*-hypomorph | 0.687 |
| *P. aeruginosa* | *tolB*-hypomorph | 0.638 |

*The Z'-factor was calculated from a separate pilot experiment. Since the focus of the screen was gram-negative pathogens there was no optimization of screening conditions to improve the Z'-factor for *S. aureus*.

**Table S2B.** Table listing the number of screening hits (SLF< –3) and single strain hits (compounds with SLF less than –3 only against that one specific strain) identified from the ~54k compound library.

| Species | Strain/Gene | Subcellular Location | Screening Hits* | Single strain hits |
|---|---|---|---|---|
| *E. coli* | CFT073 | | 568 | 90 |
| *S. aureus* | RB120 | | 5149 | 4381[#] |
| *K. pneumoniae* | MGH48 | | 24 | 2 |
| *A. baumannii* | ATCC19606 | | 227 | 135 |
| *P. aeruginosa* | PA14 | | 2 | NA |
| *P. aeruginosa* | *bamA*-hypomorph | Outer membrane | 6 | NA |
| *P. aeruginosa* | *bamE*-hypomorph | Outer membrane | 17 | 4 |

| | | | | |
|---|---|---|---|---|
| *P. aeruginosa* | *folA*-hypomorph | Cytoplasm | 1 | NA |
| *P. aeruginosa* | *folP*-hypomorph | Cytoplasm | 1 | 1 |
| *P. aeruginosa* | *gcp*-hypomorph | Cytoplasm | 4 | 1 |
| *P. aeruginosa* | *gyrA*-hypomorph | Cytoplasm | 9 | 1 |
| *P. aeruginosa* | *leuS*-hypomorph | Cytoplasm | 30 | 5 |
| *P. aeruginosa* | *lolA*-hypomorph | Periplasm | 5 | 1 |
| *P. aeruginosa* | *lolB*-hypomorph | Outer membrane | 3 | 1 |
| *P. aeruginosa* | *lppL*-hypomorph | Outer membrane | 8 | NA |
| *P. aeruginosa* | *lptA*-hypomorph | Periplasm | 7 | NA |
| *P. aeruginosa* | *lptD*-hypomorph | Outer membrane | 235 | 16 |
| *P. aeruginosa* | *lpxC*-hypomorph | Cytoplasm | 11 | 2 |
| *P. aeruginosa* | *mreC*-hypomorph | Periplasm | 2 | 1 |
| *P. aeruginosa* | *murA*-hypomorph | Cytoplasm | 3 | NA |
| *P. aeruginosa* | *oprL*-hypomorph | Outer membrane | 381 | 60 |
| *P. aeruginosa* | *tolB*-hypomorph | Periplasm | 93 | 9 |

*These numbers do not include the two pan-active compounds, since those were deemed to be active across all species by a separate metric.
[#]The low Z'-factor for *S. aureus* in pilot experiment (Table S2) precluded confidence in hits called for this strain.

**Table S2C.** Demultiplexed confirmation rates for compound-strain interactions with SLF < -3.

| | >30% growth inhibition | >50% growth inhibition | >70% growth inhibition |
|---|---|---|---|
| % Compounds | 39% (49 of 126) | 21% (27 of 126) | 10% (13 of 126) |
| % Compound-strain combinations | 37% (59 of 160) | 23% (36 of 160) | 10% (16 of 160) |
| % Compounds with indicated growth inhibition only in strains with SLF < -3, and not for strains with SLF > -3 | 12% (6 of 49) | 52% (14 of 27) | 92% (12 of 13) |

**Table S2D.** Demultiplexed confirmation rates for compound-strain interactions with SLF < -6.

| | >30% growth inhibition | >50% growth inhibition | >70% growth inhibition |
|---|---|---|---|
| % Compounds | 72% (21 of 29) | 41% (12 of 29) | 31% (9 of 29) |
| % Compound-strain combinations | 68% (25 of 37) | 43% (16 of 37) | 27% (10 of 37) |
| % Compounds with indicated growth inhibition only in strains with SLF < -6, and not for strains with SLF > -6 | 19% (4 of 21) | 42% (5 of 12) | 100% (9 of 9) |
